# A multilayered in silico analysis links UHRF1, DNA methylation and developmental chromatin memory to lineage-dependent prognosis in gastric, renal and adrenal cancers

**DOI:** 10.64898/2026.08.14.742686

**Authors:** Jacopo Biotti, Livio Muccillo, Filippo Macchi, Marco Spadarotto, Chiara Gino, Marco Finocchiaro, Elena Magnani, Simona Corso, Cristina Migliore, Daniela Conticelli, Simone Serio, Roberto Papait, Federica Donnarumma, Pellegrino Mazzone, Francesco Albano, Vittorio Colantuoni, Mariangela Tamburello, Gianluigi Mazzoccoli, Tommaso Colangelo, Tiziana Alberio, Geppino Falco, Sandra Sigala, Silvia Giordano, Mauro Fasano, Daniela Furlan, Ian Marc Bonapace

## Abstract

Aberrant DNA methylation is a hallmark of cancer, but its clinical interpretation remains debated. UHRF1, a key epigenetic adaptor for DNA methylation maintenance and chromatin bivalency regulation in embryonic stem cells, is frequently overexpressed yet shows context-dependent prognostic behaviour. By integrating bulk and single-cell transcriptomics, CpG-resolution methylation, developmental chromatin states, immune profiling and clinical outcomes across gastric (STAD), clear-cell renal (KIRC) and adrenal (ACC) carcinomas, we identified a four-class UHRF1-embryonic morphogenesis (UHRF1-EM) framework resolving this paradox. This axis revealed an inverse prognostic pattern: whilst across all three tumours EM-low and EM-high states mark better or worse prognosis, respectively, UHRF1-high levels associate with favourable outcome in STAD (UH-EML), and unfavourable in KIRC and ACC (UH-EMH). The classification proved reproducible and independently prognostic after adjustment for stage and molecular subtypes, outperforming existing classifiers and exceeding pathological stage in KIRC and ACC. Multivariable models incorporating UHRF1-EM yielded uniformly positive ΔC-indices.

Hypermethylation associated with the UHRF1-EM axis was enriched at ESC bivalent developmental loci (EM and oncofoetal genes), but not at housekeeping cell-cycle sites. In STAD, this pattern was related to oncofoetal gene downregulation and best prognosis, whereas in KIRC and ACC it matched with gene-body/enhancer methylation, higher EM expression, immunosuppressive microenvironments and worst prognosis.

Together, these findings establish the UHRF1-EM axis as a clinically robust molecular classifier and support a mechanistic model in which tumour-specific epigenetic engagement of developmental loci may contribute to the prognostic inversion, providing a foundation for further mechanistic experimental validation.

## Introduction

DNA methylation plays a crucial role in regulating gene expression in mammals and is extensively restructured in cancer genomes. Aberrant methylation patterns aid tumour classification and prognostic stratification, but understanding the underlying mechanisms remains a challenge. Whilst promoter hypermethylation can silence tumour suppressor genes, changes in non-canonical methylation at enhancers, gene bodies and repetitive elements frequently reflect deeper alterations in cellular identity, developmental plasticity and chromatin organisation that profoundly shape tumour behaviour^1–5^. Understanding how specific epigenetic regulators link methylation dynamics to clinically significant tumour states is therefore both a scientific and a translational priority.

UHRF1 (ubiquitin-like with PHD and RING finger domains 1) is a key player in this regulatory environment. It functions as an essential scaffolding hub within several interacting multi-protein assemblies, each of which coordinates a distinct dimension of epigenetic regulation. It identifies hemi-methylated CpGs through its SRA domain and specific histone marks via its PHD (non-methylated H3R2) and TTD (H3K9me2/3) domains and mono-ubiquitinates H3K18, thus serving as both a recruiter and activator of DNMT1 onto replicating chromatin^6^. In concert with DNMT1 and the chromatin remodeller HELLS, UHRF1 forms a complex that couples ATP-dependent nucleosome repositioning to maintenance methylation, providing the mechanical access to otherwise occluded silencer regions that passive replication-fork recruitment alone cannot achieve^7,8^. All this is essential for maintaining DNA methylation across cell divisions^6,9–11^, as evidenced by studies showing that Uhrf1-KO mice cannot sustain global CpG methylation and die around day 9 in development^9^, highlighting its critical role in epigenetic inheritance from the earliest stages of mammalian development. Disruption of the UHRF1–DNMT1 interface^12,13^, therefore, induces defects across both histone and DNA methylation layers^14^.

Beyond this primary function, UHRF1 also coordinates DNMT3- and TET-linked methylation stability, maintains heterochromatin, ensures genomic stability, suppresses transposable elements and modulates immune and tumour suppressor programmes^1,10,11,15,16^. Collectively, these roles position UHRF1 not merely as a passive maintenance factor but as an interpreter of chromatin context, capable of stabilising or altering methylation states.

UHRF1 is overexpressed across multiple tumour types^2,17–19^. Experimental disruption of the integrity of UHRF1 complexes, whether through UHRF1 depletion or through direct targeting of the DNMT1– UHRF1 or EHMT2–UHRF1 interfaces, reduces proliferation and tumorigenic features in gastric, renal, hepatic and lung cancers^12,13,14^, although reactivation of methylation-silenced genes is frequently incomplete, owing to residual HDAC-dependent repression within the same complex^20,21^. Accordingly, UHRF1 has emerged as a therapeutic target^13,22,23^. Nevertheless, the consequences of UHRF1 perturbation vary markedly across cancer lineages: in some contexts, UHRF1 activity sustains malignant features through canonical methylation-based silencing, whilst in others it shapes cell state or immune interactions through non-canonical mechanisms that are independent of cell proliferation^15,21,24,25^. These lineage-specific differences suggest that the composition and activity of UHRF1 are fundamentally conditioned by the chromatin landscape of each tissue of origin.

A unifying explanation for this context dependency may reside in developmental epigenetic memory. In embryonic stem cells, developmental regulator genes occupy bivalent chromatin domains co-marked by activating H3K4me3 and repressive H3K27me3, maintaining them in a transcriptionally poised state until lineage commitment^5^. These bivalent loci are disproportionately susceptible to aberrant hypermethylation in cancer, and the embryonic chromatin state at such regions can predict the methylation landscape of future tumours years in advance^26–28^. Crucially, the COMPASS complex—acting through UHRF1 and Setd1a—regulates H3K4me3 establishment at bivalent domains in pluripotent cells, supporting orderly lineage specification^29^. Conversely, the ECReM complex, by coupling H3K9 methylation to DNA methylation maintenance, is well placed to convert bivalent poising into stable heritable repression when developmental signalling is aberrantly reactivated^12,14^. This raises the possibility that tumour methylomes retain a molecular imprint of the embryonic complex activity specific to their tissue of origin, and that the dominant UHRF1-associated complex in a given lineage determines whether aberrant methylation enforces stable silencing or instead allows the permissive, oncofoetal transcriptional programmes associated with tumour plasticity, invasion and treatment resistance^4^.

In this study, we investigate this question across three cancer types—gastric adenocarcinoma (STAD), clear-cell renal cell carcinoma (KIRC) and adrenocortical carcinoma (ACC)—selected for their markedly different prognostic correlations with UHRF1 expression and their distinct developmental origins. By integrating patient transcriptomes, CpG-resolution DNA methylation profiles, embryonic stem cell chromatin annotations, tissue development data, single-cell and immune-stromal analyses, and long-term clinical outcomes derived from TCGA and complementary public datasets, we explore three related hypotheses: whether UHRF1-associated methylation alterations preferentially target developmental regulatory pathways; whether they reflect the bivalent chromatin architecture established in embryonic stem cells; and whether they account for the divergent patient outcomes observed across these lineages.

## Results

### Context-dependent prognostic impact of UHRF1 and embryonic morphogenesis programmes across human tumours

Given UHRF1’s controversial role in oncogenesis and metastasis, pan-cancer transcriptome analysis stratifying TCGA tumours by type-specific median UHRF1 expression revealed two opposing prognostic patterns (Figure 1A).

**Figure 1.**
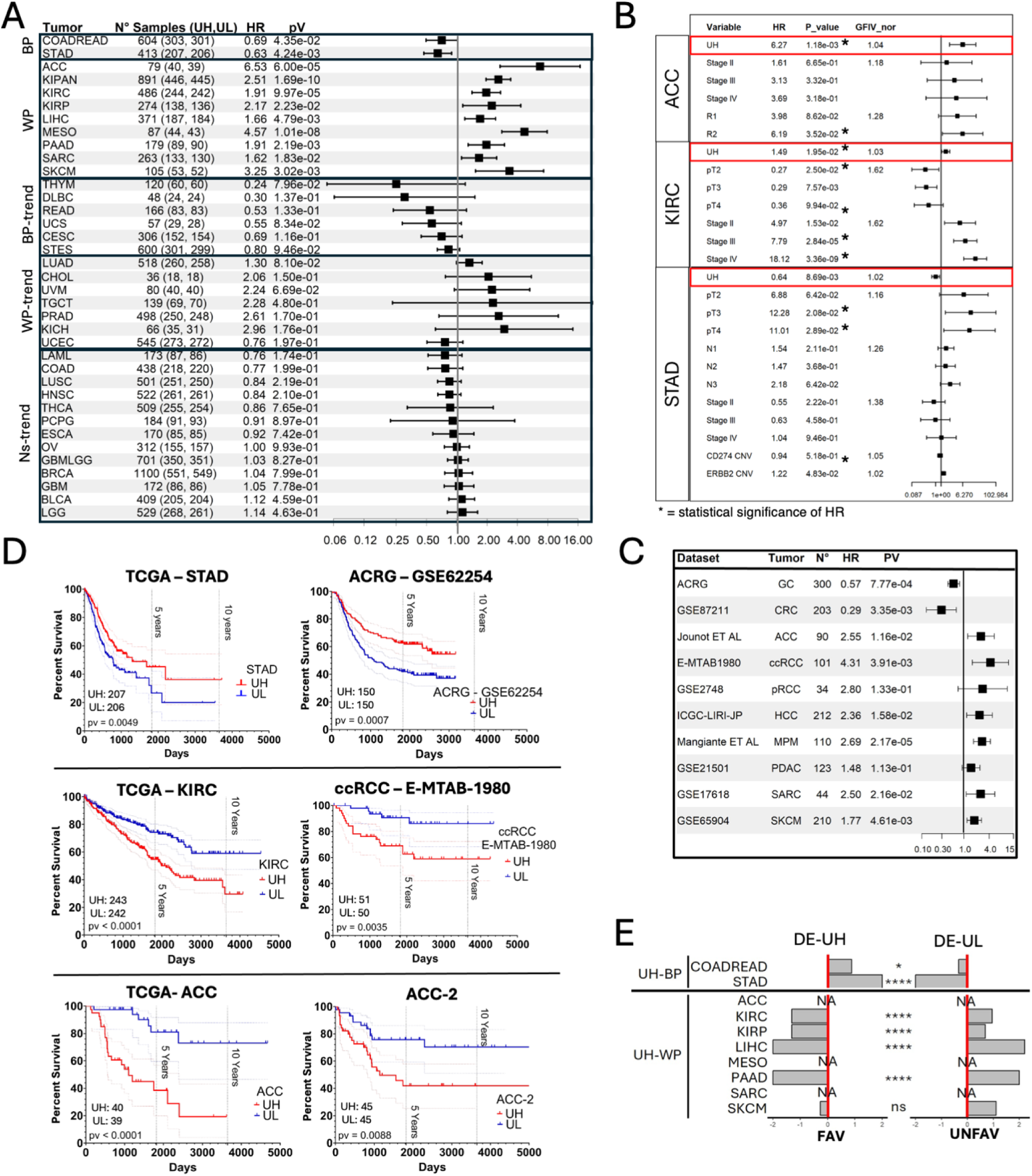
UHRF1 has an independent prognostic role in TCGA tumours. (A) Hazard ratio (HR) from survival analysis, comparing high (UH) with low (UL) UHRF1 expression in TCGA dataset. Horizontal lines stand for 95% confidence intervals for the ratios; (B) Hazard Ratio (HR) and Generalized Variance Inflation Factor (GVIF) from multivariate Cox Regression analysis of UH against UL classes after adjustment for other clinical covariates in TCGA-ACC, KIRC and ACC (pathological T (pT), N (pN), stage, residual tumour (R), copy number variations (CNV) of CD274 and ERBB2); (C) Hazard ratio (HR) from survival analysis, comparing UH with UL patients in second datasets of UHRF1-best and –worst prognosis tumours. Univariate Cox proportional hazards model was used for HR calculations; (D) overall survival of UH and UL patients in stomach (TCGA-STAD, ACRG-*GSE62254*), kidney (TCGA-KIRC, ccRCC – BioStudies *E-MTAB-1980*) and adrenocortical (TCGA-ACC, ACC-2GSE19750) cancer; UHRF1-*high* (red), UHRF1-*low* (blue) (log-rank *Mantel-Cox* p value); (E) Log scale ratio (-2;2) between favourable (FAV) genes in UH/UL and unfavourable (UNFAV) genes in UH/UL in UHRF1-*significative* tumours (*Fisher’s test* p value) defined by The Human Protein Atlas (see M&M for details).

UHRF1-high associated with better prognosis (UH-BP) in stomach adenocarcinoma (STAD) and colorectal cancer (COADREAD), but with poorer outcome (UH-WP) in hepatocellular (LIHC), papillary and clear cell renal (KIRP, KIRC), adrenocortical (ACC), skin melanoma (SKCM), pancreatic (PAAD), mesothelioma (MESO) and sarcoma (SARC). The trend was largely independent of pathological features, with staging dependency in kidney and adrenocortical cohorts (Table 1; Figure 1B; Figure S1B), and was confirmed in external validation datasets (Figure 1C). Other tumour types showed weak or no association (Figure 1A, forest plot). Kaplan-Meier plots for the ten most significant TCGA cancers are shown in Figure 1D and Figure S1.

**Table 1:**
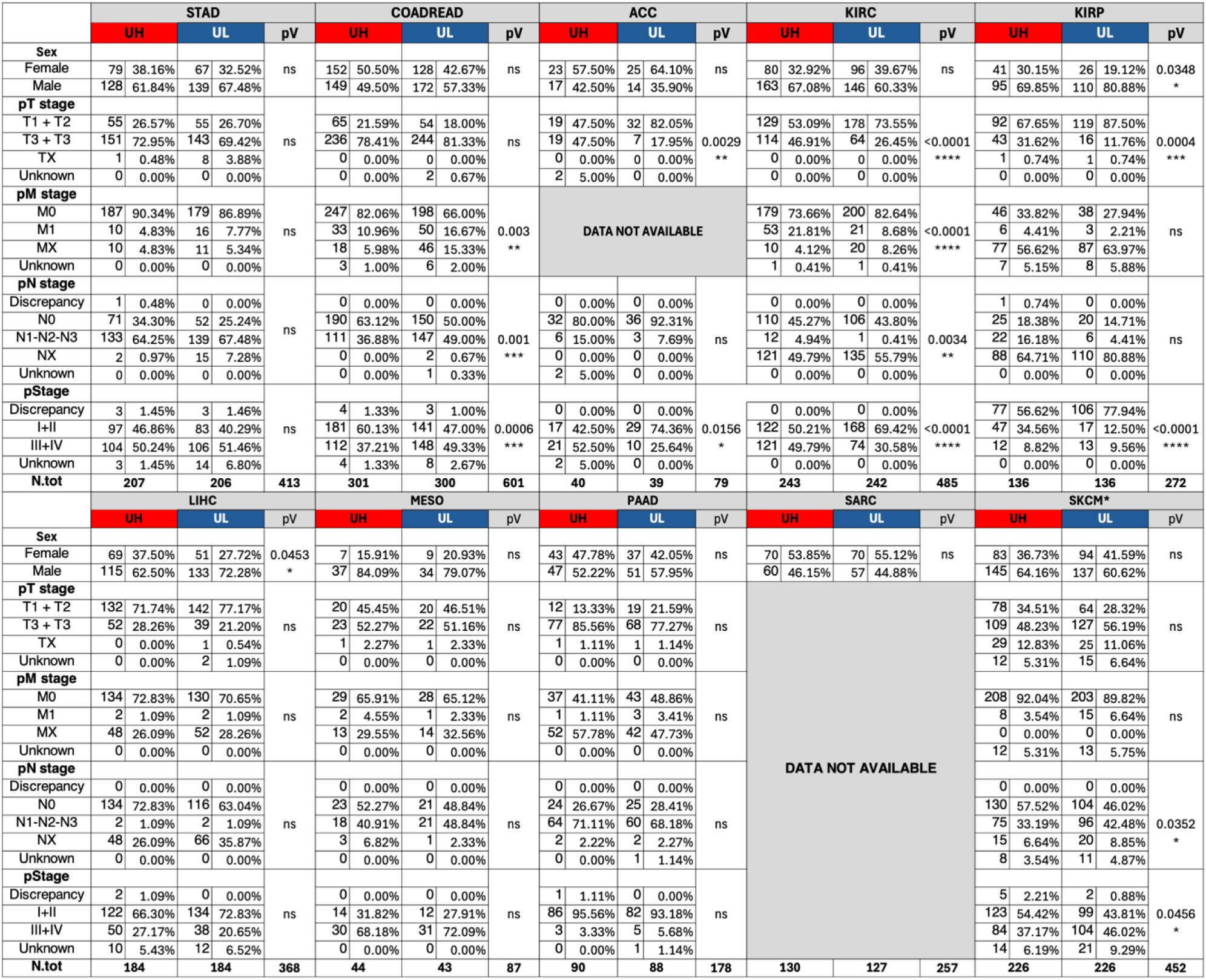
Distribution of UH and UL patients across known pathological and clinical classes. pV: Chi-Square or Fisher Test p Value. Ns: not significant. pT = pathological T stage, pN = pathological N stage, pM = pathological M stage, p stage = pathologcal stage. *= p value < 0.05, **= p value <0.01, ***= p value < 0.001, ****=p value < 0.0001. pT

Differential expression analyses between UH and UL tumours (DE-UH, DE-UL; FC≥1.4, log2FC≥0.485; Mann-Whitney U-test; Additional file 1), integrated with Human Protein Atlas^30^ enrichment of favourable (FAV) and unfavourable (UNFAV) prognostic genes, revealed an inversion of prognostic signatures (Figure 1E): in UH-BP tumours, FAV genes were upregulated in DE-UH and UNFAV in DE-UL; in UH-WP tumours the pattern was reversed.

### Embryonic morphogenesis GO category untangles UHRF1’s opposite prognostic impact

GO pathway analysis confirmed association of DE-UH with DNA repair, chromatin organisation and cell cycle, but highlighted Embryonic Morphogenesis (EM; GO:0048598) as the only term consistently mirroring the inversion of UHRF1’s prognostic effect (Figure 2A, Figure S2; Additional file 2).

**Figure 2.**
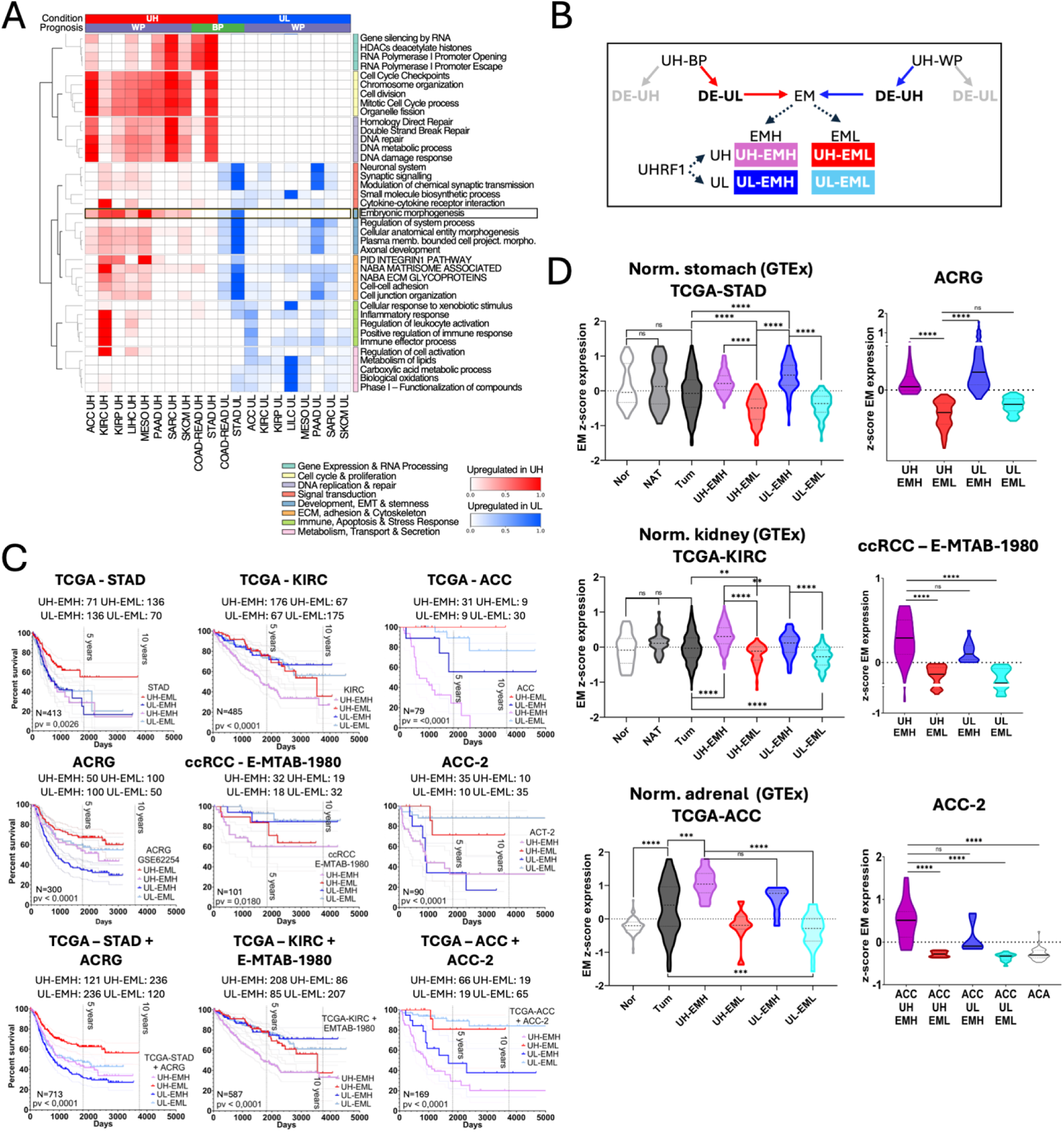
Combined UHRF1 and embryonic morphogenesis expression enable transcriptional, prognostic and clinical stratification. (A) Selection of the top 5 enriched pathways in differentially expressed DE-UH and DE-UL genes based on a custom ranking metric which integrates maximum absolute significance (Max |-LogP|) and inter-sample variance. columns: tumour-specific DE-UH gene list (Red); tumour-specific DE-UL gene lists (Blue). Prognosis: BP (green): best prognosis in UHRF1-high condition; WP (purple): worst prognosis in UHRF1-high; rows: Metascape categories; (B) Definition of the four UHRF1-EM categories: EM genes are from UHRF1 DE-UL genes of the UH-BP tumours, and UHRF1 DE-UH genes of the UH-WP tumours; (C) Differences in overall survival within TCGA-STAD, KIRC, ACC, ACRG, E-MTAB-1980 and ACC-2, and aggregated databases, dependent upon UHRF1-EM categories. UHRF1-high EM-high (violet), UHRF1-high EM-low (RED), UHRF1-low EM-high (BLUE), UHRF1-low EM-low (light blue) (log-rank Mantel-Cox p value); D) Violin plots displaying differences in Embryonic Morphogenesis (GO:0048598) expression across normal adult tissue (Nor), normal adult tissue adjacent tumour (NAT) and UHRF1-EM conditions in TCGA-STAD, KIRC, ACC, ACRG, E-MTAB-1980 and ACC-2. **** = p value<0.0001; *** = p value<0.001; **= p value<0.01; * = p value<0.05; ns= not significant (Kruskal Wallis and Dunn test p value).

EM genes encompass foetal-like reprogramming, stemness, EMT/metastasis and therapy resistance. In UH-BP tumours, EM signatures were downregulated in UH patients; in UH-WP tumours they were co-regulated, suggesting EM upregulation is the primary driver of poor prognosis across lineages. Combining UHRF1 and EM into four groups (Figure 2B) further stratified outcomes: in STAD (UH-BP), UH-EML and UL-EMH defined best and worst outcomes, respectively, while in UH-WP tumours like KIRC and ACC this was reversed, with UH-EMH showing worst prognosis and UL-EML the best (Figure 2C-D; Figure S1B, Figure S3; Additional file 3).

### STAD, KIRC and ACC as model systems for dissecting context-dependent UHRF1 prognostic functions

STAD (UH-BP), KIRC and ACC (UH-WP) were selected for further dissection. UHRF1 expression significantly influenced survival up to 10 years (Figure 1D; Table 1; Additional file 4), confirmed in validation datasets and after restricting to tumour purity ≥60% (Figure 1C, Figure S1C).

Multivariate Cox models (Figure 1B) including pStage, pT, pN, resection status and, in STAD, CD274/ERBB2 copy number changes^31^, confirmed UHRF1 as an independent prognostic factor with low multicollinearity: favourable in STAD (HR=0.64, P=8.69×10⁻³) but adverse in ACC (HR=6.27, P=1.18×10⁻³) and KIRC (HR=1.49, P=1.95×10⁻²). EM levels followed the prognostic hierarchy of UHRF1-EM subgroups in discovery and validation cohorts (Figure 2C-D), supporting these three tumours as models for context-dependent UHRF1 functions.

Across all three tumours, UHRF1 consistently coupled to proliferation GO categories, while its relationship to oncofoetal and microenvironment categories was context-dependent (Figures S4 and S5; Additional File 2).

### UHRF1-EM classes stratify prognosis independently of mutational and molecular subtypes

Bar charts (Figure 3A) and multilayer heatmaps annotated by molecular subtype, stage, survival, recurrent mutations and mutation load (Figure 3B) exhibited robust expression patterns, with best/worst prognosis categories preserved as in Figure 2D. Somatic alteration recurrence was assessed across UHRF1-EM categories, focusing on non-silent mutations in key drivers (TP53, PIK3CA, MLH1, CDH1 in STAD; VHL, PBRM1, SETD2 in KIRC; TP53 in ACC) and total mutation burden (TCGA calls^31–33^; Figure 3B; Additional file 5).

**Figure 3.**
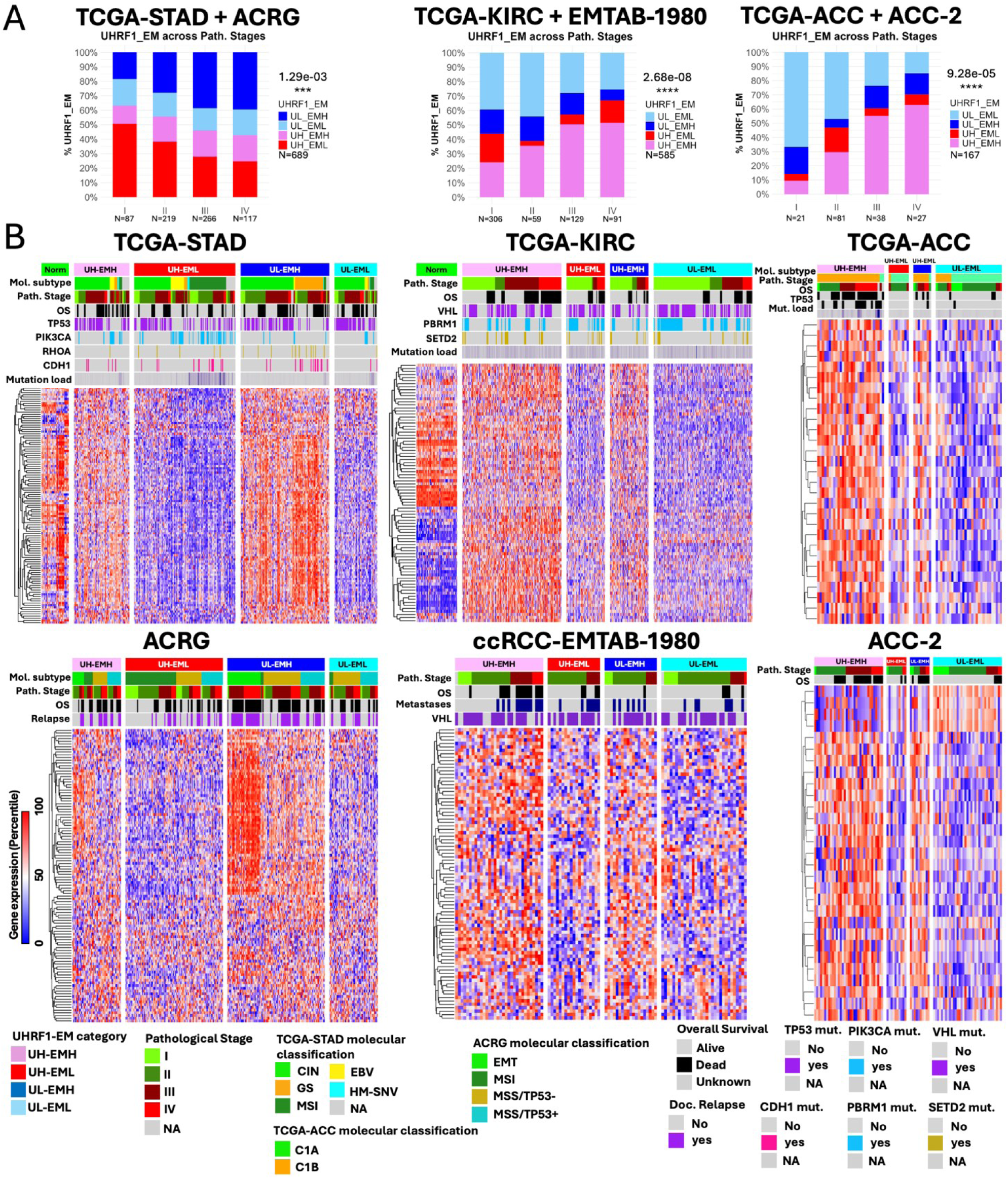
Clinical and molecular features coherently distribute across UHRF1-EM prognostic categories. (A) Bar chart showing proportion of UHRF1-EM cases across pathological stages in the three cumulative cohorts of gastric, kidney clear cell and adrenocortical carcinoma (Chi-square p-value); (B) Heatmap of mRNA expression profile (represented as percentiles) of Embryonic Morphogenesis genes across UHRF1-EM categories in TCGA-STAD (DE-UL), KIRC and ACC (both DE-UH) and a second database for each tumour: ACRG for stomach, E-MTAB-1980 for kidney, ACC-2 for adrenocortical cancer. UHRF1-EM categories, TCGA molecular subtypes (Mol. subtype), tumour pathological stage (Path. Stage), vital status (OS), relevant mutations and mutation load are indicated above TCGA-STAD, KIRC, ACC heatmap. UHRF1-EM categories, ACRG Mol. subtypes, Path. Stage, OS and documented disease recurrency are indicated above ACRG heatmap; UHRF1-EM categories, Path. Stages, OS are indicated above ccRCC-EMTAB-1980 heatmap; UHRF1-EM categories, Path. Stage, OS are indicated above the ACC heatmap.

In STAD, UH-EML (favourable) displayed higher mutation burden than other categories, including UL-EMH (p<0.0001). TP53 mutations were similar in UH-EML and UL-EMH (∼42%), while PIK3CA mutations were enriched in UH-EML (31.62% vs 4.48%; p<0.0001), as were MLH1 mutations (Additional file 5A-B). This non-linear pattern suggested high mutation burden in UH-EML reflected MSI-driven, immune-active biology rather than aggressiveness^34^. In KIRC, no significant difference in mutation burden was observed between prognostic categories. VHL and PBRM1 mutations were enriched in UL-EML (p=0.0023 and p<0.0001), and rare TP53 mutations peaked in UH-EMH at 5% (p=0.05; Additional file 5A, 5C). In ACC, EM expression matched prognostic ranking and UH-EMH showed the highest mutation burden (p<0.0001), with TP53 mutations concentrated in this poor-prognosis class (p<0.003). None of these variations reproduced the survival separation of UHRF1-EM classes (Figure S6).

We next examined whether UHRF1-EM status refined established molecular classifications. In TCGA-STAD and ACRG, ∼70% of hypermethylated MSI/EBV patients with best prognosis fell into UH-EML. Poor-prognosis categories were dominated by UL-EMH: 80% of GS and 35% of CIN tumours in TCGA-STAD, and 75% of EMT tumours in ACRG, with UH-EML under-represented (TCGA p=7.07×10⁻²¹; ACRG p=1.30×10⁻²¹). MSI tumours showed lowest EM and best outcome, whereas GS/CIN and EMT subtypes displayed high EM and poor survival. In ACC, UH-EMH comprised 65% of the very-high-risk C1A subgroup, while only 15% of UL-EML fell within C1A; UL-EML were enriched in C1B (65%; p=1.84×10⁻⁸). Thus, UHRF1-EM classification sharpened, and in some settings outperformed, existing subtype systems (Figure 3B; Figure S7A, S7C, S8A, S8C). Multivariate Cox regression showed the four UHRF1-EM categories retained independent prognostic significance after adjustment for clinical variables and molecular subtypes (Figure S9), defining a reprogramming axis largely independent of stage, subtype or mutational profile (Figure 3A-B; Figure S7A, 8A).

### UHRF1-EM categories define an embryonic–proliferative continuum underlying relapse risk

Long-term clinical features differed across datasets. In TCGA-STAD, UH-EML cases more often achieved complete response or stable disease than UL-EMH, though not statistically significant (Chi-Square t-test); in ACRG, recurrence was lower in UH-EML and higher in UL-EMH (27% vs 58%, p=0.0003; Figure 3B; Figure S8A). In TCGA-ACC, UH-EMH was enriched for progressive disease after primary therapy (38.71% vs 6.67% in UL-EML) and UL-EML for complete remission (83.33% vs 29.03%, p=0.0022) (Figure S7A). Mechanistically, relapse genes were enriched for developmental/oncofoetal programmes and non-relapse-associated genes for cell-cycle, proliferation and DNA repair (Figure S10A). Scoring ACRG and STAD patients by mean expression of relapse-associated embryonic genes versus non-relapse-associated proliferative genes showed UHRF1-EM categories position tumours along an embryonic–proliferative continuum underlying relapse risk (Figure S10B).

To refine this framework, tumour-specific oncofoetal (OnF) signatures^4^ were combined with UHRF1 expression using the same UHRF1-EM criteria. In STAD, most favourable outcomes occurred in subgroups with lowest OnF expression, whereas in KIRC and ACC the poorest prognosis showed the highest OnF score (all p<0.0001; Figure S11A–C; additional files 2, 7; Figure S11B). Oncofoetal GO programmes associated with adverse outcomes-transport, angiogenesis, WNT signalling and stemness-mirrored EM patterns and displayed opposite correlations with UHRF1 in STAD versus KIRC/ACC, pointing to context-dependent UHRF1 regulation and shared transcriptional logic underlying EM and OnF programmes.

### Tumour-specific UHRF1-centred epigenetic factors oppositely correlate with embryonic and oncofoetal programmes across cancer types

Correlation analysis for putative epigenetic factors acting with UHRF1 identified 15 genes positively correlated (Epi-Co) and 4 inversely correlated (Epi-Anti) (Figure 4A; Figure S12A). Many Epi-Co genes are established UHRF1 partners or components of shared repressive machineries, including the core DNA methylation module (UHRF1–DNMT1–SUV39H1/2–CBX3) and PRC1/2 complexes (CBX7/CBX8, EZH1/EZH2)^6,8,12,18,35^. In STAD, Epi-Co genes correlated negatively with EM/OnF expression while Epi-Anti correlated positively; the reverse occurred in KIRC and ACC (Figure 4A; Figure S12A).

**Figure 4.**
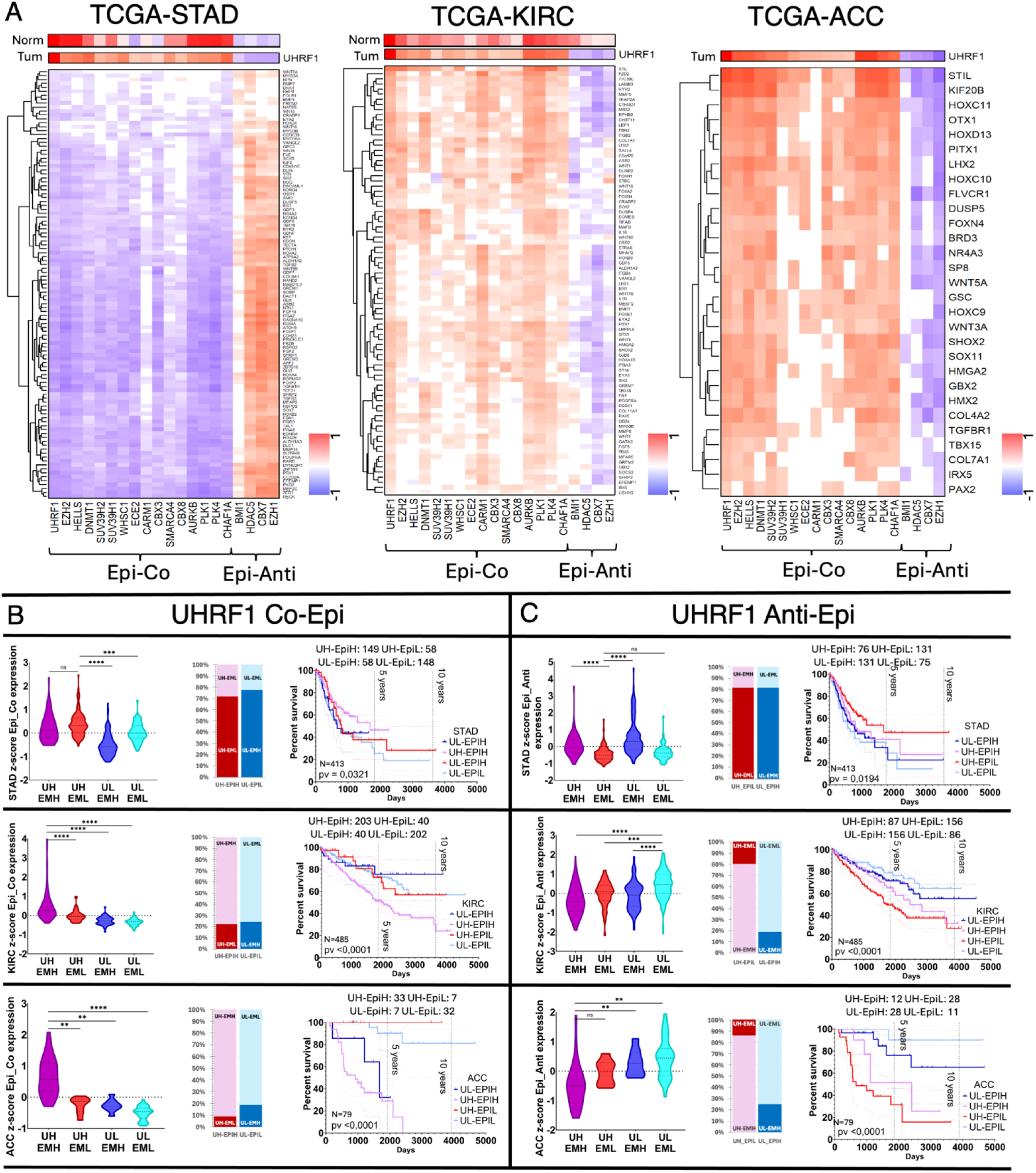
Tumour-specific UHRF1-centred epigenetic factors oppositely correlate with embryonic and oncofoetal programmes across cancer types. (A) heatmap reporting the Spearman coefficient of the correlation between epigenetic effectors (columns) and embryonic morphogenesis genes (rows). The coefficient varies from -1 (blue, anti-correlation) to 1 (red, correlation); (B) Violin plot indicating the z-scored expression of UHRF1 co-related epigenetic effectors across UHRF1-EM categories in TCGA-STAD (upper panel), KIRC (intermediate panel), ACC (lower panel) (Kruskal-Wallis and Dunn’s test); bar plot indicating the relative distribution of UHRF1-EPI patients (x-axis) across UHRF1-EM categories in TCGA-STAD, KIRC and ACC; Differences in patients’ overall survival within TCGA-STAD, KIRC, dependent upon the four UHRF1-EPI categories (log-rank Mantel-Cox p value); (C) Violin plot indicating the z-scored expression of UHRF1 anti-corelated epigenetic effectors among the four UHRF1-EM categories in TCGA-STAD (upper panel), KIRC (intermediate panel), ACC (lower panel); bar plot indicating the relative distribution of UHRF1-EPI patients (x-axis) across UHRF1-EM categories in TCGA-STAD, KIRC and ACC; Differences in patients’ overall survival within TCGA-STAD, KIRC, dependent upon the four UHRF1-EPI categories.

Kaplan-Meier analyses sustained the EpiCo-UHRF1 correlation in all three tumours, confirming the central role of chromatin modifiers (Figure 4B). EZH2 closely tracked UHRF1 across tumour types: co-expressed, sharing 35–65% of differentially expressed targets, displaying the same tumour-specific prognostic directionality, and inversely correlated with EZH1 (Figure S12B, D, E). These results suggest UHRF1 influences EM/OnF programmes through tumour-specific epigenetic assemblies switching from repressive (STAD) to activating (KIRC/ACC) roles, with involvement of inversely correlated factors (EZH1, CBX7, HDAC5, BMI1).

### UHRF1-EM classes orchestrate tumour-specific immune-stromal ecosystems and oncofoetal microenvironments

Immune and stromal landscapes, characterized via multi-algorithm TIMER2.0 deconvolution, differed significantly across UHRF1-EM classes and broadly aligned with prognosis (Figure 5A). In STAD, UL-EMH displayed high infiltrates of endothelial cells, haematopoietic stem cells, CAFs, myeloid dendritic cells, monocytes and stromal scores, while UH-EML was enriched for NK, Tfh, CD8⁺, γδ and CD4⁺ Th2 cells; Tfh and γδ T cells associated with improved outcome, whereas high stromal score, haematopoietic stem cells and endothelial cells were linked to adverse prognosis^36^. In KIRC, CAFs, Tregs, Tfh and CD4⁺ Th2 cells were increased in UH-EMH, whereas UL-EML had higher endothelial cells, non-regulatory CD4⁺ T cells, neutrophils and activated mast cells; eosinophils, activated mast cells, haematopoietic stem cells, neutrophils, non-regulatory CD4⁺ T cells and endothelial cells correlated with favourable prognosis, while CAFs, plasmacytoid dendritic cells, Tregs and Tfh cells were detrimental^37^. In ACC, typically “immune cold”^33^, the worst-prognosis UH-EMH showed enrichment for activated myeloid dendritic cells, plasma B cells, CD4⁺ Th1 and Th2 cells, whilst UL-EML was infiltrated by favourable-outcome cytotypes. Survival analyses (Figure 4A; HR lanes) corroborated these patterns: in ACC, CD4⁺ effector memory T cells, myeloid dendritic cells, microenvironment score, activated mast cells and B cells were favourable (UL-EML), while CAFs, NK cells and CD4⁺ Th1/Th2 cells associated with worse survival (UH-EMH).

**Figure 5.**
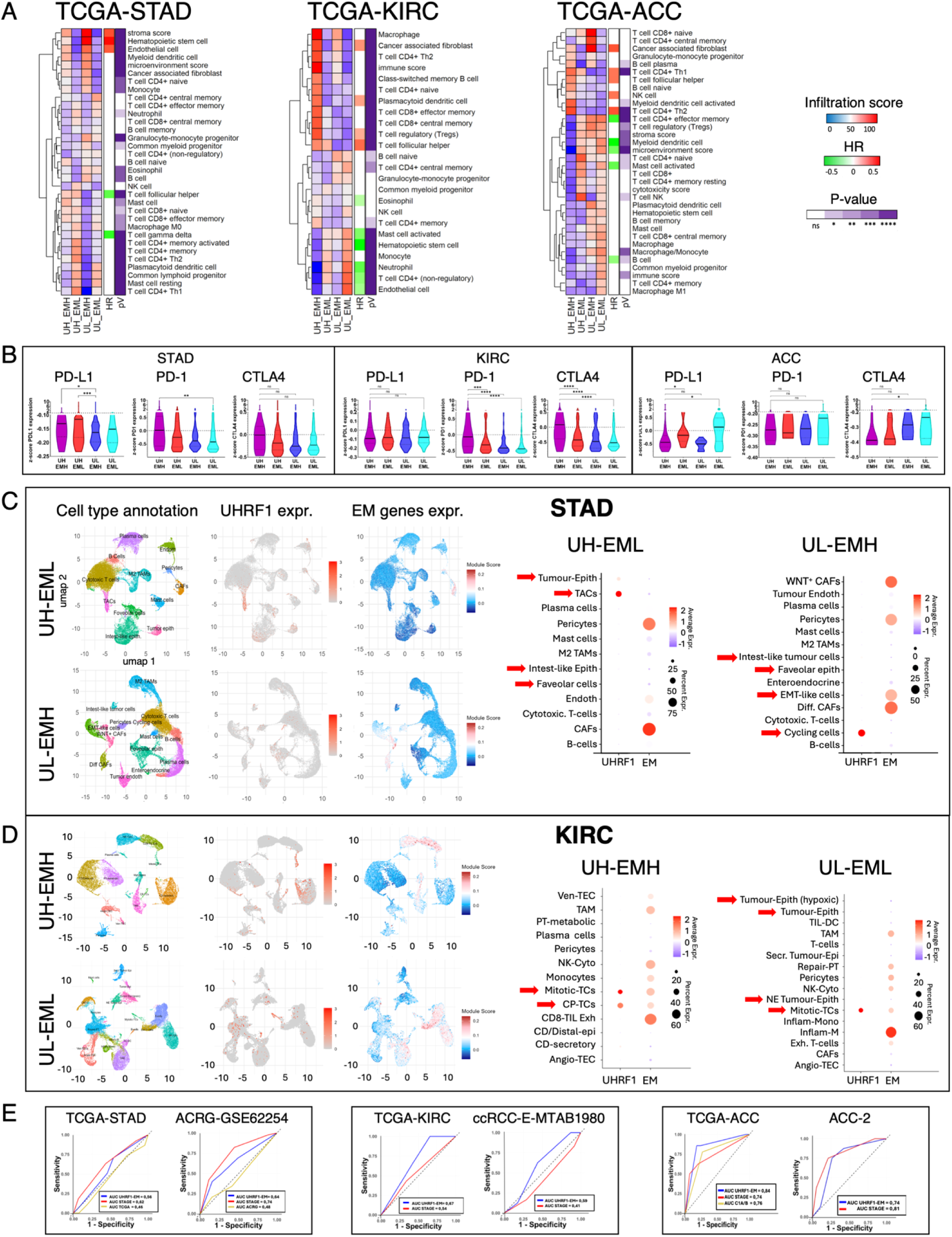
Immune infiltration prediction reveals a differential immune microenvironment across UHRF1-EM categories and single-cell expression of UHRF1 and EM genes in STAD and KIRC. A) Heatmap showing immuno-cellular infiltration scores (rows) predicted by the TIMER2.0 deconvolution algorithm across UHRF1-EM categories (columns). HR: hazard ratio calculated by comparing patients with infiltrate levels above and below the median for each tumour; pV: ANOVA and post hoc Dunn’s test p-value for multiple comparisons across UHRF1-EM categories. B) PD-1, PD-L1, and CTLA4 expression across UHRF1-EM categories in TCGA-STAD, KIRC, and ACC (Kruskal-Wallis and Dunn’s test p-values). C) ROC curves indicating the prediction performance of UHRF1-EM classes compared with traditional pathologic stages and other molecular subtypes, based on overall survival (1 = dead, 0 = alive) of patients with gastric, kidney, and adrenocortical carcinoma in TCGA and a second dataset (AUC = area under the curve). D) Single-cell analysis of UHRF1 and EM genes in gastric cancer (STAD). From left to right: UMAP representation of gastric cancer cell-type annotation, and UHRF1 and embryonic morphogenesis expression in UH-EML and UL-EMH patients; dot plot showing average and proportional expression of UHRF1 and EM genes (x-axis) across cell types (y-axis). E) Single-cell analysis of UHRF1 and EM genes in clear cell renal carcinoma (KIRC). From left to right: UMAP representation of cell-type annotation, and UHRF1 and embryonic morphogenesis expression in UH-EMH and UL-EML patients; dot plot showing average and proportional expression of UHRF1 and EM genes (x-axis) across cell types (y-axis).

Immune checkpoint expression mirrored these contexts. In STAD, PD-L1 was highest in UH-EML and lowest in UL-EMH, consistent with lymphocyte-rich MSI/EBV tumours. In KIRC, PD-1 and CTLA4 peaked in UH-EMH, reflecting a checkpoint-rich, immunosuppressive milieu. In ACC, UH-EMH showed modest reductions in PD-L1 and CTLA4, yet the UHRF1-EM axis still separated immune-active from immune-cold tumours (Figure 5B).

Single-cell RNA-seq data from gastric^38^ and renal carcinomas^39^ reclassified by UHRF1-EM revealed UHRF1 expression was largely confined to cycling transit-amplifying and malignant epithelial cells, whilst EM genes were preferentially expressed in stromal and immune compartments (Figure 5C-D). In STAD, UH-EML tumours showed increased cycling cells with EM activity restricted to pericytes and CAFs, whereas UL-EMH exhibited expansion of EM-positive WNT⁺ CAFs, differentiated CAFs and EMT-like cells-consistent with an oncofoetal microenvironment and invasive niches. In KIRC, UH-EMH tumours were enriched for cycling tumour cells co-expressing UHRF1 and EM genes alongside EM-positive NK cells, macrophages and monocytes, whereas UL-EML contained fewer EM-positive mitotic cells.

### UHRF1-EM classes refine or outperform stage-based and molecular classifiers in prognostic stratification

Receiver Operating Characteristic (ROC) analyses compared UHRF1-EM classes against pathological stage and established molecular classifiers (Figure 5E). In STAD, UHRF1-EM achieved an Area Under the Curve (AUC) of 0.56, exceeding TCGA molecular classes (0.46) but below stage (0.62). In ACRG, UHRF1-EM (AUC=0.64) again outperformed ACRG subtypes (0.48) but was less predictive than stage (0.74). Discrimination improved in kidney cancer, with UHRF1-EM AUCs of 0.67 (TCGA-KIRC) and 0.59 (E-MTAB-1980), both exceeding stage (0.54 and 0.41). In ACC, UHRF1-EM showed strong prognostic value (AUC=0.85), outperforming both stage (0.75) and the “Cluster of Clusters” classifier (0.81); in an independent ACC cohort, it remained highly predictive (AUC=0.74), only slightly below stage (0.81).

C-index analyses from single-predictor and multivariable Cox models provided complementary assessment (Suppl. Table 1). In STAD and ACRG, UHRF1-EM C-indices were 0.57 and 0.60, outperforming molecular subtyping (0.54 and 0.59) but inferior to stage (0.69 and 0.71). In KIRC, UHRF1-EM underperformed stage (0.60 vs 0.69 in TCGA-KIRC; 0.69 vs 0.77 in E-MTAB-1980), whereas in TCGA-ACC it achieved the highest C-index (0.82), exceeding stage (0.75) and C1A/C1B classification (0.74). Multivariable Cox models combining UHRF1-EM, molecular subtypes and stage consistently yielded higher C-indices than single predictors across all tumours, with UHRF1-EM contributing a positive ΔC-index, supporting its independent, additive prognostic value.

Overall, UHRF1-EM classes provided superior prognostic stratification by capturing the context-specific intersection of oncofoetal programmes and tumour proliferation. Whilst UHRF1 supported proliferation across all three tumours, its epigenetic coupling to EM/oncofoetal circuits diverged markedly: acting as a favourable repressive mark in STAD, whilst driving immunosuppression and adverse outcomes in KIRC and ACC.

### Tumour-specific UHRF1-linked CpG methylation architectures oppositely regulate embryonic and oncofoetal programmes and stratify prognosis

Given UHRF1’s key role in maintaining DNA methylation, we asked whether its context-dependent prognostic effects were encoded at the methylome. Ranking promoter- and gene-body CpGs by correlation of methylation with EM expression (EM expr/5meC) and with UHRF1 expression (UHRF1 expr/5meC) identified four recurrent clusters (A-D) with distinct epigenetic and clinical profiles (Fig. 6A-C; Table 2; Additional file 8).

**Figure 6.**
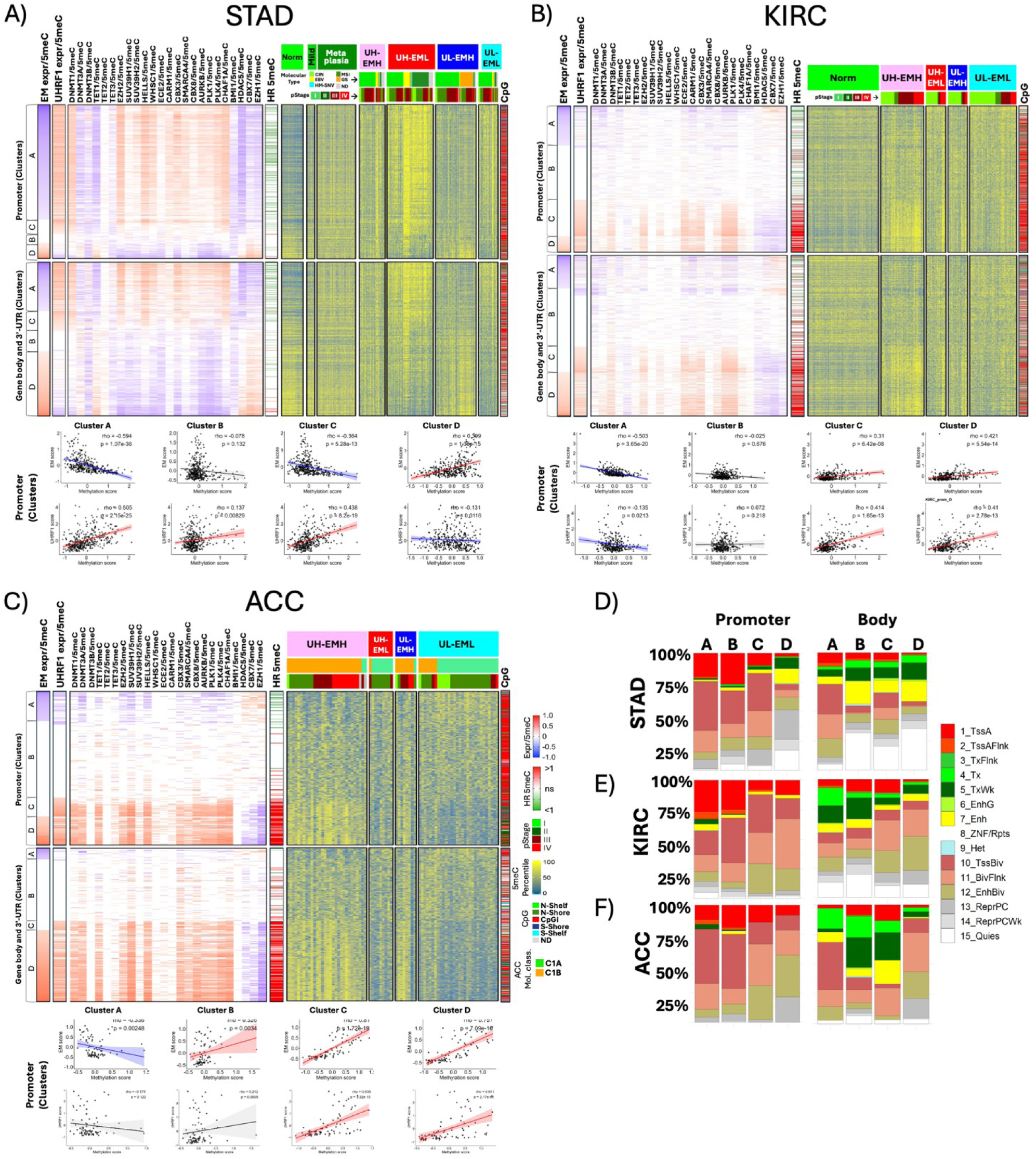
UHRF1-associated DNA methylation at bivalent sites regulates EM gene expression and influences prognosis. Heatmap of EM CpG methylation of across the four UHRF1-EM categories, normal and metaplastic tissue - when available - in (A) TCGA-STAD, (B) TCGA-KIRC, (C) TCGA-ACC. DNA methylation varies according to a yellow (high methylation) to blue (low methylation) gradient. UHRF1-EM categories, TCGA molecular subtypes and tumour pathological stage are indicated above the heatmap; rows: cytosines; columns: patients; EM expr UX-EMX: expression of the gene associated to the correspondent CpG; CpG: genomic context of the reported cytosines (blue: s-shore, light blue: s-shelf, red: island, light green: n-shelf, dark green: n-shore, gray: non-CpG); HR 5meC: hazard ratio for high vs low methylation level of the cytosine (green: HR<1, better prognosis, red: HR>1, worse prognosis); EM expression/5meC: correlation between EM gene expression and cytosine-specific methylation; UHRF1/5meC: correlation between the expression of the indicated epigenetic effector (UHRF1, DNMT1…) and CpGs’ methylation (Spearman’s correlation test, blue: anti-correlation; red: correlation; white: no statistically significant); Scatterplots A) TCGA-STAD, B) TCGA-KIRC, C) TCGA-ACC: correlation between 1) upper row: EM z-score expression (y-axis) and methylation z-score of CpG clusters (x-axis); 2) lower row: UHRF1 z-score expression (y-axis) and methylation z-score of CpG clusters (x-axis). Clusters of the promoter regions are shown, together with rho correlation coefficient. Colour code: red, blue and black regression line stand for statistically significant positive, negative, and non-significant correlation, respectively. 95% confidence interval were also overlaid. (D-E-F) Frequency distribution of cytosines across Epigenetic Roadmap genomic region annotations; X-axis: CpG clusters; y-axis: percentage of cytosines located in annotated genomic regions; Roadmap 15-state Chromatin Model abbreviations: 1_TssA: Active TSS, 2_TssAFlnk: Flanking Active TSS, 3_TxFlnk: 5’/3’ Transcription Flanking, 4_Tx: Strong Transcription, 5_TxWk: Weak Transcription, 6_EnhG: Genic Enhancer, 7_Enh: Enhancer, 8_ZNF/Rpts: ZNF Genes & Repeats, 9_Het: Heterochromatin, 10_TssBiv: Bivalent/Poised TSS, 11_BivFlnk: Bivalent Flanking, 12_EnhBiv: Bivalent Enhancer, 13_ReprPCWk: Repressed Polycomb; 14_ReprePCWk: Weak Repressed Polycomb; 15_Quiesc: Quiescent/Low signal.

**Table 2.** Definition and prognostic behaviour of EMexpr–5meC/UHRF1–5meC clusters.

| CLUSTER | EM Expr-5meC CORRELATION | UHRF1-5meC TREND* | METHYLATION PATTERNS ACROSS PROGNOSTIC GROUPS (Additional File 8) | PROGNOSTIC ASSOCIATION (HR-5meC in Figure 6A-C) |
| --- | --- | --- | --- | --- |
| A | Negative<br>(Canonical repressive methylation) | STAD: positive<br>KIRC/ACC: negative or week | Lowest methylation in poorest prognosis groups:<br>• UL-EMH in STAD<br>• UH-EMH in KIRC/ACC | Hypermethylation generally associated with HR < 1 |
| B | Weak or absent | No consistent trend | No distinct pattern relative to EM classes or survival | No consistent prognostic signal |
| C | None | Positive in all tumours | Highest methylation in:<br>• UH-EML in STAD<br>• UH-EMH in KIRC/ACC | Favourable in STAD; adverse in KIRC/ACC |
| D | Positive<br>(methylation linked activation) | STAD: negative;<br>KIRC/ACC: positive | Highest methylation in:<br>• UL-EMH in STAD<br>• UH-EMH in KIRC/ACC | Consistently associated with poor prognosis (HR > 1) |

Highest methylation in both promoters and gene bodies occurred at CpGs showing positive correlation between UHRF1 expression and EM-locus methylation, predominantly within CpG islands, and marked opposite prognostic extremes by lineage: the favourable UH-EML class in STAD (clusters A and C), and the adverse UH-EMH class in KIRC and ACC (clusters C-D). Crucially, the functional meaning of this hypermethylation diverged. In STAD, UHRF1-linked hypermethylation coincided with EM repression (cluster A; negative EM expr/5meC; cluster C with no significative EM expr/5meC correlation) and favourable outcome (HR<1; green lines in HR-5meC), consistent with canonical methylation-dependent silencing. In KIRC and ACC, hypermethylation associated with increased EM expression (cluster D; positive EM expr/5meC) and poor prognosis (HR>1; red lines in HR-5meC), indicating a non-canonical, transcriptionally permissive state (Fig. 6A-C). A partial non-canonical pattern was also detectable in poor-prognosis UL-EMH STAD tumours (cluster D), particularly at gene bodies, where hypermethylation showed negative correlation with UHRF1 but positive association with DNMT3A and a TET1-dominant signature. Cluster B in all three tumours comprises cytosines whose methylation shows no clear correlation with either EM or UHRF1 expression, although a subset still associates significantly with either favourable or adverse hazard ratios.

Distinct cofactor networks supported these states. In STAD clusters A and C, UHRF1-linked methylation coincided with positive DNMT1-5meC correlations together with co-effectors including EZH2, SUV39H1/2, HELLS, WHSC1, CBX3, SMARCA4, CBX8, CARM1, AURKB, PLK1, PLK4 and CHAF1A, and inverse coupling to EZH1, CBX7, HDAC5 and BMI1. In KIRC and ACC, the architecture shifted towards broader coupling with all three DNMTs and with TET enzymes (DNMT3B > DNMT3A > DNMT1 and TET3 > TET1 in KIRC; DNMT1, DNMT3A and TET1 > TET3 in ACC), consistent with, possibly, hydroxymethylation-linked and permissive chromatin. Despite this functional inversion, coupling to several co-effectors and inverse coupling to HDAC5, CBX7 and EZH1 were retained (Figure 6A; Supplementary tables 2–3).

Comparison with adjacent normal tissue confirmed disease specificity: STAD clusters A/C and KIRC clusters C/D showed tumour-acquired methylation, whereas STAD cluster D and KIRC cluster A were already hypermethylated in normal tissue. Oncofoetal genes showed concordant clustering and prognostic behaviour across all three tumour types (Figure S13). UHRF1-correlated hypermethylation therefore segregates prognostic extremes, but carries inverted biological meaning across lineages: repressive in gastric, permissive in renal and adrenocortical tumours, confirming its context-dependent oncogenic role.

### Single-gene UHRF1-linked methylation of developmental WNT/EM loci shows opposite prognostic roles in gastric versus renal and adrenal cancers

Single-gene CpG resolution confirmed tumour-specific prognostic associations with UHRF1 methylation (Figure S15). In STAD, EM genes linked to WNT signalling, development and oncogenesis (SFRP2^40^, DKK1^41^, GPC3^42^, TGFB2^43^) showed the greatest promoter/gene body hypermethylation, positive correlation with DNMT1 and EZH2, negative with EZH1 and UHRF1, and improved survival (TGFB2 HR:0.71; DKK1 HR:0.61; GPC3 HR:0.68; SFRP2 HR:0.68). Conversely, in KIRC and ACC, hypermethylation at EM genes including PITX1^44^, WNT16, SIX2^45^, GREM1^46^, HOXC9, HOXC10 and HOXD13^47^ was enriched in UH-EMH tumours and linked to poor outcome (KIRC: PITX1 HR:2.37, SIX2 HR:2.12, WNT16 HR:2.94, GREM1 HR:2.24; ACC: HOXC10 HR:4.66, HOXC11 HR:6.58, HOXD13 HR:3.33). Methylation correlated positively with UHRF1, EZH2 and DNMT3B/TET3 (KIRC) or DNMT3A/TET1 (ACC), remained negatively associated with EZH1, and tracked with EM expression. Unchanged-methylation CpGs lacked prognostic value, underscoring UHRF1 as a tumour-specific prognostic marker.

### Cell-cycle genes decouple UHRF1-linked methylation from expression, highlighting specificity for bivalent EM/OnF programmes

To test whether UHRF1-EM methylation dynamics were specific to EM/oncofoetal developmentally regulated genes, we analysed CpGs of cell-cycle (CC) genes (Reactome R-HSA 1640170), a hallmark strongly associated with cancer and UHRF1^1,48^. CC genes were consistently enriched among DE-UH genes across all three tumours, with substantial overlap (Additional file 2). However, their CpGs showed remarkably homogeneous methylation across tissues and UHRF1-EM categories, with higher methylation in gene bodies than promoters (Figure S14A, 14E, 14I). CC gene expression varied strongly: in KIRC and ACC it mirrored EM behaviour (lowest in UL-EML, highest in UH-EMH, poor prognosis) (Figure S14G-H, S14K-L), whereas in STAD it peaked in UH-EML, was lowest in UL-EMH and linked to better outcome (Figure S14C-D). Thus, DNA methylation is not a major determinant of CC expression, and the prognostic and methylation plasticity signals appear specific to tissue-restricted EM/OnF programmes.

### Developmentally primed EM/OnF categories and ESC bivalency shape tumour-specific, UHRF1-dependent methylation and prognostic behaviour in stomach and kidney

Integration of UHRF1-centred methylation with transcriptional and survival data suggested EM and OnF genes belong to conserved developmental categories repurposed in cancer. In STAD, KIRC and ACC, UHRF1 correlation with CpG hypermethylation of EM/OnF loci stratified molecular and prognostic groups (Figure 6; Figure S13; Suppl. Tables 2–3), with repressive methylation in gastric and activating methylation in renal/adrenocortical cancers. These genes include core canonical and non-canonical WNT components and other morphogenetic pathways supporting stemness and glandular morphogenesis in stomach versus differentiation, homeostasis and zonation in kidney and adrenal cortex^49^ (Figure S15).

Since EM/OnF categories are intrinsically developmental, reanalysis of ontogeny RNA-seq data from normal mouse stomach and kidney, and human normal adrenal gland defined developmentally regulated (DEV) gene sets^50–52^ (Figure 7).

**Figure 7.**
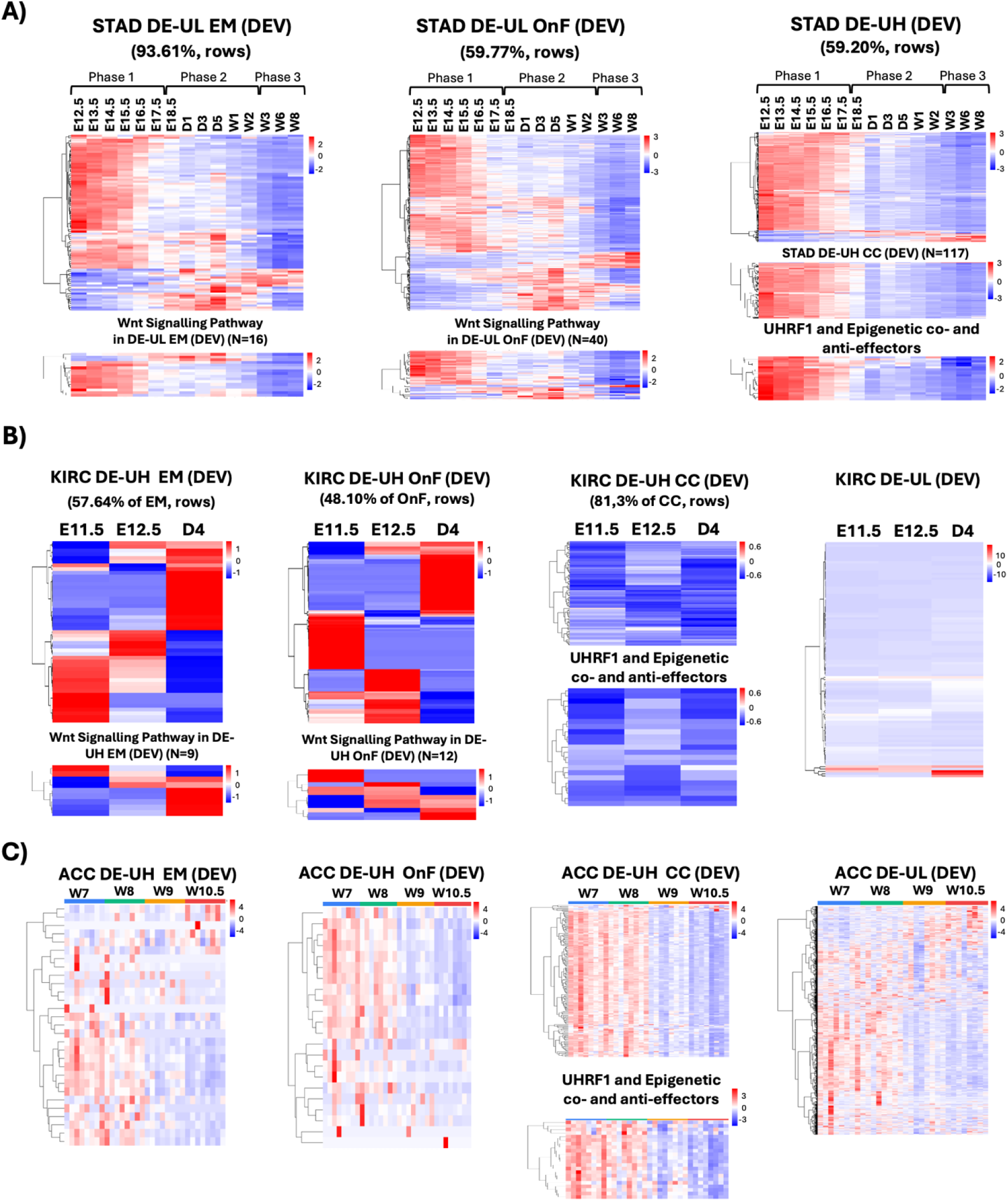
Embryonic Morphogenesis, Oncofoetal and proliferation genes are temporally regulated in mouse stomach and kidney and human adrenal development. A) Stomach heatmap clustering analysis of Embryonic Morphogenesis (EM), Onco-Foetal (OnF), Cell Cycle (CC) and epigenetic effectors DEV genes (rows) throughout developmental stages (phase I, II, III in columns; E: embryonic day; D: post-natal day; W: post-natal week); B) Kidney heatmap clustering analysis of DE-UH EM, DE-UH OnF, DE-UH CC and DE-UL developmental (DEV) genes (E: embryonic day; D: post-natal day 4); C) Adrenal gland heatmap clustering analysis of DE-UH EM, DE-UH OnF and DE-UH CC developmental (DEV) genes (W: post conception week).

Consistent with a developmental origin for EM/OnF reactivation, a large fraction of UHRF1-associated DE genes overlapped DEV: in STAD (DE-UL), 93.61% of EM and 59.77% of OnF genes; in KIRC (DE-UH), 57.64% of EM and 48.10% of OnF genes (Additional file 9). In stomach, DEV genes overlapping the STAD DE-UL set-including multiple WNT-pathway genes (56 mediators, including Wnt2b/6/9a/16, Sfrp1/2, Rspo2/3) and development/cancer regulators (Barx1, Gpc3, Tgfb2, Gli1/3, Sox7, Foxf1, Hoxa2/a4/d4)-peaked during early embryogenesis (Phase 1, Figure 7A) and were progressively silenced thereafter (Phases 2–3), consistent with preventing inappropriate intestinal priming^53^ (Figure 7A, STAD DE-UL EM and OnF). A smaller subset showed delayed/increasing expression, consistent with postnatal homeostatic functions. DEV CC genes (enriched in STAD DE-UH) largely followed the early-peaking trajectory, linking the UHRF1-stratified proliferative programme to embryonic transcriptional states. A similar trend was observed for several UHRF1 co- and anti-effectors (Figure 7A).

In KIRC DE-UH, DEV/EM-OnF overlaps recapitulated major nephrogenesis phases (Figure 7C)^49,54,55^. Canonical oncofoetal regulators (SIX2, EYA1, SALL4) clustered in early metanephric mesenchyme phases (Figure 7B, E11.5; Additional file 9), maintaining progenitor pools. ccRCC WNT components (WNT4, SFRP2, LHX1, LEF1) peaked during later epithelial maturation (Figure 7B, D4 stage; Figure 7C, Wnt Signalling Pathway, DE-UH EM and OnF), consistent with nephron formation/segmentation^56^.

In developing adrenal cortex, UHRF1 and epigenetic machinery declined progressively after early stages (post-conception weeks 7–8) during organogenesis (Figure 7C, ACC DE-UH co- and anti-epigenetic effectors), alongside oncofoetal genes (PHF19, TGFBR1; ACC DE-UH OnF, Figure S13 - ACC) and embryonic morphogenesis regulators (HOXC9; WNT5A, essential for adrenal patterning/zonation^57^; PITX1, whose hypermethylation predicts unfavourable outcome; Figure 6C). These signatures attenuate alongside proliferative programmes as tissue differentiates.

Altogether, tumour EM/OnF programmes largely represent redeployment of developmentally regulated modules, consistent with prior developmental analyses.

### UHRF1 rewires ESC-bivalent EM/OnF loci into tumour-specific methylation plasticity and prognostic programmes, while leaving housekeeping cell-cycle genes methylation unchanged

To mechanistically connect CpG clusters to chromatin states, we mapped EM (Figure 6A–C; clusters A–D) and CC CpGs (Figure S14A, S14E, S14I; clusters A–C) onto the 15-state ChromHMM annotation of H1 human ESCs (H1-ESC) using their genomic coordinates. This reference partitions the genome into 15 recurrent chromatin states inferred by a Markovian model, assigning each segment to a putative functional class^58^.

This framework distinguished actively transcribed regions (TssA), bivalent/poised regulatory elements (TssBiv, BivFlnk, EnhBiv; activating H3K4me3 or repressive H3K27me3) and repressed/heterochromatic domains, mapping selected methylated CpGs to cell-type–specific chromatin context and regulatory potential^58^ (EM: Figure 6D–F; CC: Figure S14B, S14F, S14J). EM genes showed a tight association between ESC bivalency and tumour-associated hypermethylation at promoters and gene bodies: in STAD this involved Clusters A and C (hypermethylated in UH-EML, BP), whereas in KIRC and ACC it involved Clusters C and D (hypermethylated in UH-EMH, WP). These CpGs were enriched for bivalent states (TssBiv, BivFlnk, EnhBiv), with KIRC and ACC shifting toward EnhBiv-dominated regions, consistent with non-canonical activation of developmental genes from hypermethylated bivalent enhancers^59^ (Figure 6D-F). By contrast, CC CpGs mapped predominantly to active promoters (TssA) or strongly transcribed regions (Tx) in ESCs across all tumours (Figure S14B, 14F, 14J)-contexts typical of housekeeping genes with low methylation plasticity, explaining the minimal CC methylation differences across UHRF1-EM categories (Figure S14A–C, 14E–G, 14I–K).

Together, these results indicate that bivalent ESC chromatin endows EM (and OnF) genes with high methylation plasticity that cancer selectively remodels into UHRF1-associated patterns, stratifying tumours into molecular and prognostic groups (Figure 6; Figure S13; Tables 2–3). CC genes, by contrast, sit in stable active chromatin that constrains methylation remodelling despite substantial transcriptional reprogramming (Figure S14). Thus, in gastric cancer UHRF1-linked methylation at EM/OnF genes is mainly repressive and associated with better outcome (Figure 6A); in renal and adrenocortical cancers, EM-UHRF1-correlated CpG hypermethylation (often in bivalent/enhancer-bivalent contexts) coincides with transcriptional activation and poorer prognosis (Figure 6B-C). The UHRF1–epigenetic co-regulator axis rewires conserved, tissue-specific developmental EM/OnF modules rooted in ESC bivalency to shape tumour biology and clinical behaviour in STAD, KIRC and ACC.

## Discussion

### Clinical Implications of the UHRF1–Embryonic Morphogenesis Axis

Our results establish the UHRF1–embryonic morphogenesis (UHRF1–EM) categorisation as a compact, biologically grounded and context-dependent prognostic framework that performs consistently across three tumour types of distinct developmental origin — gastric adenocarcinoma (STAD), clear-cell renal cell carcinoma (KIRC) and adrenocortical carcinoma (ACC) — whilst retaining a clinically meaningful and mechanistically interpretable directionality specific to each lineage. Crucially, the prognostic relevance of this axis reflects the functional alignment — or anti-alignment — between UHRF1 activity and the transcriptional state of embryonic morphogenesis (EM) and oncofoetal (OnF) gene programmes within each tumour context. In doing so, the UHRF1–EM axis enhances the classification and prognostic stratification of all three tumour types considered, extending beyond current classifiers: it is reproducible across cohorts, remains prognostically informative after adjustment for clinicopathological covariates and established subtype systems, and refines — and in several settings outperforms — existing molecular classifiers, often rivalling pathological stage, particularly in KIRC and ACC. Taken together, these findings indicate that integrating UHRF1 with EM/OnF activity captures a developmental axis insufficiently represented in current classification schemes, thereby providing a clearer and more informative framework for the classification of STAD, KIRC and ACC^31,33,60^.

Pan-cancer analysis of UHRF1 revealed a strongly polarised prognostic landscape and identified Embryonic Morphogenesis (GO:0048598) as the only ontological category that consistently mirrors the reversal of UHRF1 prognostic valence across the three tumour types, thereby providing the basis for the four-class UHRF1–EM framework. Importantly, this axis is largely orthogonal to the mutational landscape: although alterations in TP53, PIK3CA, VHL, PBRM1 and CDH1 are unevenly distributed across classes, none reproduces the survival separation defined by UHRF1–EM membership (Figure S6). In STAD, most MSI and EBV tumours fall within the favourable UH-EML class, whereas the aggressive GS and EMT subtypes are predominantly UL-EMH; in ACC, UH-EMH is enriched for the very-high-risk C1A subgroup, while UL-EML is associated with the more indolent C1B class. Thus, rather than merely recapitulating existing taxonomies, UHRF1–EM refines them, captures risk gradients they do not resolve, and retains independent prognostic value after adjustment for stage, sex and molecular subtype (Figure S9). Crucially, the clinical significance of UHRF1-high status depends not on expression alone, but on whether it is linked to repression of EM/OnF programmes, as in STAD, or to their activation, as in KIRC and ACC.

### UHRF1 Functions Within Context-Dependent Epigenetic Assemblies

The prognostic and transcriptional impact of UHRF1 must be understood within tumour-specific epigenetic assemblies whose composition and functional orientation vary systematically across tissue contexts. Co-expression and antagonism analyses identified recurring partnerships (notably DNMT1, SUV39H1/2, EZH2, and PRC components) and consistent oppositions (EZH1, CBX7, HDAC5, BMI1) that track UHRF1’s own prognostic polarity across STAD, KIRC and ACC. Critically, whilst these co-regulators maintain their cooperative or antagonistic relationships with UHRF1 across all three tumours, their functional consequence — repression or activation of EM and oncofoetal programmes — is inverted in a manner that parallels UHRF1’s own tumour-type-specific role. This inversion implies that the net epigenetic state at developmental loci is determined not by the intrinsic activity of individual factors, but by the relative dominance of cooperating versus opposing complexes within a lineage-specific chromatin landscape. The therapeutic implication is direct: targeting only UHRF1 may prove insufficient or counterproductive if the broader epigenetic assembly is not simultaneously addressed. Understanding which co-regulators dominate in a given tumour type, and how they collectively constrain or facilitate EM/oncofoetal activation, is therefore essential for rational epigenetic intervention.

### Immunological context and translational relevance

The immune–stromal ecosystems associated with UHRF1–EM classes further strengthen its translational relevance. In STAD, the favourable UH-EML class is enriched for cytotoxic and helper lymphoid populations and shows increased PD-L1 expression, consistent with the immune-active biology of MSI and EBV tumours, whereas the adverse UL-EMH class is dominated by stromal components, including CAFs and endothelial cells [20]. In KIRC, UH-EMH is associated with CAFs, Tregs and checkpoint-rich but immunosuppressive infiltrates, while in ACC the adverse class shows limited immune enrichment within an overall suppressive microenvironment (Figure 5A–B). Single-cell analyses further indicate that UHRF1 expression is largely confined to proliferating malignant cells, whereas EM activity is predominantly stromal and immune in origin (Figure 5C–D), suggesting that the EM-high state reflects an oncofoetal niche shaped by tumour-driven microenvironmental reprogramming.

### Additive value in integrated prognostic models

ROC and concordance index analyses further support the clinical utility of the UHRF1–EM framework. Across tumour types, it consistently outperforms molecular subtype systems and, in KIRC and ACC, matches or exceeds pathological stage, particularly in settings where developmental reprogramming appears to be a major determinant of outcome. In TCGA-ACC, UHRF1–EM achieves the highest C-index of any single predictor, a finding reproduced in the independent validation cohort. Notably, across all six cohorts, integrated models combining UHRF1–EM, stage and molecular subtype outperform any individual variable, with UHRF1–EM providing a positive incremental gain in every case (Supplementary Table 1). Together, these results indicate that its prognostic contribution is both additive and non-redundant.

### Mechanistic Basis of the UHRF1–Embryonic Morphogenesis Axis: a Methylation-Centred Conceptual Framework

A major strength of the UHRF1–EM framework is that it captures tissue-specific developmental programmes, rather than generic signatures. In STAD, the EM component is enriched for WNT pathway regulators and gastrointestinal developmental transcription factors; in KIRC, for mediators of renal tubulogenesis, including PITX1, SIX2 and several HOX genes; and in ACC, for genes involved in adrenocortical specification and adrenal stemness (Additional File 2). This lineage specificity places the UHRF1–EM axis within a biologically meaningful embryonic-to-proliferative continuum, consistent with the observation that, in gastric cancer, relapse-associated genes are enriched for developmental and oncofoetal processes, whereas non-relapse genes are dominated by cell-cycle and DNA-repair functions (Figure S10A). Taken together, these data suggest that the prognostic information captured by the UHRF1–EM axis relates less to proliferation per se than to the extent to which developmentally primed transcriptional programmes remain epigenetically plastic in adult tumours.

Within this framework, our data are consistent with the presence of two broad functional modules with different chromatin properties. The first comprises a housekeeping cell-cycle/proliferation module, including replication, mitotic, DNA-repair and canonical MYC/E2F programmes, which is broadly accessible across the life course and can be re-engaged during tissue renewal or repair^61–63^. The second comprises a bivalent oncofoetal module, encompassing developmental regulators, lineage transcription factors, morphogens and EMT/stemness programmes, which appears poised in embryonic stem cells and may retain greater epigenetic plasticity in adult tissues^5,64^. At the chromatin level, cell-cycle loci are therefore likely to resemble constitutively active or broadly accessible promoter states, whereas oncofoetal loci are more likely to retain H3K4me3/H3K27me3 bivalency and the associated susceptibility to tumour-specific epigenetic remodelling. In this light, the UHRF1–EM axis may be viewed as a predominantly methylation-centred developmental stratifier, in which the clinically worst prognosis signal arises mainly from the behaviour of the bivalent developmental module rather than from the proliferative programme that is nevertheless cancerogenic.

Our methylation analyses support this interpretation. Mapping EM CpG clusters onto H1-ESC ChromHMM states showed that tumour-associated hypermethylation at EM/OnF loci in STAD, KIRC and ACC is concentrated in bivalent embryonic chromatin states (TssBiv, BivFlnk, EnhBiv), whereas cell-cycle/proliferation CpGs map mainly to active promoter (TssA) or transcribed states (Tx), typical of housekeeping loci with limited methylation plasticity^65,66^. This distinction is important because it indicates that the principal prognostic methylation signal is not a generic feature of tumour proliferation, but is preferentially associated with developmentally primed loci. Consistently, EM/OnF genes show marked prognosis-linked CpG shifts across UHRF1–EM classes and between normal and tumour tissues, whereas cell-cycle genes display comparatively homogeneous methylation despite substantial transcriptional and prognostic variation. Together, these observations support the view that a bivalency-rich chromatin landscape predisposes developmental genes, but not housekeeping loci, to selective epigenetic remodelling in cancer^28,59,67,68^.

This methylation-centred distinction becomes particularly informative when considered alongside developmental history. Intersecting tumour-dysregulated genes with developmental transcriptomes showed that most EM/OnF genes, together with many dysregulated cell-cycle genes, are embedded within organ-specific embryonic programmes^38,49,51,52^. Consistent with developmental transcriptomic profiling of mouse stomach, kidney and adrenocortical tissues, the genes most tightly associated with UHRF1 in our tumour cohorts are highly expressed during early embryogenesis, when they support progenitor maintenance, WNT-driven patterning and morphogenesis. As tissues mature, these modules are progressively reconfigured, in keeping with the resolution of bivalent chromatin into more stable active or repressed states. Our data therefore support an associative model of developmental epigenetic memory, in which residual bivalency, Polycomb-associated repression and DNA methylation architecture established during organogenesis may influence how UHRF1-linked methylation is distributed across EM/OnF loci in adult tumours.

Within this framework, UHRF1-associated hypermethylation does not have a uniform meaning across lineages. In STAD, the CpG clusters most strongly associated with favourable UH-EML tumours (clusters A/C) are characterised by hypermethylation linked either to EM repression or to the absence of a positive EM–methylation relationship, together with favourable hazard ratios, consistent with a predominantly canonical repressive configuration. This pattern fits well with the developmental trajectory of the gastric epithelium, in which WNT/β-catenin genes and lineage-specifying transcription factors are active during early organogenesis but become progressively restrained and largely silenced in adult tissue, thereby limiting inappropriate intestinal commitment^69,70^. In this setting, UHRF1-linked hypermethylation may therefore be viewed as a marker of a developmentally consolidated, transcriptionally constrained state at EM/OnF loci. By contrast, the poor-prognosis UL-EMH class in STAD shows a partial deviation from this pattern, particularly at cluster D gene-body loci, where hypermethylation is negatively associated with UHRF1 but positively associated with DNMT3A and a TET1-dominant signature, consistent with a more permissive methylation context accompanying developmental programme reactivation^15,71–73^.

In KIRC and ACC, the developmental context appears different. Here, WNT and HOX-related programmes are linked to nephrogenesis, cortical organisation, zonation and regenerative competence, and remain more dynamically regulated across life^74–76^. Consistently, the adverse UH-EMH states in these tumours are associated not merely with higher methylation, but with a methylation architecture centred on EM/OnF loci that tracks with higher EM expression, poor outcome, and broader coupling to methylation–demethylation networks, including DNMT3B/TET3 in KIRC and DNMT3A/TET1 in ACC. These associations do not imply that methylation is directly activating transcription; rather, they suggest that, in these lineages, hypermethylation can coexist with and mark a more plastic, transcriptionally permissive chromatin configuration, particularly at gene-body and enhancer-associated developmental loci^77–79^. The shift towards EnhBiv-dominated regions in KIRC and ACC, together with the relative absence of comparable methylation variation in cell-cycle genes, further supports the view that the relevant methylation signal is linked specifically to developmental chromatin plasticity, rather than to proliferation alone.

The broader epigenetic context is also consistent with this interpretation. A notable finding is the inverse relationship between EZH2 and EZH1, particularly in STAD and ACC, together with the close tracking of EZH2 with UHRF1 across all three tumour types. Although both are catalytic subunits of PRC2, EZH2 is generally associated with de novo H3K27me3 deposition and more dynamic chromatin remodelling, whereas EZH1 is linked to maintenance of pre-existing H3K27me3 and more stable repression^80–84^. In the context of our data, this reciprocal pattern suggests that the EZH2/EZH1 balance may be associated with different degrees of developmental locus plasticity: EZH1-associated states are compatible with more stable repression of EM programmes, whereas EZH2-associated states, especially where they coincide with high UHRF1, are consistent with a more plastic chromatin configuration compatible with EM activation. Likewise, the tissue-specific association of TET enzymes—with TET3 correlation predominating in KIRC and TET1 in ACC— raises the possibility that active methylation turnover may contribute to the permissive methylation states observed in renal and adrenocortical tumours. Again, these patterns should be interpreted associatively: they do not establish mechanism, but they do support the idea that UHRF1-linked methylation outcomes are shaped by the surrounding epigenetic circuitry, rather than by UHRF1 alone.

Finally, the single-cell and deconvolution data indicate that the biological consequences of these methylation states may not be confined to malignant cells. UHRF1 expression is concentrated predominantly in tumour cells, whereas EM/OnF activity is enriched in stromal and immune compartments, particularly CAFs, Tregs and other immune infiltrates. This spatial separation suggests that UHRF1-linked epigenetic remodelling in malignant cells may be associated with broader microenvironmental states, potentially through paracrine signalling rather than direct regulation of EM genes within all cellular compartments^25,85–87^. In this view, the permissive EM-high states observed in KIRC and ACC may correspond, at least in part, to tumour-associated stromal and immune niches enriched for developmental signals, whereas in STAD, repressive EM methylation states may coincide with a less permissive microenvironmental configuration.

Taken together, these findings support a methylation-centred, developmentally informed interpretation of the UHRF1–EM axis. The same EM/OnF loci appear to follow different developmental trajectories across tissues—more stably repressed in the stomach, but more permissively regulated in the kidney and adrenal—and these lineage-specific histories are associated with distinct UHRF1-linked methylation architectures in cancer. On this basis, the prognostic divergence between STAD and KIRC/ACC appears to depend less on hypermethylation per se than on the developmental chromatin context in which that hypermethylation occurs, and on whether it is associated with a relatively repressive or permissive EM state.

This remains a correlative model derived from in silico analyses; direct experimental approaches, including ChIP-seq, locus-specific methylation editing, chromatin accessibility profiling and spatially resolved systems, will be required to determine whether these methylation states are functionally involved in stabilising the lineage-specific tumour phenotypes identified here.

## Conclusions and limitations

Taken together, our findings establish the clinical relevance of the UHRF1–EM axis in robust terms. Across STAD, KIRC and ACC, this framework is reproducible across independent cohorts, remains independently prognostic after adjustment for stage and established molecular subtypes, refines current classification systems, and adds consistent predictive value in integrated models. These results support the view that the UHRF1–EM categorisation captures a clinically meaningful dimension of tumour behaviour that is not adequately represented by existing molecular or pathological classifiers.

Beyond the clinical and molecular robustness of the UHRF1–EM framework, this study is intended to provide an integrative computational framework rather than a definitive causal demonstration of UHRF1-mediated chromatin remodelling. Although this methylation-development model is biologically plausible and supported by the convergence of transcriptomic, methylomic, developmental, chromatin-state and single-cell analyses, it remains a correlative framework derived from in silico analyses and should be further tested through locus-specific functional and epigenetic experiments for definitive confirmation.

Nevertheless, the model proposes a novel epigenetic conceptual framework for interpreting DNA methylation variation in cancer. Rather than viewing hypermethylation as uniformly repressive, it suggests that its biological and clinical meaning may be shaped by developmental context. Beyond their clinical and molecular robustness, our findings support a unifying view in which embryonic bivalent chromatin, tissue-specific epigenetic memory, tumour-associated methylation architectures and EM/OnF transcriptional reprogramming converge to shape lineage-specific tumour behaviour and opposite prognostic states across cancers.

## Supporting information

Description of additional files

## Materials and Methods

1. mRNA expression data source. mRNA expression data for TCGA tumours were retrieved from Firebrowse^88,89^, a TCGA repository maintained by the Broad Institute. RSEM-normalised expression data from IlluminaHiSeq and IlluminaGA platforms were downloaded for tumour and adjacent non-tumour tissues across all 36 tumour types. For analyses across normal, NAT and tumour samples, XENA–UCSC TOIL RSEM TPM data (n=7,862) from the UCSC Toil RNA-seq Recompute resource were downloaded. A second independent validation set was assembled for 10 tumours with significant UHRF1 associations: gastric cancer (GC), ACRG–GEO GSE62254 (n=300)^90^; clear cell renal cell carcinoma (ccRCC), E-MTAB-1980 (Biostudies) (n=100) (https://www.ebi.ac.uk/biostudies/arrayexpress/studies/E-MTAB-1980); adrenocortical tumour (Jouinot, A. *et al* cohort, ACC-2) (n=85)^91^; colorectal cancer (CRC), GEO-GSE87211 (n=203)^92^; papillary renal cell carcinoma (pRCC), GEO-GSE2748 (n=34)^93^; hepatocellular carcinoma (HCC), ICGC-LIRI-JP (n=212)^94^; malignant pleural mesothelioma (MPM), EGA EGAS00001004812 (n=110)^95^; pancreatic ductal adenocarcinoma (PDAC), GEO-GSE21501 (n=123)^96^; sarcoma (SARC), GEO-GSE17618 (n=44)^97^; skin cutaneous melanoma (SKCM), GEO-GSE65904 (n=210)^98^. mRNA expression data for developmental mouse stomach, mouse kidney and human adrenal gland were obtained from GEO-GSE118083^50^, GEO-GSE59130^51^ and ArrayExpress/Biostudies E-MTAB-12492^52^, respectively.
2. Overall survival, clinical data retrieval and survival analysis. ‘Curated survival data’ and ‘Phenotypes’ datasets were downloaded from XENA–UCSC to obtain overall survival (OS), follow-up time and clinical information. Survival data for the validation datasets were obtained from the same accessions listed above. Survival curves and log-rank (Mantel–Cox) P values were generated in GraphPad Prism v8.0.2; forest plots were produced using the R packages survival, survminer and forestplot. For multivariable Cox regression, OS was defined as the interval from diagnosis to death or last follow-up, and survival status as binary. Analyses included only patients with complete clinical and molecular annotation. UHRF1–EM status, pathological stage, molecular subtype and gender were modelled as categorical variables with one reference level each. Univariate Cox models were first used to assess associations between individual covariates and OS. Multivariate Cox models then tested the independent prognostic value of UHRF1–EM after adjustment for clinicopathological variables. Hazard ratios (HRs), 95% confidence intervals (CIs) and two-sided P values were reported; P < 0.05 was considered significant. All analyses were performed in R v4.5.0 using survival, survminer and forestmodel.
3. Patient classification by gene/signature expression and differential expression analysis. Patients were classified by UHRF1 mRNA expression using the cohort median: values above the median were assigned to the UHRF1-high (UH) group and values below to the UHRF1-low (UL) group. The same median split was applied to all other genes/signatures analysed. Differentially expressed (DE) genes were identified by comparing UH vs UL within each tumour type showing significant UHRF1 associations. Gene-level P values were computed using a one-tailed Mann–Whitney U test and adjusted by the Benjamini–Hochberg method. FDR calculations used the wilcoxTest function in the GSALightning R package (v3.15). Genes with q < 0.05 were considered significant. Median expression was then calculated in each group and fold-change (FC) between group medians derived; genes with Log2FC ≥ 0.48542683 or ≤ −0.48542683 were retained. Only genes meeting both q-value and Log2FC thresholds were defined as DE. For the ACRG cohort, differential expression between relapse and non-relapse cases was analysed with the limma R package based on linear modelling with empirical Bayes moderation^99^. Genes with FDR-adjusted P < 0.05 were considered DE; no fold-change threshold was applied to maximise likelihood of observing relapse-associated pathways.
4. Enrichment of UH-DE and UL-DE genes in Human Protein Atlas favourable/unfavourable gene lists (FAV/UNF). Favourable and unfavourable prognostic gene lists (FAV/UNF) were downloaded from the Human Protein Atlas (HPA) portal for each tumour type with significant UHRF1 associations. Enrichment of UH-DE and UL-DE gene sets within the corresponding HPA FAV and UN-FAV lists was tested using one-sided Fisher’s exact tests based on 2×2 contingency tables. P values were computed with the R stats package (v3.6.2).
5. Functional enrichment of differentially expressed genes. DE genes from UHRF1-high (DE-UH) and UHRF1-low (DE-UL) groups were analysed separately for functional enrichment using Metascape^100^. For visualisation (Fig. S2 and reduced version in Fig. 2A), enriched terms were prioritised by selecting the top 100 pathways with a custom composite ranking based on maximum enrichment significance (Max |-LogP|) and variance across cohorts to highlight context-specific signals. Terms were then manually grouped into 7 classes: cell cycle & proliferation; DNA replication & repair; signal transduction; development, EMT & stemness; ECM, adhesion & cytoskeleton; immune, apoptosis & stress response; metabolism, transport & secretion. Redundant parent–child ontological terms were reduced using string-matching filters. Metascape enrichment analysis was also applied to DE genes between relapsed and non-relapsed ACRG patients to identify relapse-associated ontologies.
6. Expression-based signature scoring, combinatorial stratification and association analyses (UHRF1–EM/OnF/Epi/CC). Expression-based signature scoring and stratification were performed within each tumour type. For each signature-Embryonic Morphogenesis (EM), Oncofoetal (OnF), epigenetic (Epi) and cell-cycle (CC)-gene expression values were z-score standardised across patients, and a per-patient score was calculated as the mean z-score of genes in the signature. Patients were dichotomised into High or Low groups using the within-tumour median score. Combinatorial classes were then defined with UHRF1 (e.g. UH–EMH, UH–EML, UL–EMH, UL–EML; similarly for UHRF1–OnF, UHRF1–Epi and UHRF1–CC). Differences in gene or signature expression across classes were tested using the Kruskal–Wallis test followed by Dunn’s multiple-comparisons test. Survival analyses across classes used GraphPad Prism (log-rank Mantel–Cox test). To characterise onco-foetal/stem-like transcriptional programmes across UHRF1–EM classes, an onco-foetal gene set (OnF) was built by combining genes annotated to predefined ontologies/pathways: Transport of Small Molecules (R-HSA-382551), Angiogenesis (GO:0001525), Epithelial to Mesenchymal Transition (GO:0001837), WNT signalling pathways (GO:0016055), Stem Cell Differentiation (GO:0048863), Cell fate Commitment (GO:0045165) and Cell–Cell adhesion (GO:0098609). An OnF signature score was then computed per patient using the same z-score/mean/median workflow, enabling UHRF1–OnF stratification and downstream non-parametric and survival analyses. Associations between UHRF1 and epigenetic mediators were evaluated by Spearman correlation between UHRF1 expression and a curated set of 115 epigenetic regulators in STAD, KIRC and ACC. Significant correlations were defined as Spearman R ≥ 0.2 or ≤ −0.2 with P < 0.0001. Based on these results, epigenetic genes were partitioned into positively associated (EpiCo) and negatively associated (EpiAnti) sets to compute epigenetic signature scores and generate UHRF1–Epi classes (UHRF1 High/Low × Epi High/Low) using the same scoring and median dichotomisation procedure. Correlation structures among UHRF1, epigenetic genes and EM genes were visualised with heatmaps generated using ComplexHeatmap in R, with Spearman r values displayed on a blue (−1) to red (+1) scale. Statistical testing and survival analyses for UHRF1–Epi classes followed the procedures above.
7. Analysis of single-cell RNA sequencing data. Single-cell RNA-seq datasets were obtained from GEO. Gastric primary tumour samples from GSE183904 were analysed; ccRCC single-cell transcriptomes from were used. Raw count matrices and metadata were processed in Seurat (v4/v5). Cells expressing <200 genes or >10% mitochondrial transcripts were excluded. Data were normalised and variance-stabilised using SCTransform (vst.flavor = “v2”), and ∼3,000 highly variable genes were retained. Gene expression counts were then aggregated across all cells within each sample using Seurat’s AggregateExpression function, generating a pseudo-bulk expression matrix for each tumor. Raw pseudo-bulk counts were subsequently normalized using the median-of-ratios method implemented in the DESeq2 package, and log2-transformed normalized counts [log2(normalized counts + 1)] were used for downstream analyses. UHRF1 expression was extracted from the normalized pseudo-bulk matrix and tumors were dichotomized into UHRF1-high (UH) and UHRF1-low (UL) groups using a median split. Embryonic morphogenesis (EM) score was calculated from a predefined EM gene set by averaging scaled gene expression values across samples. UHRF1-EM specific datasets were integrated using the Reciprocal PCA (RPCA) anchoring workflow, followed by PCA and UMAP. Clustering used a shared nearest neighbour (SNN) graph with resolutions from 0.4 to 0.8, selected according to the highest average silhouette score. Cell types were annotated using canonical markers and reference atlases. Differential expression analyses were performed on non-integrated assays (SCT or RNA) to minimise integration-related artefacts.
8. Somatic alterations analysis. Somatic mutation data for TCGA-STAD, KIRC and ACC were retrieved from XENA–UCSC by downloading STAD (n=439) and ACC (n=92) MC3 gene-level non-silent mutation files and using the updated TCGA-KIRC mutation file from the original article^32^. Numbers of patients included in survival and distribution analyses were STAD = 411, KIRC = 334 and ACC = 78. Chi-square statistics were used to test the significance of mutation distributions across UHRF1–EM categories.
9. Immune infiltrate. Quantitative immune infiltration estimates for TCGA samples were downloaded from TIMER 2.0. For each deconvolution algorithm and immune cell type, infiltration values were analysed across the four UHRF1–EM categories. Log2 fold-changes were computed between the best- and worst-prognosis categories for each tumour type, using a threshold equivalent to a 1.5-fold change. Values were normalised by percentile transformation, and within each UHRF1–EM class medians were calculated to obtain one summary value per immune cell type and algorithm. Differences across the four UHRF1–EM categories were tested by one-way ANOVA followed by Dunn’s multiple-comparison test. For each tumour type and immune cell type, hazard ratios were calculated by dichotomising infiltration estimates at the median.
10. ROC curve and C-index computation. *Roc curve*. To evaluate the predictive performance of the four UHRF1–EM categories in gastric, kidney and adrenocortical cancers, TCGA and validation cohorts were analysed against the corresponding molecular and pathological classifications using receiver operating characteristic (ROC) analysis, with OS treated as a categorical outcome. Performance was summarised by sensitivity, specificity and area under the curve (AUC). ROC curves were estimated using repeated permutation/resampling (1001 iterations). At each iteration, the dataset was split into training and test subsets, the model was fitted on the training set, and predicted survival probabilities were generated for the test set. A ROC curve and AUC were then computed for that iteration. The caTools R package was used for data splitting; logistic regression and predicted probabilities were implemented with the R stats package; ROC curves and AUC values were generated with pROC; plots were produced using ggplot2. *C-index*. Additional prognostic performance of UHRF1–EM categories was assessed using Harrell’s concordance index (C-index) for single and combined predictors, including pathological stages and molecular subtypes. OS was the endpoint, with survival time defined as days from diagnosis and event as death. Categorical variables (UHRF1–EM, pathological stage and molecular subtype) were treated as factors. The C-index, which measures agreement between predicted risk and observed survival while accounting for censoring, was defined as the proportion of patient pairs in which the patient with higher predicted risk experienced the event earlier (range 0.5–1.0). Univariate Cox models were fitted for each predictor, and their discriminative ability was quantified by the corresponding C-index. Multivariate Cox models combining pathological stage, molecular subtype and UHRF1–EM were then constructed, and their C-index calculated to assess overall model discrimination. Incremental prognostic value was evaluated by comparing nested models and calculating the difference in C-index (ΔC-index) between models with and without UHRF1–EM. Cox models and concordance were computed using the R survival package. Ninety-five per cent confidence intervals were estimated by non-parametric bootstrap resampling (1000 iterations); where required, optimism-corrected performance was obtained by repeating model fitting and concordance estimation across bootstrap samples and reporting the average corrected C-index.
11. Methylation data source. DNA methylation data from normal tissues were obtained from GEO using datasets generated on platforms fully compatible with TCGA methylation platforms to ensure technical comparability. Gastric normal tissue methylation data were taken from GEO accession: GSE103186^101^, generated with the Illumina Infinium HumanMethylation450 BeadChip. For kidney normal tissue, GSE50874 was used. Pre-processed TCGA tumour methylation data were retrieved from Xena. Datasets were filtered to remove probes with unassigned methylation values (NA) across tumour samples, and only primary tumours were retained. To ensure accurate CpG annotation, the official HumanMethylation450 v1.2 Manifest File for the Illumina Infinium HumanMethylation450K platform was downloaded from Illumina. Chromosome, MapInfo, gene symbol, RefSeq and genomic region annotations were extracted for each cytosine for downstream analyses. Region-specific methylation was visualised by ordering cytosines according to MapInfo within each genomic region and displaying methylation percentiles in heatmaps generated with ComplexHeatmap in R.
12. Correlation and association analysis of DNA methylation across UHRF1-EM categories. Cytosine annotations were integrated with Xena Browser DNA methylation data to quantify CpG methylation with gene-region specificity. For region-based analyses, an “ideal gene” model was used in which CpGs were assigned to Promoter regions (TSS−1500, TSS−200, 5′UTR, 1st exon, 1st exon 5′UTR) or Gene body regions (Body, 3′UTR), representing sequences upstream and downstream of the transcription start site (TSS). For each CpG, methylation values were normalised across patients using z-scores to reduce inter-patient variability and highlight relative differences. Z-scored methylation values were used directly in heatmaps and in CpG–gene expression association analyses. For each CpG, Spearman correlation coefficients (r) were calculated between CpG methylation z-scores and z-scored expression of the corresponding gene using corr.test (Spearman method) in R, reported in the “Gene name/5meC” columns. Depending on the analysis, methylation z-scores were further: (i) converted to percentiles for visualisation in heatmaps (Figs. 6, S13, S14) and line plots (Fig. S15); and/or (ii) summarised for survival analyses by calculating, for each patient, the mean methylation across CpGs within a target region, then dichotomising patients as MH (above cohort median) or ML (below median). Hazard ratios (MH vs ML) were estimated in R using survival, survminer and forestplot. *Heatmap construction*. In methylation heatmaps, rows represented CpGs and columns patients. Patients were ordered hierarchically by: (i) UHRF1–EM category (UH-EMH, UH-EML, UL-EMH, UL-EML), (ii) molecular subtype and (iii) stage. The right panel reported gene-expression metrics for the gene linked to each CpG (first five columns: EM/OnF expression, normalised expression and UX-EMX category), followed by the CpG-specific hazard ratio and CpG–expression Spearman correlations. To visualize UHRF1-specific methylation patterns associated with gene regulation CpGs in the heatmaps were ordered by their correlation between embryonic morphogenesis (EM) gene expression and CpG methylation (EM expr/5meC), followed by correlation between UHRF1 expression and CpG methylation (UHRF1/5meC). Spearman correlations between methylation-derived cluster scores and EM scores were performed at patient level. For each sample, cluster methylation scores were calculated as the mean of CpG-wise z-scored methylation values within each predefined cluster. Scatterplots were used to visualise associations, with linear regression fits and 95% confidence intervals overlaid. For each cluster, correlation coefficients (rho) and P values were reported. CpGs from embryonic morphogenesis (EM), oncofoetal (OnF) and cell-cycle (CC) genes were grouped into correlation-based clusters. EM/OnF CpGs were assigned to four clusters: (A) negative EM expr/5meC correlation; (B) no correlation with either EM or UHRF1; (C) no EM correlation and positive UHRF1/5meC correlation; (D) positive EM expr/5meC correlation. CC CpGs were assigned to three clusters: (A) negative CC expr/5meC correlation; (B) no significant correlation; (C) positive CC expr/5meC correlation. Differences in cluster methylation across normal samples and tumour UHRF1–EM categories were tested using Kruskal–Wallis followed by Dunn’s post hoc test with Benjamini–Hochberg adjustment (dunn.test in R).
13. Epigenomic distribution of CpG clusters. Chromatin state segmentation data were downloaded from the NIH Roadmap Epigenomics Project^66^, which provides 127 reference epigenomes segmented into 15 chromatin states based on combinatorial patterns of core histone modifications, including 16 from ENCODE. For this analysis, the H1 embryonic stem cell (H1-ESC, code E003) chromatin state segmentation file (E003_15_coreMarks_dense.bed.gz) was used, corresponding to the 15-state ChromHMM model for the reference pluripotent epigenome. Overlap between CpG genomic positions and ChromHMM-annotated regions of the H1-ESC epigenome was performed in R (v4.5.1) using data.table package, by matching CpG chromosomal coordinates (chr, start, end) to those of the corresponding chromatin segments in the dense ChromHMM track. This provided a chromatin state annotation for each CpG. Frequency distributions of CpGs across ChromHMM states were computed for each cluster, and significance of distribution differences was assessed by Chi-square test followed by pairwise post hoc comparisons between clusters (A–B, A–C, A–D, B–C, B–D, C–D). Post hoc P values were Bonferroni-adjusted. All analyses and visualisations were performed in R using data.table, dplyr, ggplot2 and readxl. The 15 chromatin states were interpreted according to the original ChromHMM classification.
14. Clustering analysis of developmental gene expression profiles. Gene expression values (z-score normalised across developmental timepoints) were subjected to unsupervised clustering to identify groups of genes with similar temporal dynamics. For hierarchical clustering, used for heatmap visualisation, genes were clustered with the pheatmap package (version 4.5.1). Pairwise gene distances were computed using Euclidean distance, and clusters were merged using Ward’s minimum variance method. Developmental stages (columns) were kept in biological order to preserve temporal structure. Heatmaps were generated with a blue–white–red palette and colour breaks symmetrically centred at zero to balance visualisation of up- and downregulated expression patterns.
15. Language editing. Large language model (LLM)-based tools (ChatGPT, OpenAI) were used exclusively for English language editing, grammar checking, and improvement of readability. The authors reviewed and approved all scientific content, analyses, interpretations, and conclusions.

## Additional Materials

**Additional file 1: Additional file 1.xlsx. List of differentially expressed (DE) genes selected based on Q value and Log2FC.** For each *UHRF1-significative* TCGA tumour, the list of DE-UH and DE-UL genes are reported with the respective log2FC and adjusted P value of UH against UL expression;

**Additional file 2: Additional file 2.xlsx. Enrichment analysis of DE-UH and DE-UL genes.** Sheet: GO_AllLists: Metascape enrichment analysis of DE-UH and DE-UL lists. Pathway Description, Log(qvalue) of enrichment, Enrichment score and Gene lists are reported. Sheet: EM_Genes: worst prognosis associated Embryonic Morphogenesis genes for all 10 *UHRF1-significative* tumours.

**Additional file 3: Additional file 3.xlsx. Distribution of UHRF1-EM classified patients across tumours clinical-pathological classes.** Sheet 1: Distribution of UHRF1-EM patients across clinical variables for 10 UHRF1-signifcative tumours and Chi-square statistics of the distribution, excluded STAD, KIRC, ACC.

**Additional file 4: Additional file 4.xlsx. Distribution of UH and UL patients of gastric ACRG, renal EMTAB-1980 and adrenocortical ACC2 across clinical classes.** Sheet 1: Distribution of UHRF1-high and UHRF1-low patients across clinical variables for tumours and Chi-square/Fischer statistics.

**Additional file 5: Additional file 5.xlsx. Distribution of relevant genetic mutations between UH-UL, EMH-EML and among UHRF1-EM categories.** Sheet: Distribution of TCGA mutations. 5A: distribution of cumulative non-silent mutational burden across UHRF1-EM classes in STAD, TCGA and ACC and relative Kruskal-Wallis Dunn post hoc statistics. 5B, 5C, 5D: distribution of pivotal initiating non-silent mutations indicated by TCGA across UHRF1-EM classes in STAD, TCGA and ACC and relative Chi Square/Fisher Test statistics.

**Additional file 6: Additional file 6.xlsx. ACRG differential gene expression analysis between relapsing and non-relapsing tumours.** Sheet: Relapse annotation. Tumour ID labelled as relapsing (1) or non-relapsing (0). Sheet: DEGs: differentially expressed genes. Light blue: down-regulated in relapse; Light red: up-regulated in relapse. Differentially expressed genes: Log2FC: fold change of expression between relapse and non-relapse groups; t: moderate t test statistics; adj.P.Val: t-test P-value adjusted with Benjamini Hochberg method for multiple comparison; B: log-odds of differential expression. Sheet: Relapse genes Metascape: Metascape enrichment analysis of genes up-regulated in relapse patients, pathway description, Log(qvalue), enrichment score and gene lists are reported for each ontology; Sheet: Non-relapse genes Metascape: Metascape enrichment analysis of genes up-regulated in relapse patients, pathway description, Log(qvalue), enrichment score and gene lists are reported for each ontology;

**Additional file 7: Additional file 7.xlsx. ONCO-signatures of STAD, KIRC, ACC.** Sheet 1: DE-UL oncofoetal gene signatures for STAD (STAD-UL column), KIRC (KIRC-UH column) and ACC (ACC-UH column).

**Additional file 8: Additional file 8.xlsx. Differences of methylation level of embryonic morphogenesis CpGs across UHRF1-EM categories and normal/metaplasic tissue (where available in TCGA_STAD, KIRC and ACC).** For each tumour type (STAD, KIRC and ACC), the corresponding sheet reports Kruskal–Wallis analyses with Dunn’s post hoc pairwise comparisons of DNA methylation levels across all UHRF1–EM categories and normal or metaplastic tissues. Z value and adjusted P value are reported. **Statistical significance of Roadmap Epigenetics annotated genomic context across CpGs clusters A - B - C - D (Chi-Square P-value and post hoc Bonferroni adjustment);** this analysis is available for Embryonic Morphogenesis and Cell Cycle CpGs across the three tumours.

**Additional file 9: Additional file 9.xlsx. DEV_UH and DEV_UL Genes listed according to heatmap gene expression analysis clustering in Figure 7**.

## List of abbreviations

UHRF1: Ubiquitin Like With PHD And Ring Finger Domains 1
EZH2: Enhancer Of Zeste 2 Polycomb Repressive Complex 2 Subunit
EZH1: Enhancer Of Zeste 1 Polycomb Repressive Complex 2 Subunit
ESC: embryonic stem cells
EMT: epithelial to mesenchymal transition
EM: embryonic morphogenesis
OnF: onco-foetal
TCGA: the cancer genome atlas
ACRG: Asian cancer research group
STAD: stomach adenocarcinoma
KIRC: kidney clear cell carcinoma according to TCGA
ccRCC: clear cell renal cell carcinoma
COADREAD: Colorectal cancer
LIHC: liver hepatocellular carcinoma
PAAD: pancreatic adenocarcinoma
ACC: adrenocortical carcinoma
MESO: mesothelioma
KIRP: papillary kidney carcinoma according to TCGA
CRC: colorectal cancer
pRCC: papillary renal cell carcinoma
HCC: hepatocellular carcinoma
MPM: malignant pleural mesothelioma
PDAC: pancreatic ductal adenocarcinoma
SARC: sarcoma
SKCM: skin cell melanoma
mRNA: messenger RNA
GEO: gene expression omnibus
OS: overall survival
HR: hazard ratio
CI: confidence interval
UH: UHRF1-*high*
UL: UHRF1-*low*
DE: differentially expressed
EZH2H: EZH2-*high*
EZH2L: EZH2-*low*
FC: fold change
UH-DE: differentially expressed (upregulated) genes in UH patients
UL-DE: differentially expressed (upregulated) genes in UL patients
FDR: false discovery rate
HPA: human protein atlas
FAV/UNFAV: favourable/unfavourable
GO: gene ontology
BP/WP: best prognosis/worst prognosis
EMH: EM-*high*
EM: EM-*low*
CC: cell cycle
Epi: epigenetic
scRNA-seq: single-cell RNA sequencing
RPCA: reciprocal principal component analysis
PCA: principal component analysis
UMAP: uniform manifold approximation and projection
SNN: shared nearest neighbour
ANOVA: analysis of variance
TIMER: Tumour IMmune Extimation Resource
ROC: receiver operating characteristics
AUC: area under the curve
TSS: transcription start site
UTR: untranslated
MH/ML: methylation-*high/low*
pT: pathological T stage
pN: pathological N stage
TP53: tumor protein 53
PIK3CA: Phosphatidylinositol-4,5-Bisphosphate 3-Kinase Catalytic Subunit Alpha
MLH1: MutL Homolog 1
CDH1: Cadherin 1
VHL: Von Hippel-Lindau Tumor Suppressor
PBRM1: Polybromo 1
SETD2: SET Domain Containing 2, Histone Lysine Methyltransferase
MSI: microsatellite instability
GS: genome stability
CIN: chromosomal instability
DNMT1/3A/3B: DNA methyl transferase 1/3A/3B
SUV39H1/H2: Suppressor Of Variegation 3-9 Homolog 1/2
CBX: chromobox
CAF: cancer associated fibroblasts
PD-L1: Programmed Cell Death 1 Ligand 1
PD-1: Programmed Cell Death 1
Tregs: regulatory T cells
Tfh: T follicular helper cells
CTLA4: Cytotoxic T-Lymphocyte Associated Protein 4
5meC: 5-methyl-cyosine
Expr: expression
AURKB: Aurora Kinase B
PLK1/4: Polo Like Kinase ¼
HDAC5: Histone Deacetylase 5
TET1/2/3: Ten-Eleven Translocation 1/2/3 Gene Protein
SFRP2: Secreted Frizzled Related Protein 2
DKK1: Dickkopf Wnt Signaling Pathway Inhibitor 1
GPC3: Glypican 3
TGFB2: Transforming Growth Factor Beta 2
PITX1: Paired Like Homeodomain 1
WNT16: Wingless-Type MMTV Integration Site Family, Member 16
GREM1: Gremlin 1, DAN Family BMP Antagonist
HOX: homeobox
SIX2: Sine Oculis Homeobox Homolog 2
DEV: development
Barx1: Homeobox Protein BarH-Like 1
Gli1/3: Glioma-Associated Oncogene Homolog 1/3 (Zinc Finger Protein)
Sox7: SRY-Box Transcription Factor 7
Foxf1: Forkhead Box 1
Rspo2/3: R-Spondin 2/3
EYA1/2: Eyes Absent Homolog ½
SALL4: Spalt Like Transcription Factor 4
LHX1: LIM Homeobox 1
LEF1: Lymphoid Enhancer Binding Factor 1
TSSBiv: Bivalent TSS
BivFlnk: Flaning Bivalent Region
EnhBiv: Bivalent Enhancer
TssA: active TSS
Tx: Strongly transcribed regions
TssAFlnk: region flanking actve TSS
TxFlnk: region flanking strongly transcribed region
TxWk: weakly transcribed region
EnhG: strong enhancer
Enh: enhancer
ZNF/Repeats: Zing finger domains/repeated elements
Het: heterochromatin
ReprPC: region repressed by Polycomb
ReprPCWk: region weakly repressed by polycomb
Quiesc: chromatin giving not relevant signals

## Data availability

The datasets generated and analyzed during the current study are available from the corresponding author on reasonable request. Dataset that were analysed to obtain our results are listed in materials and methods, section “mRNA Expression data source” for gene expression in tumour, normal tissue and developmental stages, “Overall survival, clinical data retrieval and survival analysis” for survival datasets, “Analysis of single-cell RNA sequencing data” for single cell RNA-seq datasets, “Somatic alterations analysis” for mutational data, “Methylation data source” for methylation datasets, “Epigenomic distribution of CpG clusters” for chromatin annotation in H1 embryonic stem cells genome.

## Data Availability Details

All datasets analysed in this study are publicly available. TCGA RNA-seq expression, clinical, survival, somatic mutation and DNA methylation data were obtained from FireBrowse and the UCSC Xena platform. Normal, tumour and normal-adjacent tissue transcriptomic data were retrieved from the UCSC Toil RNA-seq Recompute resource.

Independent validation cohorts included gastric cancer (ACRG cohort, GSE62254), clear cell renal cell carcinoma (E-MTAB-1980), adrenocortical carcinoma (Jouinot, A. *et al* cohort, ACC-2), colorectal cancer (GSE87211), papillary renal cell carcinoma (GSE2748), hepatocellular carcinoma (ICGC-LIRI-JP), malignant pleural mesothelioma (EGAS00001004812), pancreatic ductal adenocarcinoma (GSE21501), sarcoma (GSE17618), and skin cutaneous melanoma (GSE65904).

Developmental transcriptomic datasets comprised mouse stomach development (GSE118083), mouse kidney development (GSE59130), and human adrenal gland development (E-MTAB-12492). Single-cell RNA sequencing datasets included gastric cancer samples (GSE183904) and clear cell renal cell carcinoma samples (GSE207493).

DNA methylation data from normal gastric and kidney tissues were obtained from GSE103186 and GSE50874, respectively, while TCGA tumour methylation data were downloaded from UCSC Xena. Immune cell infiltration estimates were retrieved from TIMER 2.0. Chromatin-state annotations were obtained from the NIH Roadmap Epigenomics Project (H1 embryonic stem cell reference epigenome, E003). Functional enrichment analyses were performed using Metascape, and favourable and unfavourable prognostic gene lists were obtained from the Human Protein Atlas.

All datasets are available from the corresponding public repositories under the accession numbers reported above and in the Materials and Methods section.

## Competing interests

The authors declare that they have no competing interests

## Funding

I.M.B. discloses support for the research of this work from institutional resources of the University of Insubria (Varese, Italy) and the National Research Council (CNR) Flagship Project 2013, Subproject No. 6.8, *Decoding the epigenetic basis of DNA hypo- and hypermethylation in cancer*.

L.M. discloses support for the research of this work from the NextGenerationEU project PNRR “PRP@CERIC” (IR0000028).

S.S. discloses support for the research of this work from Fondazione AIRC per la Ricerca sul Cancro (AIRC Investigator Grant 2022, ID 27233).

S.G. discloses support for the research of this work from Fondazione AIRC per la Ricerca sul Cancro (AIRC Investigator Grant 2024, ID 27531).

D.F. discloses support for the research of this work from institutional resources of the University of Insubria (Varese, Italy).

G.F. discloses support for the research of this work from institutional resources of the University of Naples Federico II (Naples, Italy).

## Authors’ contributions

J.B. conceptualization, substantial contribution to the data analysis, writing and revising the manuscript

L.M. conceptualization, substantial contribution to the data analysis, revising the manuscript

F.M. conceptualization, study design, data analysis

M.S. substantial contribution to the data analysis

C.G. contribution to data analysis

M.F. substantial contribution to the data analysis

E.M. conceptualization, study design

S.C. contributions to the conception and design of the manuscript for gastric cancer aspects

C.M. contributions to the conception and design of the manuscript for gastric cancer aspects

D.C. contribution to the data analysis and revision of the manuscript

F.D. critically revising the manuscript for intellectual content and for data analysis

P.M. contributions to the drafting and revision of the manuscript for gastric cancer aspects

F.A. contributions to the drafting and revision of the manuscript for gastric cancer aspects

V.C. revising the manuscript

M.T. contributions to the conception and design for adrenocortical carcinoma

G.M. critically revising the manuscript for intellectual content

T.C. critically revising the manuscript for intellectual content

T.A. contribution to the data analysis and revision of the manuscript

G.F. contributions to the conception and design of the manuscript for gastric cancer aspects

S.S. contributions to the conception and design for adrenocortical carcinoma

S.G. contributions to the conception and design for gastric cancer

M.F. substantial contribution to the data analysis

D.F. conceptualization, study design

I.M.B. conceived the idea, designed the study, oversaw the work, and funding

## Acknowledgements

## Notes

### Competing Interest Statement

The authors have declared no competing interest.

## Bibliography

1. Mancini, M., Magnani, E., Macchi, F. & Bonapace, I.M. The multi-functionality of UHRF1: epigenome maintenance and preservation of genome integrity. Nucleic Acids Res 49, 6053–6068 (2021).

2. Ashraf, W. et al. The epigenetic integrator UHRF1: on the road to become a universal biomarker for cancer. Oncotarget 8, 51946–51962 (2017).

3. Alhosin, M. et al. Signalling pathways in UHRF1-dependent regulation of tumor suppressor genes in cancer. J Exp Clin Cancer Res 35, 174 (2016).

4. Sharma, A., Blériot, C., Currenti, J. & Ginhoux, F. Oncofetal reprogramming in tumour development and progression. Nat Rev Cancer 22, 593–602 (2022).

5. Macrae, T.A., Fothergill-Robinson, J. & Ramalho-Santos, M. Regulation, functions and transmission of bivalent chromatin during mammalian development. Nat Rev Mol Cell Biol 24, 6–26 (2023).

6. Qin, W. et al. DNA methylation requires a DNMT1 ubiquitin interacting motif (UIM) and histone ubiquitination. Cell Res 25, 911–29 (2015).

7. Benavente, C.A. et al. Chromatin remodelers HELLS and UHRF1 mediate the epigenetic deregulation of genes that drive retinoblastoma tumor progression. Oncotarget 5, 9594–608 (2014).

8. Han, M. et al. A role for LSH in facilitating DNA methylation by DNMT1 through enhancing UHRF1 chromatin association. Nucleic Acids Res 48, 12116–12134 (2020).

9. Sharif, J. et al. The SRA protein Np95 mediates epigenetic inheritance by recruiting Dnmt1 to methylated DNA. Nature 450, 908–12 (2007).

10. Li, T. et al. Structural and mechanistic insights into UHRF1-mediated DNMT1 activation in the maintenance DNA methylation. Nucleic Acids Res 46, 3218–3231 (2018).

11. Rajakumara, E. et al. PHD finger recognition of unmodified histone H3R2 links UHRF1 to regulation of euchromatic gene expression. Mol Cell 43, 275–284 (2011).

12. Liu, Y. et al. DNA hypomethylation promotes UHRF1-and SUV39H1/H2-dependent crosstalk between H3K18ub and H3K9me3 to reinforce heterochromatin states. Mol Cell 85, 394–412.e12 (2025).

13. Kamel, E.M. et al. Disrupting the epigenetic alliance: structural insights and therapeutic strategies targeting DNMT1-UHRF1. Funct Integr Genomics 25, 194 (2025).

14. Zhao, Q. et al. Dissecting the precise role of H3K9 methylation in crosstalk with DNA maintenance methylation in mammals. Nat Commun 7, 12464 (2016).

15. Yamaguchi, K. et al. Non-canonical functions of UHRF1 maintain DNA methylation homeostasis in cancer cells. Nat Commun 15, 2960 (2024).

16. Magnani, E. et al. uhrf1 and dnmt1 loss induces an immune response in zebrafish livers due to viral mimicry by transposable elements. Front Immunol 12, 627926 (2021).

17. Kim, A. & Benavente, C.A. Oncogenic Roles of UHRF1 in Cancer. Epigenomes 8(2024).

18. Babbio, F. et al. The SRA protein UHRF1 promotes epigenetic crosstalks and is involved in prostate cancer progression. Oncogene 31, 4878–87 (2012).

19. Sabatino, L. et al. UHRF1 coordinates peroxisome proliferator activated receptor gamma (PPARG) epigenetic silencing and mediates colorectal cancer progression. Oncogene (2012).

20. Niinuma, T. et al. UHRF1 depletion and HDAC inhibition reactivate epigenetically silenced genes in colorectal cancer cells. Clin Epigenetics 11, 70 (2019).

21. Kong, X. et al. Defining UHRF1 Domains that Support Maintenance of Human Colon Cancer DNA Methylation and Oncogenic Properties. Cancer Cell 35, 633–648.e7 (2019).

22. Xue, B. et al. Epigenetic mechanism and target therapy of UHRF1 protein complex in malignancies. Onco Targets Ther 12, 549–559 (2019).

23. Abdullah, O. et al. Thymoquinone Is a Multitarget Single Epidrug That Inhibits the UHRF1 Protein Complex. Genes (Basel) 12(2021).

24. Tan, L. et al. Aberrant cytoplasmic expression of UHRF1 restrains the MHC-I-mediated anti-tumor immune response. Nat Commun 15, 8569 (2024).

25. Gu, Y. et al. UHRF1 drives subtype-independent aggressiveness and immune evasion in small cell lung cancer through PRC2 interactions. iScience 29, 115475 (2026).

26. Bernstein, B.E. et al. A bivalent chromatin structure marks key developmental genes in embryonic stem cells. Cell 125, 315–26 (2006).

27. Ohm, J.E. et al. A stem cell-like chromatin pattern may predispose tumor suppressor genes to DNA hypermethylation and heritable silencing. Nat Genet 39, 237–42 (2007).

28. Dunican, D.S. et al. Bivalent promoter hypermethylation in cancer is linked to the H327me3/H3K4me3 ratio in embryonic stem cells. BMC Biol 18, 25 (2020).

29. Kim, K.Y. et al. Uhrf1 regulates active transcriptional marks at bivalent domains in pluripotent stem cells through Setd1a. Nat Commun 9, 2583 (2018).

30. Digre, A. & Lindskog, C. The human protein atlas-Integrated omics for single cell mapping of the human proteome. Protein Sci 32, e4562 (2023).

31. Bass, A.J. et al. Comprehensive molecular characterization of gastric adenocarcinoma. Nature 513, 202–209 (2014).

32. Ricketts, C.J. et al. The Cancer Genome Atlas Comprehensive Molecular Characterization of Renal Cell Carcinoma. Cell Rep 43, 113063 (2024).

33. Zheng, S. et al. Comprehensive Pan-Genomic Characterization of Adrenocortical Carcinoma. Cancer Cell 29, 723–736 (2016).

34. Puliga, E., Corso, S., Pietrantonio, F. & Giordano, S. Microsatellite instability in Gastric Cancer: Between lights and shadows. Cancer Treat Rev 95, 102175 (2021).

35. Peng, Y. et al. PLK1 maintains DNA methylation and cell viability by regulating phosphorylation-dependent UHRF1 protein stability. Cell Death Discov 9, 367 (2023).

36. Sun, W., Jiang, Z. & Deng, Z. Comprehensive analysis of the tumor immune microenvironment in gastric cancer and peritoneal metastasis based on single-cell RNA sequencing analysis. Sci Rep 15, 32090 (2025).

37. Shapiro, D.D. et al. Understanding the Tumor Immune Microenvironment in Renal Cell Carcinoma. Cancers (Basel) 15(2023).

38. Kumar, V. et al. Single-Cell Atlas of Lineage States, Tumor Microenvironment, and Subtype-Specific Expression Programs in Gastric Cancer. Cancer Discov 12, 670–691 (2022).

39. Yu, Z. et al. Integrative Single-Cell Analysis Reveals Transcriptional and Epigenetic Regulatory Features of Clear Cell Renal Cell Carcinoma. Cancer Res 83, 700–719 (2023).

40. Cheng, Y.Y. et al. Frequent epigenetic inactivation of secreted frizzled-related protein 2 (SFRP2) by promoter methylation in human gastric cancer. Br J Cancer 97, 895–901 (2007).

41. Shi, T. et al. DKK1 Promotes Tumor Immune Evasion and Impedes Anti-PD-1 Treatment by Inducing Immunosuppressive Macrophages in Gastric Cancer. Cancer Immunol Res 10, 1506–1524 (2022).

42. Hishinuma, M. et al. Hepatocellular oncofetal protein, glypican 3 is a sensitive marker for α-fetoprotein-producing gastric carcinoma. Histopathology 49, 479–486 (2006).

43. Han, B., Fang, T., Wang, Y., Zhang, Y. & Xue, Y. TGFβ2 is a Prognostic Biomarker for Gastric Cancer and is Associated With Methylation and Immunotherapy Responses. Frontiers in Genetics Volume 13 - 2022(2022).

44. Zhang, Y., Zhang, Z., Zhang, W., Hu, H. & Bao, G. Upregulated Transcription Factor PITX1 Predicts Poor Prognosis in Kidney Renal Clear Cell Carcinoma-Based Bioinformatic Analysis and Experimental Verification. Dis Markers 2021, 7694239 (2021).

45. Cheng, N., Li, H., Han, Y. & Sun, S. Transcription factor Six2 induces a stem cell-like phenotype in renal cell carcinoma cells. FEBS Open Bio 9, 1808–1816 (2019).

46. van Vlodrop, I.J. et al. Prognostic significance of Gremlin1 (GREM1) promoter CpG island hypermethylation in clear cell renal cell carcinoma. Am J Pathol 176, 575–84 (2010).

47. Francis, J.C. et al. HOX genes promote cell proliferation and are potential therapeutic targets in adrenocortical tumours. Br J Cancer 124, 805–816 (2021).

48. Caglar, H.O. & Biray Avci, C. Alterations of cell cycle genes in cancer: unmasking the role of cancer stem cells. Mol Biol Rep 47, 3065–3076 (2020).

49. Miao, Z. et al. Single cell regulatory landscape of the mouse kidney highlights cellular differentiation programs and disease targets. Nat Commun 12, 2277 (2021).

50. Li, X. et al. A time-resolved multi-omic atlas of the developing mouse stomach. Nat Commun 9, 4910 (2018).

51. Brunskill, E.W. et al. Single cell dissection of early kidney development: multilineage priming. Development 141, 3093–101 (2014).

52. Del Valle, I., et al. An integrated single-cell analysis of human adrenal cortex development. JCI Insight 8(2023).

53. Kim, B.M., Buchner, G., Miletich, I., Sharpe, P.T. & Shivdasani, R.A. The stomach mesenchymal transcription factor Barx1 specifies gastric epithelial identity through inhibition of transient Wnt signaling. Dev Cell 8, 611–22 (2005).

54. Khoshdel Rad, N., Aghdami, N. & Moghadasali, R. Cellular and Molecular Mechanisms of Kidney Development: From the Embryo to the Kidney Organoid. Front Cell Dev Biol 8, 183 (2020).

55. Wang, Y., Zhou, C.J. & Liu, Y. Wnt Signaling in Kidney Development and Disease. Prog Mol Biol Transl Sci 153, 181–207 (2018).

56. Lescher, B., Haenig, B. & Kispert, A. sFRP-2 is a target of the Wnt-4 signaling pathway in the developing metanephric kidney. Dev Dyn 213, 440–51 (1998).

57. Lyraki, R. & Schedl, A. Adrenal cortex renewal in health and disease. Nat Rev Endocrinol 17, 421–434 (2021).

58. Kundaje, A. et al. Integrative analysis of 111 reference human epigenomes. Nature 518, 317–30 (2015).

59. Bernhart, S.H. et al. Changes of bivalent chromatin coincide with increased expression of developmental genes in cancer. Sci Rep 6, 37393 (2016).

60. Network, C.G.A.R. Comprehensive molecular characterization of clear cell renal cell carcinoma. Nature 499, 43–9 (2013).

61. Liu, H. et al. Redeployment of Myc and E2f1-3 drives Rb-deficient cell cycles. Nat Cell Biol 17, 1036–48 (2015).

62. Mjelle, R. et al. Cell cycle regulation of human DNA repair and chromatin remodeling genes. DNA Repair (Amst) 30, 53–67 (2015).

63. Konagaya, Y., Rosenthal, D., Ratnayeke, N., Fan, Y. & Meyer, T. An intermediate Rb-E2F activity state safeguards proliferation commitment. Nature 631, 424–431 (2024).

64. Kumar, D., Cinghu, S., Oldfield, A.J., Yang, P. & Jothi, R. Decoding the function of bivalent chromatin in development and cancer. Genome Res 31, 2170–2184 (2021).

65. Ernst, J. & Kellis, M. ChromHMM: automating chromatin-state discovery and characterization. Nat Methods 9, 215–6 (2012).

66. Chromhmm. http://egg2.wustl.edu/roadmap/data/byFileType/chromhmmSegmentations/ChmmModels/coreMarks/jointModel/final/.

67. Lu, Y. et al. Pan-cancer analysis revealed H3K4me1 at bivalent promoters premarks DNA hypermethylation during tumor development and identified the regulatory role of DNA methylation in relation to histone modifications. BMC Genomics 24, 235 (2023).

68. Zaidi, S.K. et al. Bivalent Epigenetic Control of Oncofetal Gene Expression in Cancer. Mol Cell Biol 37(2017).

69. Simmini, S. et al. Transformation of intestinal stem cells into gastric stem cells on loss of transcription factor Cdx2. Nat Commun 5, 5728 (2014).

70. Radulescu, S. et al. Acute WNT signalling activation perturbs differentiation within the adult stomach and rapidly leads to tumour formation. Oncogene 32, 2048–57 (2013).

71. Baubec, T. et al. Genomic profiling of DNA methyltransferases reveals a role for DNMT3B in genic methylation. Nature 520, 243–7 (2015).

72. Song, C.X. et al. Selective chemical labeling reveals the genome-wide distribution of 5-hydroxymethylcytosine. Nat Biotechnol 29, 68–72 (2011).

73. Neri, F. et al. Intragenic DNA methylation prevents spurious transcription initiation. Nature 543, 72–77 (2017).

74. Kamei, C.N., Gallegos, T.F., Liu, Y., Hukriede, N. & Drummond, I.A. Wnt signaling mediates new nephron formation during zebrafish kidney regeneration. Development 146(2019).

75. Di-Poï, N., Zákány, J. & Duboule, D. Distinct roles and regulations for HoxD genes in metanephric kidney development. PLoS Genet 3, e232 (2007).

76. Meng, P., Zhu, M., Ling, X. & Zhou, L. Wnt signaling in kidney: the initiator or terminator? J Mol Med (Berl) 98, 1511–1523 (2020).

77. Yang, X. et al. Gene body methylation can alter gene expression and is a therapeutic target in cancer. Cancer Cell 26, 577–90 (2014).

78. Lister, R. et al. Human DNA methylomes at base resolution show widespread epigenomic differences. Nature 462, 315–22 (2009).

79. Charlet, J. et al. Bivalent Regions of Cytosine Methylation and H3K27 Acetylation Suggest an Active Role for DNA Methylation at Enhancers. Mol Cell 62, 422–431 (2016).

80. Lee, S.H. et al. The role of EZH1 and EZH2 in development and cancer. BMB Rep 55, 595–601 (2022).

81. Boros, J., Arnoult, N., Stroobant, V., Collet, J.F. & Decottignies, A. Polycomb repressive complex 2 and H3K27me3 cooperate with H3K9 methylation to maintain heterochromatin protein 1α at chromatin. Mol Cell Biol 34, 3662–74 (2014).

82. Zheng, Y. et al. A pan-cancer analysis of CpG Island gene regulation reveals extensive plasticity within Polycomb target genes. Nat Commun 12, 2485 (2021).

83. Margueron, R. et al. Ezh1 and Ezh2 maintain repressive chromatin through different mechanisms. Mol Cell 32, 503–18 (2008).

84. Shen, X. et al. EZH1 mediates methylation on histone H3 lysine 27 and complements EZH2 in maintaining stem cell identity and executing pluripotency. Mol Cell 32, 491–502 (2008).

85. Topper, M.J. et al. Epigenetic Therapy Ties MYC Depletion to Reversing Immune Evasion and Treating Lung Cancer. Cell 171, 1284–1300.e21 (2017).

86. Elyada, E. et al. Cross-Species Single-Cell Analysis of Pancreatic Ductal Adenocarcinoma Reveals Antigen-Presenting Cancer-Associated Fibroblasts. Cancer Discov 9, 1102–1123 (2019).

87. Sharma, A. et al. Onco-fetal Reprogramming of Endothelial Cells Drives Immunosuppressive Macrophages in Hepatocellular Carcinoma. Cell 183, 377–394.e21 (2020).

88. Deng, M., Brägelmann, J., Kryukov, I., Saraiva-Agostinho, N. & Perner, S. FirebrowseR: an R client to the Broad Institute’s Firehose Pipeline. Database (Oxford) 2017(2017).

89. Firebrowse. https://firebrowse.org/.

90. Cristescu, R. et al. Molecular analysis of gastric cancer identifies subtypes associated with distinct clinical outcomes. Nat Med 21, 449–56 (2015).

91. Jouinot, A. et al. Transcriptome in paraffin samples for the diagnosis and prognosis of adrenocortical carcinoma. Eur J Endocrinol 186, 607–617 (2022).

92. Hu, Y. et al. Colorectal cancer susceptibility loci as predictive markers of rectal cancer prognosis after surgery. Genes Chromosomes Cancer 57, 140–149 (2018).

93. Yang, X.J. et al. A molecular classification of papillary renal cell carcinoma. Cancer Res 65, 5628–37 (2005).

94. Zeng, C. et al. A stratification model of hepatocellular carcinoma based on expression profiles of cells in the tumor microenvironment. BMC Cancer 22, 613 (2022).

95. Lifting the curtain on molecular differences between malignant pleural mesotheliomas. Nat Genet 55, 540–541 (2023).

96. Stratford, J.K. et al. A six-gene signature predicts survival of patients with localized pancreatic ductal adenocarcinoma. PLoS Med 7, e1000307 (2010).

97. Volchenboum, S.L. et al. Gene Expression Profiling of Ewing Sarcoma Tumors Reveals the Prognostic Importance of Tumor-Stromal Interactions: A Report from the Children’s Oncology Group. J Pathol Clin Res 1, 83–94 (2015).

98. Whole-genome expression analysis of melanoma tumor biopsies from a population-based cohort. (2015).

99. Smyth, G.K. Linear models and empirical bayes methods for assessing differential expression in microarray experiments. Stat Appl Genet Mol Biol 3, Article3 (2004).

100. Zhou, Y. et al. Metascape provides a biologist-oriented resource for the analysis of systems-level datasets. Nat Commun 10, 1523 (2019).

101. Huang, K.K. et al. Genomic and Epigenomic Profiling of High-Risk Intestinal Metaplasia Reveals Molecular Determinants of Progression to Gastric Cancer. Cancer Cell 33, 137–150.e5 (2018).

