## Supplementary material for "A multilayered in silico analysis links UHRF1, DNA methylation and developmental chromatin memory to lineage-dependent prognosis in gastric, renal and adrenal cancers": Suppl. Table 2

| **TUMOUR**  **TYPE** | **DOMINANT**  **FUNCTIONAL**  **EFFECT OF UHRF1 LINKED METHYLATION** | **CLUSTER FEATURES**  **(FIGURE 6A-C AND**  **SUPPL. FIGURE 13)** | **PROGNOSTIC EXTREME**  **IDENTIFIED BY**  **UHRF1 LINKED**  **HYPERMETHYLATION** |
| --- | --- | --- | --- |
| STAD | Predominantly  repressive | Clusters A/C: UH-EML hypermethyl.,  EM repression and HR<1  Cluster D: UL-EMH hypermethyl.,  EM activation and HR>1  Cluster B: no significance | UH-EML  (best prognosis class) |
| KIRC | Predominantly  activating | Clusters A: UH-EMH hypo-methyl., UL-EML hyper-methyl.  Cluster C/D: hyprmethyl.,  UH-EMH EM overexpr.  and HR>1  Cluster B: no significance | UH-EMH  (worst prognosis class) |
| ACC | Predominantly  activating | Similar to KIRC:  Cluster C/D: UH-EMH hypermethylation  EM overexpr., HR>1  Cluster B: no significance | UH-EMH  (worst prognosis class) |

**Suppl. Table 2: Tumour‑specific impact of UHRF1‑linked methylation on EM/OnF programmes**
