## Supplementary material for "A multilayered in silico analysis links UHRF1, DNA methylation and developmental chromatin memory to lineage-dependent prognosis in gastric, renal and adrenal cancers": Suppl. Table 3

| **CLUSTER** | **STAD: UHRF1–5MEC AND CO‑REGULATOR PATTERNS**  **(FIGURE 6A)** | **KIRC/ACC: UHRF1–5MEC AND CO‑REGULATOR PATTERNS**  **(FIGURE 6B–C)** | **INTEGRATED PROGNOSTIC IMPLICATION**  **(FIGURES 6A–C,**  **SUPPL. FIGURE 13)** |
| --- | --- | --- | --- |
| A | UHRF1–5meC positive   - co‑correlated with EZH2, SUV39H1/2, HELLS, WHSC1, CBX3, SMARCA4, CBX8, AURKB, PLK1/4, CHAF1A - negative correlation with BMI1, CBX7, HDAC5, EZH1 - DNMT3A/3B and TET1/3 positive in subsets | UHRF1–5meC weak or negative   - opposite trends for HDAC5, CBX7, BMI1, EZH1 (positive - modest hypermethylation in UL‑EML | Hypermethylation linked to   - EM repression - improved outcome in all three tumours |
| B | No stable associations with   - UHRF1–5meC - EMexpr–5meC - prognosis | Similarly weak, non‑directional correlations | No consistent  prognostic effect |
| C | UHRF1–5meC parallels co‑epigenetic factor–5meC;   - anti‑epigenetic factors inversely correlated; - hypermethylation in UH‑EML | UHRF1–5meC and co‑factor–5meC positive;   - hypermethylation in UH‑EMH with EM activation; - emerging TET1/TET3 correlations | Favourable in STAD;  adverse in KIRC/ACC |
| D | UHRF1, DNMT1, EZH2, SUV39H1/2, CBX3 negatively correlated with methylation;   - DNMT3A, TET1, HDAC5, CBX7, EZH1 positively correlated; - hypermethylation in UL‑EMH | UHRF1, DNMT1, DNMT3A, EZH2, SUV39H1 positively correlated with methylation;   - strong TET1/TET3–5meC correlations, consistent with h5meC‑rich, active chromatin; - methylation peaks in UH‑EMH | Strongly adverse in all tumours, particularly KIRC and ACC |

**Table 4. UHRF1-centred epigenetic networks across clusters and tumour types**
