## Supplementary Figures for "A multilayered in silico analysis links UHRF1, DNA methylation and developmental chromatin memory to lineage-dependent prognosis in gastric, renal and adrenal cancers"

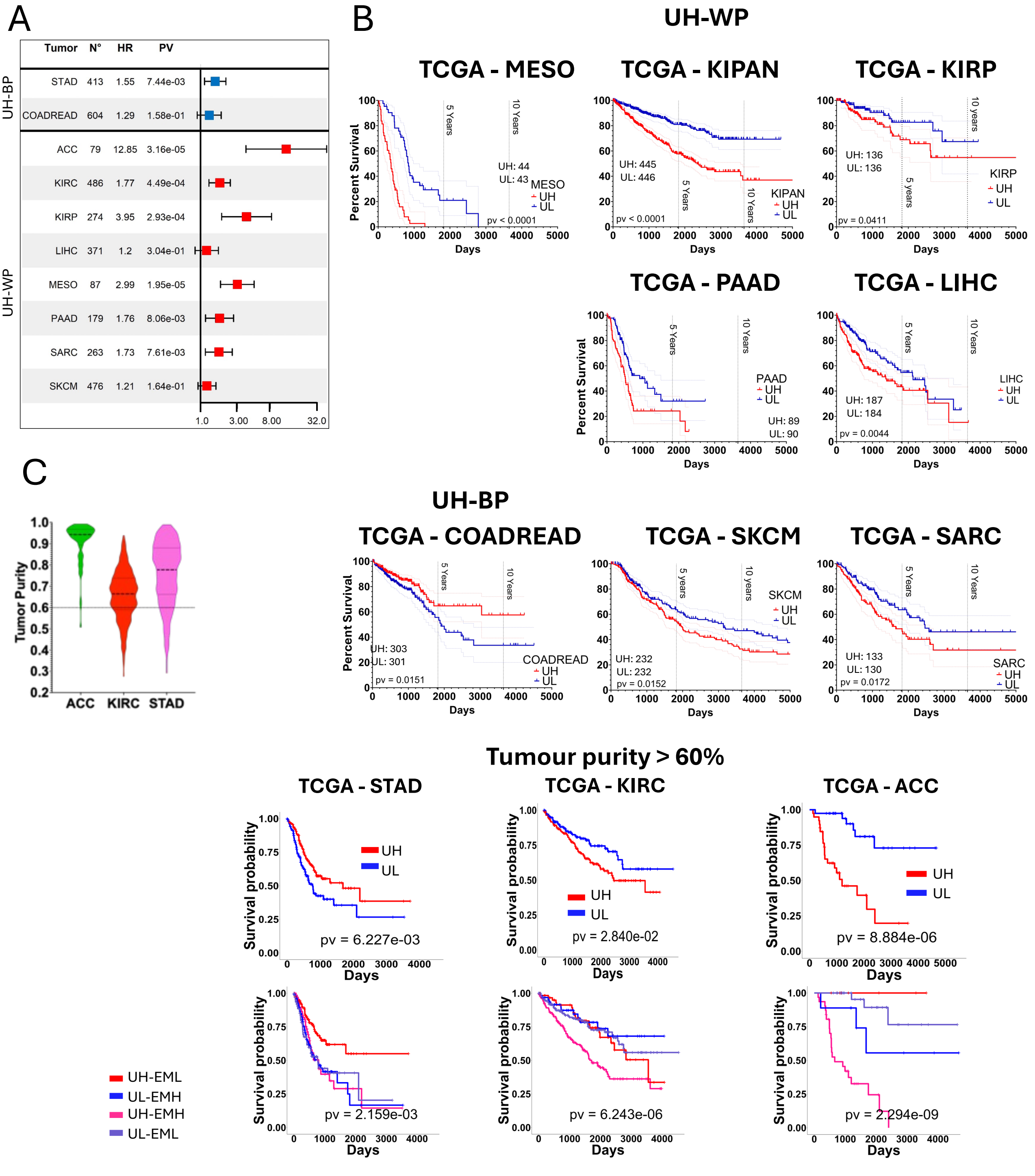


**Figure S1. UHRF1 expression has a relevant prognostic impact which is maintained after adjustment for tumour-purity.** A) Hazard ratio (HR) from survival analysis comparing high (EMH) with low (EML) Embryonic Morphogenesis expression in TCGA dataset (blue squares: UH-BP tumours; red squares: UH-WP tumours); B) UHRF1 patient overall survival UHRF1-*high* (RED), UHRF1-*low* (BLUE) (log-rank *Mantel-Cox* p value); C) Left: violin plot of the tumour purity distribution in ACC, KIRC and STAD. Right: UHRF1 and EM patient survival filtered for >60% tumour purity.

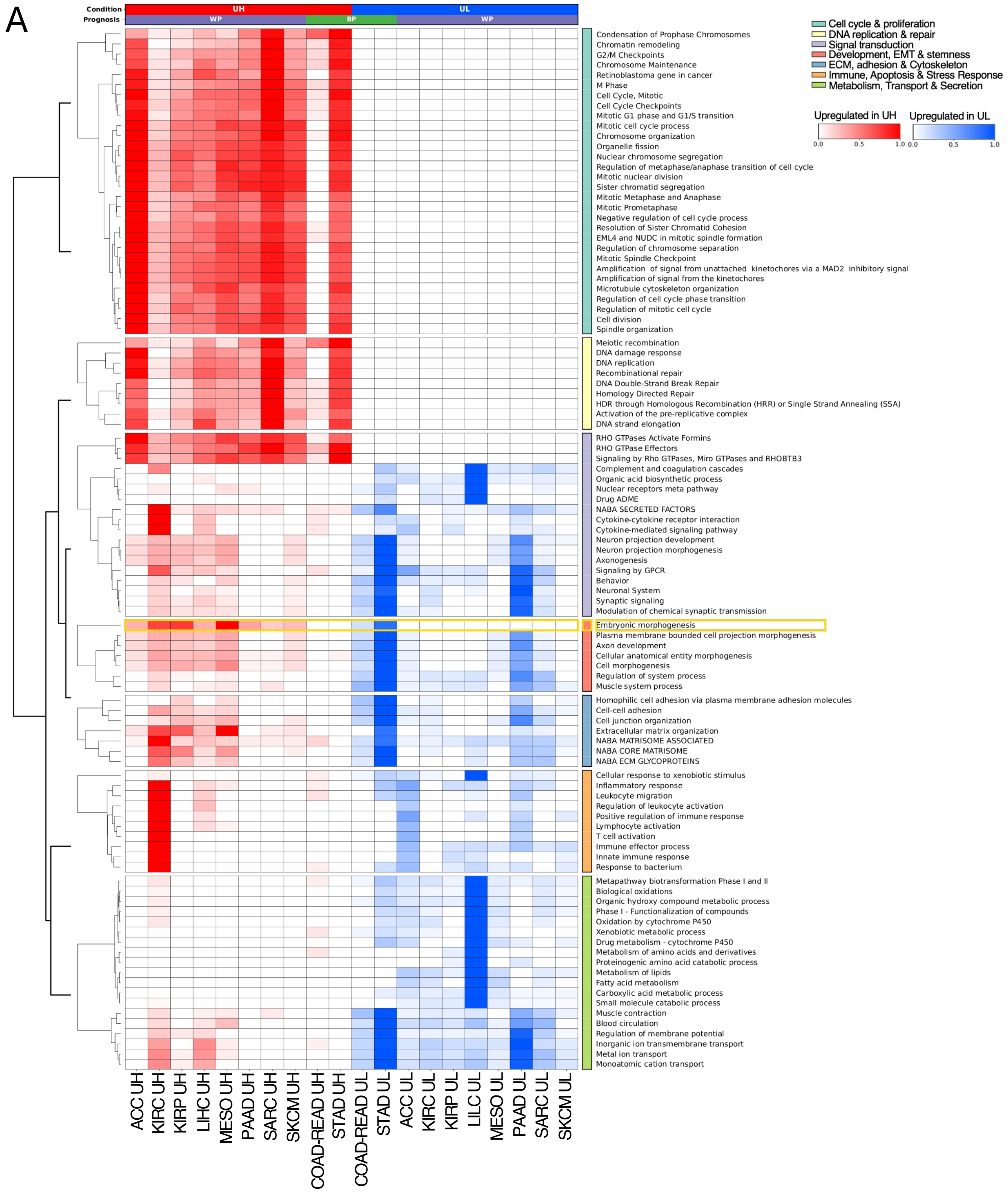


**Figure S2**. **Embryonic Morphogenesis is oppositely correlated to UHRF1 expression in BP and WP**. A) heatmap of top 100 enriched Metascape pathways clustered in macro-categories associated with cell cycle, DNA repair, signal transduction, metabolism and cell transport, development and stemness, including *Embryonic Morphogenesis*; columns: tumour-specific *DE-UH* gene list (Red); tumour-specific *DE-UL gene lists (Blue).* Colour code for ontologies enrichment: Red gradient (enrichment score 0-1, see M&M), enrichment in DE-UH gene list; Blue gradient (enrichment score 0-1, see M&M): enrichment in DE-UL gene list.

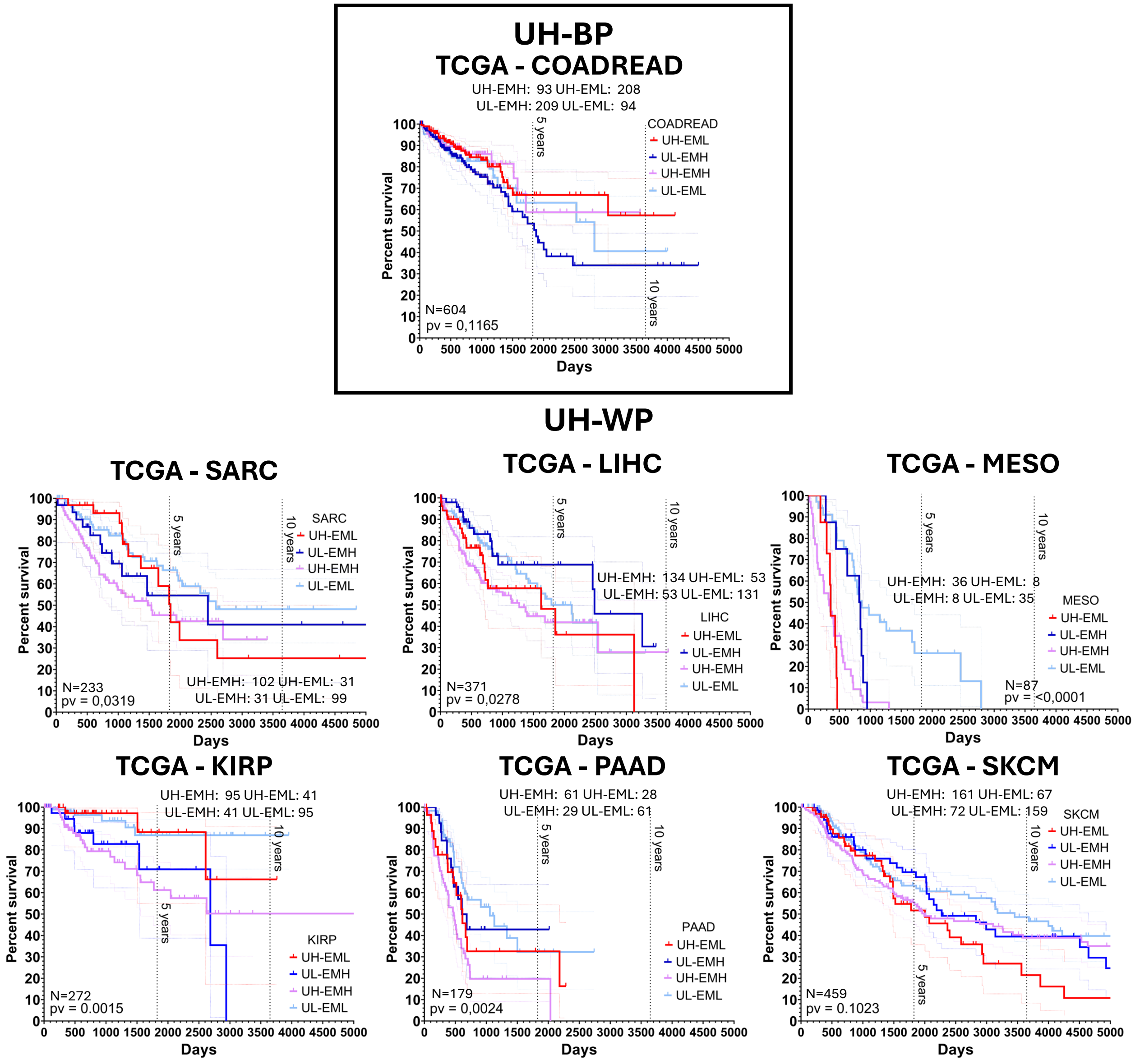


**Figure S3. Combined UHRF1 and embryonic morphogenesis expression enable prognostic stratification in a selection of TCGA tumours.** Differences in TCGA-patient overall survival within UHRF1-significant tumours dependent upon UHRF1-EM categories (log-rank *Mantel-Cox* p value). The EM signature pertains to genes associated with worst prognosis: UL associated genes for UH-BP tumours; UH associated genes for UH-WP tumours.

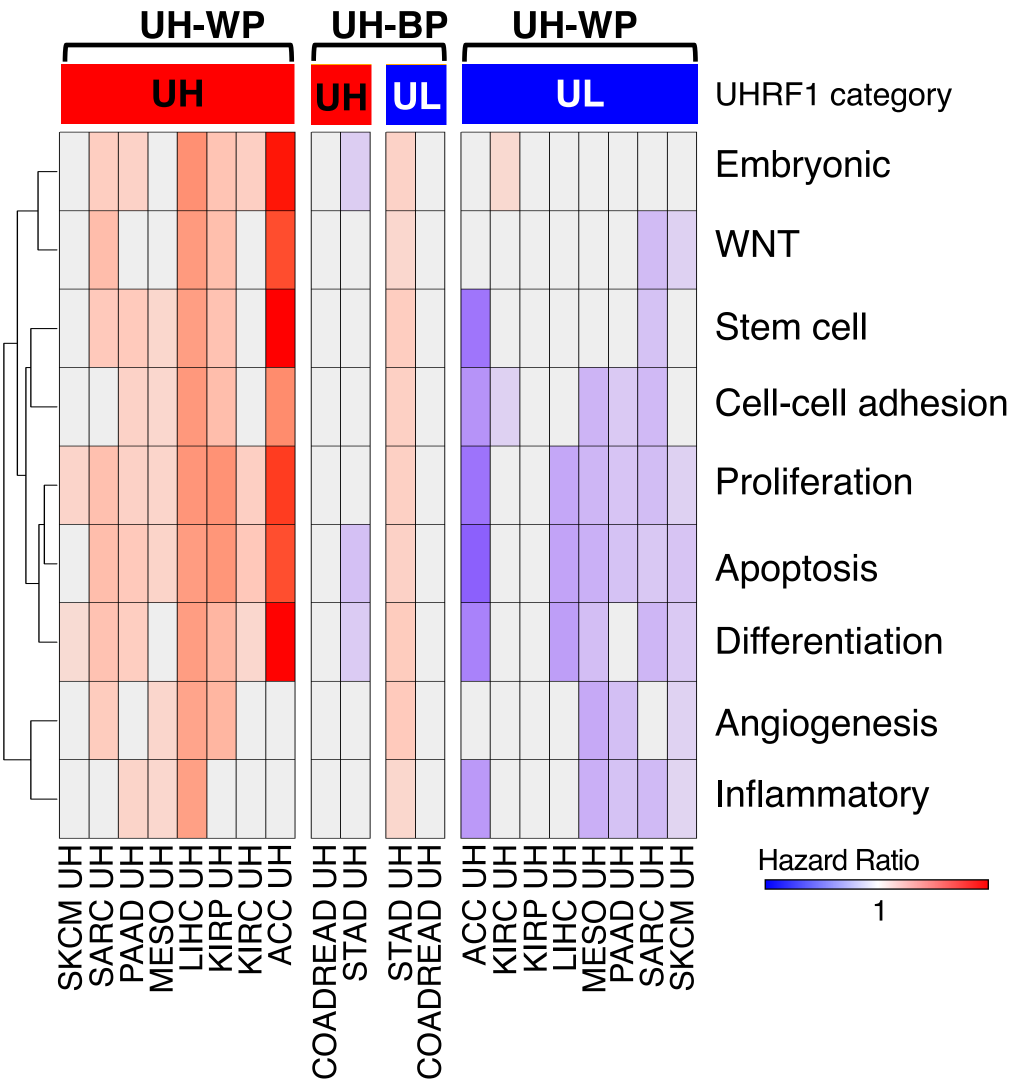


**Figure S4**. **Expression of cumulative ontologies discriminates prognostic outcome in a selection of TCGA tumours.** Hazard ratio (HR) heatmap for high versus low gene expression of cumulative ontologies of DE-UH and DE-UL gene lists. Red gradient: HR >1 worst prognosis; Bleu gradient: HR<1 best prognosis; Grey: HR not statistically significant.

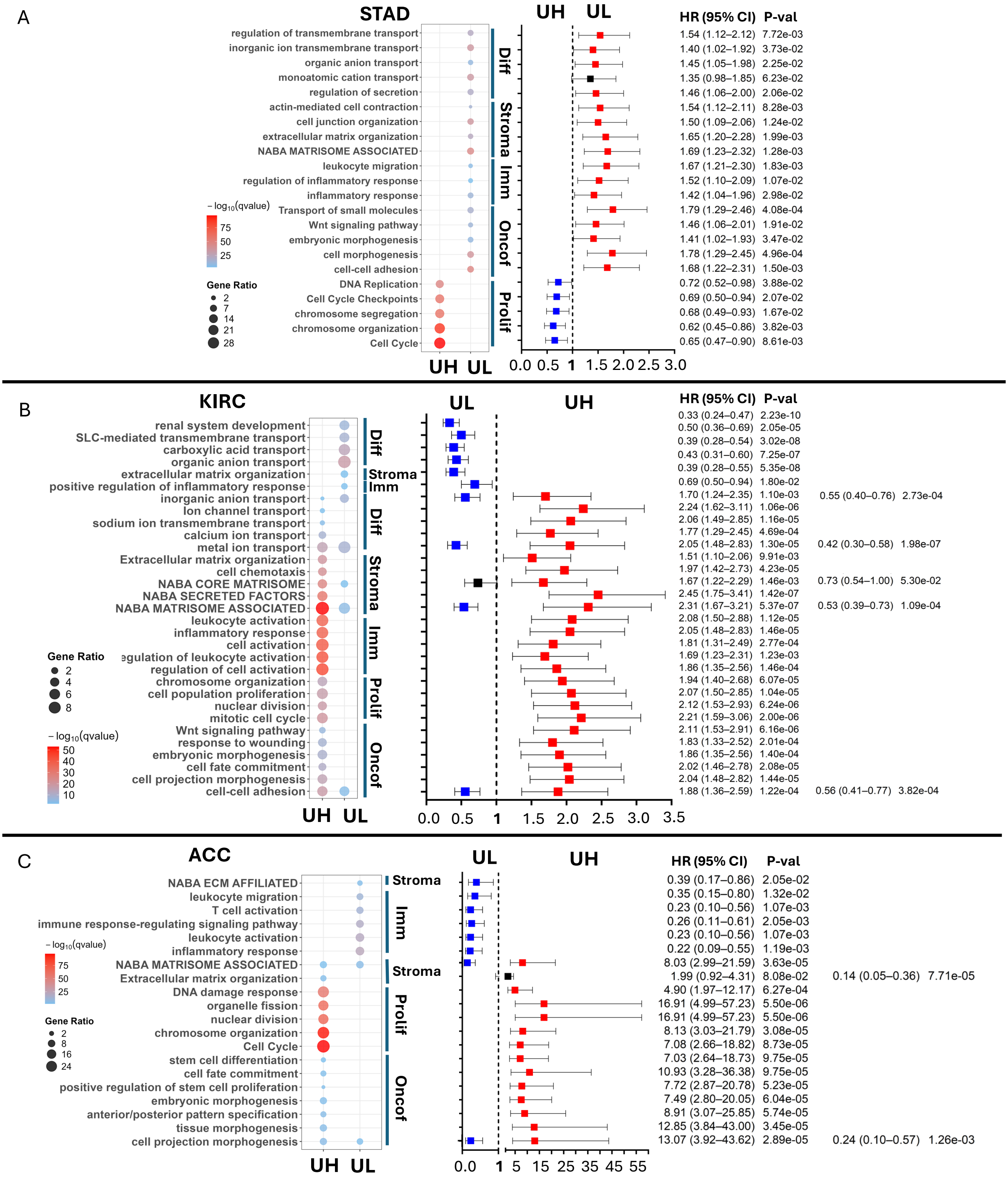


**Figure S5**. **Proliferative traits are consistently correlated with UHRF1 expression, while correlation of onco-foetal traits is tumour-specific in STAD, KIRC and ACC**. STAD A), KIRC B), ACC C): Dot Plots: selection of most significantly enriched pathways in UH and UL gene lists; statistical significance of the enrichment represented as –log10(q-value)(light blue to red gradient) and % of UH and UL genes enriched in each ontology reported as Gene Ratio; Forest Plot: hazard ratio (HR) from survival analysis, comparing UH genes with UL genes in TCGA dataset. Horizontal lines stand for 95% confidence intervals for the ratios.

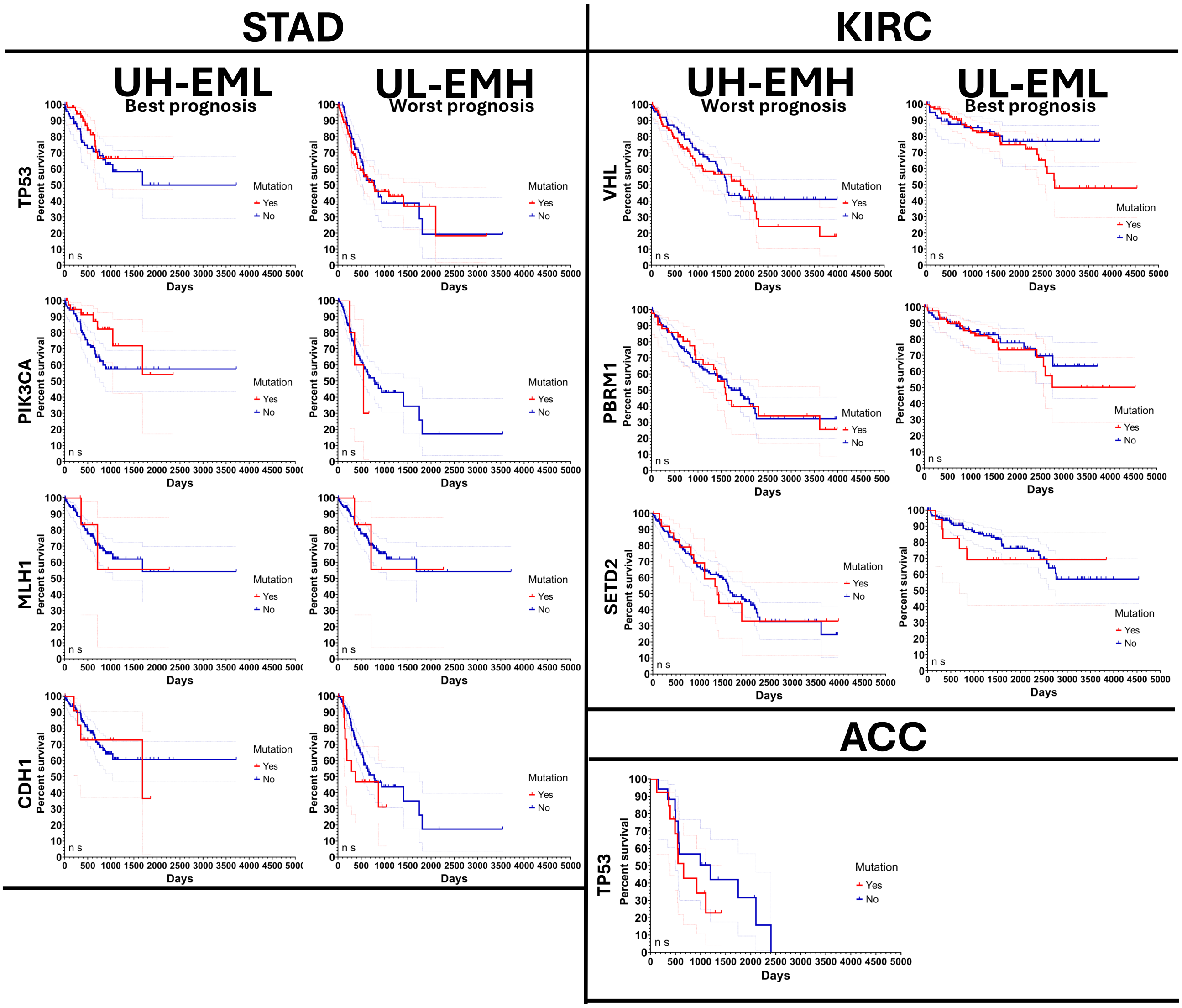


**Figure S6. Mutation of driver onco-genes does not associate with prognosis across UHRF1-EM categories**. Differences in overall Survival within UHRF1-EM best and worst prognosis categories based on key genes' mutational state. STAD - UH-EML and UL-EMH (TP53, PIK3CA, MLH1, CDH1); KIRC – UL-EML and UH-EMH (VHL, PBRM1, SETD2); ACC – UL-EML and UH-EMH (TP53).

A

TCGA - STAD

| Charact. | UH EMH | | UH EML | | UL EMH | | UL EML | | Pv |
|---|---|---|---|---|---|---|---|---|---|
| **Sex** | | | | | | | | | |
| Female | 28 | 39.44% | 51 | 37.50% | 49 | 36.03% | 18 | 25.71% | ns |
| Male | 43 | 60.56% | 85 | 62.50% | 87 | 63.97% | 52 | 74.29% | |
| **pT stage** | | | | | | | | | |
| T1 + T2 | 17 | 23.94% | 38 | 27.94% | 31 | 22.79% | 24 | 34.29% | ns |
| T3 + T4 | 54 | 76.06% | 97 | 71.32% | 100 | 73.53% | 43 | 61.43% | |
| Unknown | 0 | 0.00% | 1 | 0.74% | 5 | 3.68% | 3 | 4.29% | |
| **pM Stage** | | | | | | | | | |
| M0 | 62 | 87.32% | 125 | 91.91% | 117 | 86.03% | 62 | 88.57% | ns |
| M1 | 4 | 5.63% | 6 | 4.41% | 11 | 8.09% | 5 | 7.14% | |
| MX | 5 | 7.04% | 5 | 3.68% | 8 | 5.88% | 3 | 4.29% | |
| Unknown | | | | | | | | | |
| **pN stage** | | | | | | | | | |
| Discr. | 1 | 1.41% | 0 | 0.00% | 0 | 0.00% | 0 | 0.00% | ns |
| N0 | 18 | 25.35% | 53 | 38.97% | 36 | 26.47% | 16 | 22.86% | |
| N1 + N2 + N3 | 51 | 71.83% | 82 | 60.29% | 91 | 66.91% | 48 | 68.57% | |
| NX | 1 | 1.41% | 1 | 0.74% | 9 | 6.62% | 6 | 8.57% | |
| **pStage** | | | | | | | | | |
| Discrepancy | 2 | 2.82% | 1 | 0.74% | 2 | 1.47% | 1 | 1.43% | ns |
| I + II | 25 | 35.21% | 72 | 52.94% | 56 | 41.18% | 27 | 38.57% | |
| III + IV | 43 | 60.56% | 61 | 44.85% | 69 | 50.74% | 37 | 52.86% | |
| Unknown | 1 | 1.41% | 2 | 1.47% | 9 | 6.62% | 5 | 7.14% | |
| **P.T.O** | | | | | | | | | |
| Discr. | 0 | 0.00% | 1 | 0.74% | 1 | 0.74% | 2 | 2.86% | ns |
| Compl. remission | 44 | 61.97% | 85 | 62.50% | 75 | 55.15% | 35 | 50.00% | |
| Part. remission | 3 | 4.23% | 1 | 0.74% | 4 | 2.94% | | 0.00% | |
| Progr. disease | 17 | 23.94% | 18 | 13.24% | 24 | 17.65% | 9 | 12.86% | |
| Stable disease | 2 | 2.82% | 12 | 8.82% | 7 | 5.15% | 6 | 8.57% | |
| Unknown | 5 | 7.04% | 19 | 13.97% | 25 | 18.38% | 18 | 25.71% | |
| **Mol. Status** | | | | | | | | | |
| MSI-High + Low | 18 | 25.35% | 76 | 55.88% | 24 | 17.65% | 20 | 28.57% | <0.0001 |
| MSS | 53 | 74.65% | 59 | 43.38% | 112 | 82.35% | 50 | 71.43% | **** |
| Unknown | 0 | 0.00% | 1 | 0.74% | 0 | 0.00% | 0 | 0.00% | |
| **Mol. Sub** | | | | | | | | | |
| CIN | 46 | 64.79% | 47 | 34.56% | 80 | 58.82% | 48 | 68.57% | <0.0001 |
| EBV | 6 | 8.45% | 18 | 13.24% | 2 | 1.47% | 4 | 5.71% | **** |
| GS | 4 | 5.63% | 4 | 2.94% | 39 | 28.68% | 3 | 4.29% | |
| HM-SNV | 2 | 2.82% | 3 | 2.21% | 1 | 0.74% | 1 | 1.43% | |
| MSI | 4 | 5.63% | 52 | 38.24% | 8 | 5.88% | 9 | 12.86% | |
| Unknown | 9 | 12.68% | 12 | 8.82% | 6 | 4.41% | 5 | 7.14% | |
| **N.tot** | 71 | | 136 | | 136 | | 70 | | 413 |

TCGA - KIRC

| Charact. | UH EMH | | UH EML | | UL EMH | | UL EML | | Pv |
|---|---|---|---|---|---|---|---|---|---|
| **Sex** | | | | | | | | | |
| Female | 51 | 28.98% | 29 | 43.28% | 25 | 37.31% | 71 | 40.57% | ns |
| Male | 125 | 71.02% | 38 | 56.72% | 42 | 62.69% | 104 | 59.43% | |
| **pT stage** | | | | | | | | | |
| T1 + T2 | 82 | 46.59% | 47 | 70.15% | 49 | 73.13% | 129 | 73.71% | <0.0001 |
| T3 + T4 | 94 | 53.41% | 20 | 29.85% | 18 | 26.87% | 46 | 26.29% | **** |
| **pM Stage** | | | | | | | | | |
| M0 | 128 | 72.73% | 51 | 76.12% | 52 | 77.61% | 148 | 84.57% | 0.0171 |
| M1 | 41 | 23.30% | 12 | 17.91% | 7 | 10.45% | 14 | 8.00% | * |
| MX | 7 | 3.98% | 3 | 4.48% | 7 | 10.45% | 13 | 7.43% | |
| Unknown | 0 | 0.00% | 1 | 1.49% | 1 | 1.49% | 0 | 0.00% | |
| **pN stage** | | | | | | | | | |
| N0 | 85 | 48.30% | 25 | 37.31% | 30 | 44.78% | 76 | 43.43% | 0.0026 |
| N1 + N2 + N3 | 12 | 6.82% | 0 | 0.00% | 0 | 0.00% | 1 | 0.57% | ** |
| NX | 79 | 44.89% | 42 | 62.69% | 37 | 55.22% | 98 | 56.00% | |
| **pStage** | | | | | | | | | |
| I + II | 76 | 43.18% | 46 | 68.66% | 45 | 67.16% | 123 | 70.29% | <0.0001 |
| III + IV | 100 | 56.82% | 21 | 31.34% | 22 | 32.84% | 52 | 29.71% | **** |
| **N. tot** | 176 | | 67 | | 67 | | 175 | | 485 |

TCGA - ACC

| Charact. | UH-EMH | | UH-EML | | UL-EMH | | UL-EML | | Pv |
|---|---|---|---|---|---|---|---|---|---|
| **Sex** | | | | | | | | | |
| Female | 20 | 64.52% | 3 | 33.33% | 7 | 77.78% | 18 | 60.00% | ns |
| Male | 11 | 35.48% | 6 | 66.67% | 2 | 22.22% | 12 | 40.00% | |
| **pT stage** | | | | | | | | | |
| T1 + T2* | 12 | 38.71% | 7 | 77.78% | 7 | 77.78% | 25 | 83.33% | 0.0046 |
| T3 + T4 | 17 | 54.84% | 2 | 22.22% | 2 | 22.22% | 5 | 16.67% | ** |
| Unknown | 2 | 6.45% | 0 | 0.00% | 0 | 0.00% | 0 | 0.00% | |
| **pN stage** | | | | | | | | | |
| N0 | 23 | 74.19% | 9 | 100.00% | 6 | 66.67% | 30 | 100.00% | 0.0092 |
| N1 + N2 + N3 | 6 | 19.35% | 0 | 0.00% | 3 | 33.33% | 0 | 0.00% | ** |
| Unknown | 2 | 6.45% | 0 | 0.00% | 0 | 0.00% | 0 | 0.00% | |
| **pStage** | | | | | | | | | |
| I + II* | 10 | 32.26% | 7 | 77.78% | 4 | 44.44% | 25 | 83.33% | 0.0008 |
| III + IV | 19 | 61.29% | 2 | 22.22% | 5 | 55.56% | 5 | 16.67% | *** |
| Unknown | 2 | 6.45% | 0 | 0.00% | 0 | 0.00% | 0 | 0.00% | |
| **P.T.O** | | | | | | | | | |
| Compl. remission | 9 | 29.03% | 8 | 88.89% | 5 | 55.56% | 25 | 83.33% | 0.0022 |
| Part. remission | 0 | 0 | 0 | 0 | 0 | 0 | 1 | 3.33% | ** |
| Progr. disease | 12 | 38.71% | 1 | 11.11% | 2 | 22.22% | 2 | 6.67% | |
| Stable disease | 0 | 0.00% | 0 | 0.00% | 1 | 11.11% | 1 | 3.33% | |
| Unknown | 10 | 32.26% | 0 | 0.00% | 1 | 11.11% | 1 | 3.33% | |
| **N. tot** | 31 | | 9 | | 9 | | 30 | | 79 |

B

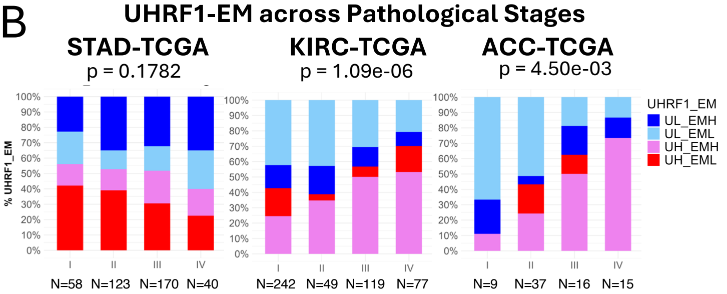


C

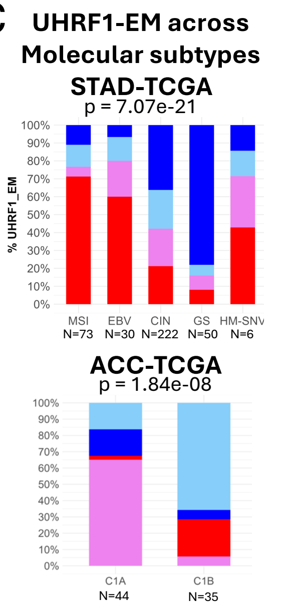


D

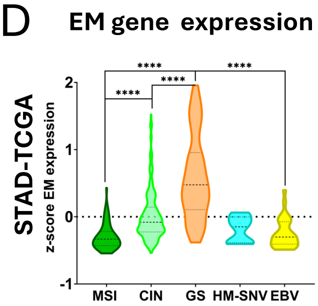


E

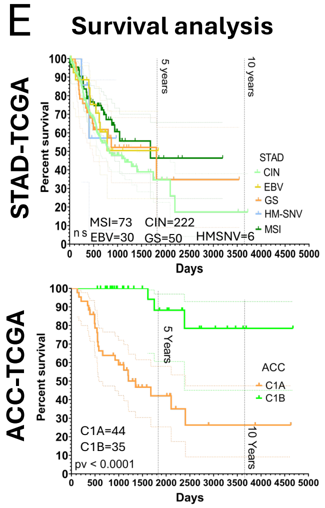


**Figure S7. UHRF1-EM categories intercepts clinical and molecular features which are coherent with prognosis in TCGA.** A) *Clinical tables*: frequency distribution of UHRF1-EM classified individuals across clinical and pathological features of tumours in TCGA datasets (Chi-square test p-value); B) Bar-charts: distribution of UHRF1-EM categories across pathological stages and C) molecular subtypes; D) *Violin plots*: EM gene expression distribution across TCGA-STAD molecular groups (Kruskal-Wallis and Dunn test p value); E) *Kaplan-Maier:* differences in patients OS depending upon STAD-TCGA and ACC-TCGA defined molecular subtypes (log-rank Mantel-Cox p value).

A

| ACRG | | | | | | | | | |
|---|---|---|---|---|---|---|---|---|---|
| Charact. | UH EMH | | UH EML | | UL EMH | | UL EML | | Pv |
| **Sex** | | | | | | | | | |
| Female | 12 | 24.00% | 28 | 28.00% | 44 | 44.00% | 17 | 34.00% | 0.04 |
| Male | 38 | 76.00% | 72 | 72.00% | 56 | 56.00% | 33 | 66.00% | * |
| **pT stage** | | | | | | | | | |
| T1 + T2 | 32 | 64.00% | 78 | 78.00% | 42 | 42.00% | 34 | 68.00% | <0.0001 |
| T3 + T4 | 18 | 36.00% | 20 | 20.00% | 58 | 58.00% | 16 | 32.00% | **** |
| TX | 0 | 0.00% | 2 | 2.00% | 0 | 0.00% | 0 | 0.00% | |
| **pM Stage** | | | | | | | | | |
| M0 | 46 | 92.00% | 95 | 95.00% | 88 | 88.00% | 44 | 88.00% | ns |
| M1 | 4 | 8.00% | 5 | 5.00% | 12 | 12.00% | 6 | 12.00% | |
| **pN stage** | | | | | | | | | |
| N0 | 5 | 10.00% | 22 | 22.00% | 7 | 7.00% | 4 | 8.00% | 0.0071 |
| N1 + N2 + N3 | 45 | 90.00% | 78 | 78.00% | 93 | 93.00% | 46 | 92.00% | ** |
| **pStage** | | | | | | | | | |
| I + II | 24 | 48.00% | 56 | 56.00% | 21 | 21.00% | 25 | 50.00% | <0.0001 |
| III + IV | 26 | 52.00% | 42 | 42.00% | 79 | 79.00% | 25 | 50.00% | **** |
| Unknown | 0 | 0.00% | 2 | 4.00% | 0 | 0.00% | 0 | 0.00% | |
| **Doc. relapse** | | | | | | | | | |
| No | 26 | 52.00% | 66 | 66.00% | 37 | 37.00% | 28 | 56.00% | 0.0003 |
| yes | 21 | 42.00% | 27 | 27.00% | 58 | 58.00% | 19 | 38.00% | *** |
| Unknown | 3 | 6.00% | 7 | 7.00% | 5 | 5.00% | 3 | 6.00% | |
| **Mol. sub.** | | | | | | | | | |
| EMT | 12 | 24.00% | 0 | 0.00% | 34 | 34.00% | 0 | 0.00% | <0.0001 |
| MSI | 9 | 18.00% | 52 | 52.00% | 3 | 3.00% | 4 | 8.00% | **** |
| MSS/TP53- | 15 | 30.00% | 26 | 26.00% | 38 | 38.00% | 28 | 56.00% | |
| MSS/TP53+ | 14 | 28.00% | 22 | 22.00% | 25 | 25.00% | 18 | 36.00% | |
| **N. tot** | **50** | | **100** | | **100** | | **50** | | **300** |

| ccRCC - E - MTAB1980 | | | | | | | | | |
|---|---|---|---|---|---|---|---|---|---|
| Charact. | UH EMH | | UH EML | | UL EMH | | UL EML | | Pv |
| **pT stage** | | | | | | | | | |
| T1 + T2 | 21 | 65.62% | 17 | 89.47% | 14 | 77.78% | 27 | 84.38% | ns |
| T3 + T4 | 11 | 34.38% | 2 | 10.53% | 4 | 22.22% | 5 | 15.63% | |
| **pM Stage** | | | | | | | | | |
| M0 | 27 | 84.38% | 18 | 94.74% | 17 | 94.44% | 27 | 84.38% | ns |
| M1 | 5 | 15.63% | 1 | 5.26% | 1 | 5.56% | 5 | 15.63% | |
| **pN stage** | | | | | | | | | |
| N0 | 27 | 84.38% | 18 | 94.74% | 18 | 100.00% | 31 | 96.88% | ns |
| N1 + N2 + N3 | 5 | 15.63% | 1 | 5.26% | 0 | 0.00% | 1 | 3.13% | |
| **pStage** | | | | | | | | | |
| I + II | 20 | 62.50% | 16 | 84.21% | 14 | 77.77% | 25 | 78.12% | ns |
| III + IV | 12 | 37.50% | 3 | 15.78% | 4 | 22.22% | 6 | 18.75% | |
| Unknown | 0 | 0.00% | 0 | 5.26% | 0 | 0.00% | 1 | 3.13% | |
| **N. tot** | **32** | | **19** | | **18** | | **32** | | **101** |

| ACC - 2 | | | | | | | | | |
|---|---|---|---|---|---|---|---|---|---|
| Charact. | UH EMH | | UH EML | | UL EMH | | UL EML | | Pv |
| **Weiss score** | | | | | | | | | |
| W0-W3 | 1 | 2.22% | 2 | 13.33% | 11 | 55.00% | 35 | 81.40% | <0.0001 |
| W4-W9 | 38 | 84.44% | 13 | 86.67% | 9 | 45.00% | 8 | 18.60% | **** |
| Unknown | 6 | 13,33% | 0 | 0,00% | 0 | 0,00% | 0 | 0,00% | |
| **ENSAT Stage** | | | | | | | | | |
| 1 | 3 | 6.67% | 3 | 20.00% | 11 | 55.00% | 27 | 62.79% | <0.0001 |
| 2 | 20 | 44.44% | 11 | 73.33% | 6 | 30.00% | 11 | 25.58% | **** |
| 3 | 14 | 31.11% | 1 | 6.67% | 2 | 10.00% | 4 | 9.30% | |
| 4 | 8 | 17.78% | 0 | 0.00% | 1 | 5.00% | 1 | 2.33% | |
| **Diagnosis** | | | | | | | | | |
| ACA | 1 | 2.22% | 2 | 13.33% | 9 | 45.00% | 29 | 67.44% | <0.0001 |
| ACT | 44 | 97.78% | 11 | 73.33% | 11 | 55.00% | 13 | 30.23% | **** |
| potential ACT | 0 | 0.00% | 2 | 13.33% | 0 | 0.00% | 1 | 2.33% | |
| **N. tot** | **45** | | **15** | | **20** | | **43** | | **123** |

B

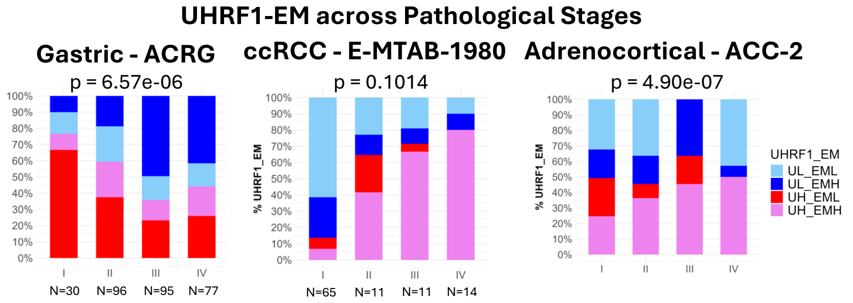


C

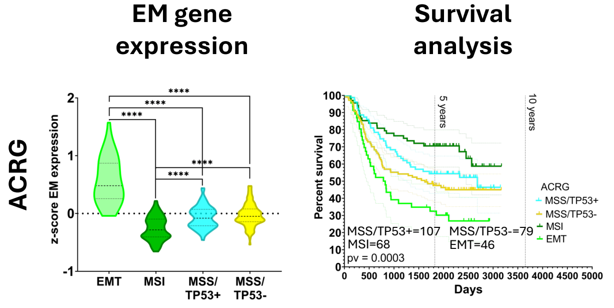


D

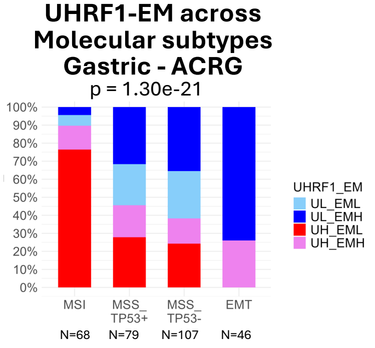


**Figure S8. UHRF1-EM categories intercepts clinical and molecular features which are coherent with prognosis in validation datasets** A) *Clinical tables*: frequency distribution of UHRF1-EM classified individuals across clinical and pathological features of tumours in secondary validation datasets (Chi-square test p-value); B) *Bar-charts*: distribution of UHRF1-EM categories across pathological stages and D) molecular subtypes; C) *Violin plots*: EM gene expression distribution across ACRG molecular groups (Kruskal-Wallis and Dunn test p value); *Kaplan-Maier:* differences in patients OS depending upon ACRG defined molecular groups (log-rank Mantel-Cox p value).

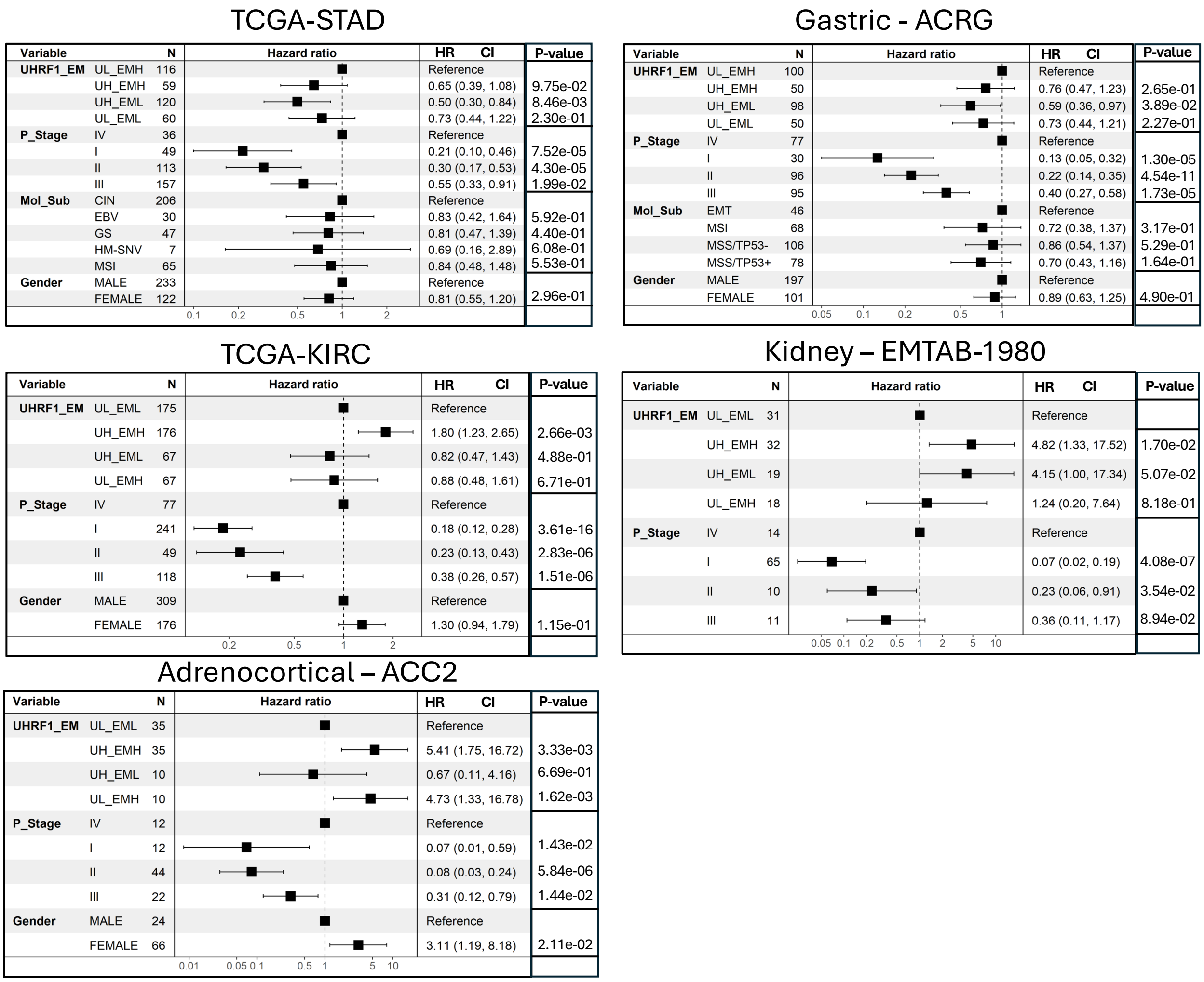


**Figure S9. Multivariate Cox analysis: UHRF1-EM categories are an independent prognostic factor after adjustment for pathological and molecular variables.** Hazard Ratio (HR) from Multivariate Cox Regression analysis of UHRF1-EM categories adjusted for other covariates including Gender, P_Stage (pathological stage), Mol_Sub (molecular subtypes), when available. HR: Hazard Ratio; CI: Confidence Interval of hazard ratio; N: number of patients for each level of the variable.

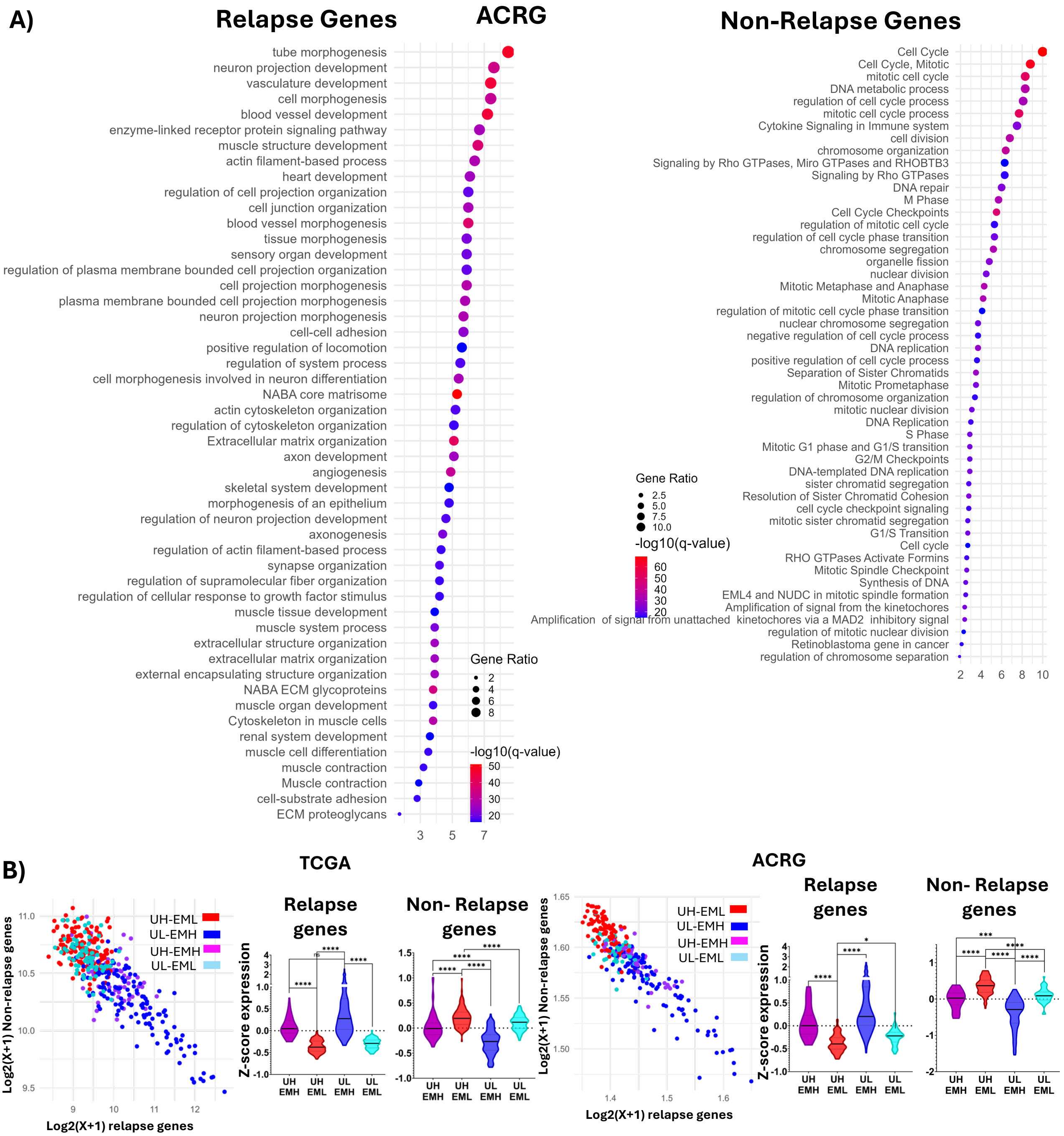


**Figure S10. Relapse and Non-Relapse associated genes are enriched for embryonic and proliferative terms and their expression impacts patients' prognosis.** A) Dot Plots: Top 70 P-Value ranked enriched pathways in Relapse (left panel) and Non-relapse (right panel) genes in ACRG gastric cancer dataset; statistical significance of the enrichment represented as –log10(q-value); B) Scatterplot: Log2(X+1) expression of Relapse (x-axis) and Non-Relapse (y-axis) signature across UHRF1-EM patients in TCGA (left) and ACRG (right); Violin Plots: Z-score expression distribution of Relapse-embryonic (Rel. genes) and Non-Relapse-proliferation genes (Non-Rel. genes) across UHRF1-EM categories in TCGA (left panel) and ACRG (right panel). Kruskal-Wallis Dunn's Test statistics. ****=p value<0.0001; ***=p value ≤0.001; **=p value ≤0.01; *=p value ≤ 0.05

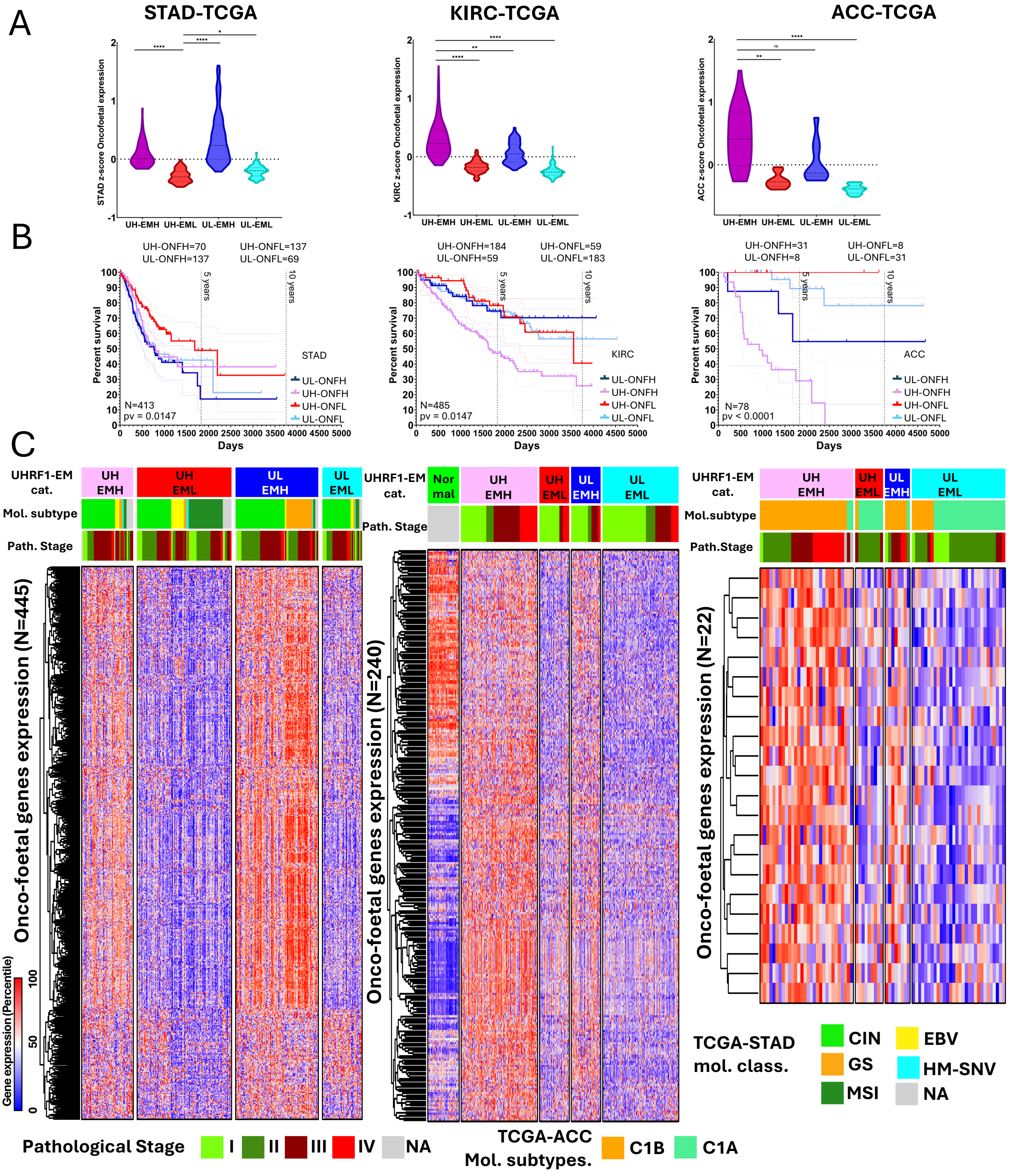


**Figure S11. Oncofoetal genes expression and survival analysis**. A) Violin plots of OnF expression levels across UHRF1-EM categories in TCGA-STAD, KIRC and ACC. ****=p value<0.0001; ***=p value ≤0.001; **=p value ≤0.01; *=p value ≤0.05 (Kruskal Wallis Dunn post test p value). B) KM: Patients' overall survival across UHRF1-ONF categories (log-rank Mantel-Cox p value); C) Heatmap representation of WP-associated OnF genes expression profile across UHRF1-EM categories in TCGA-STAD, KIRC and ACC. When available, pathological stages and molecular subtypes are indicated. Gene expression percentile varies according to a red (high expression) to blue (low expression) gradient.

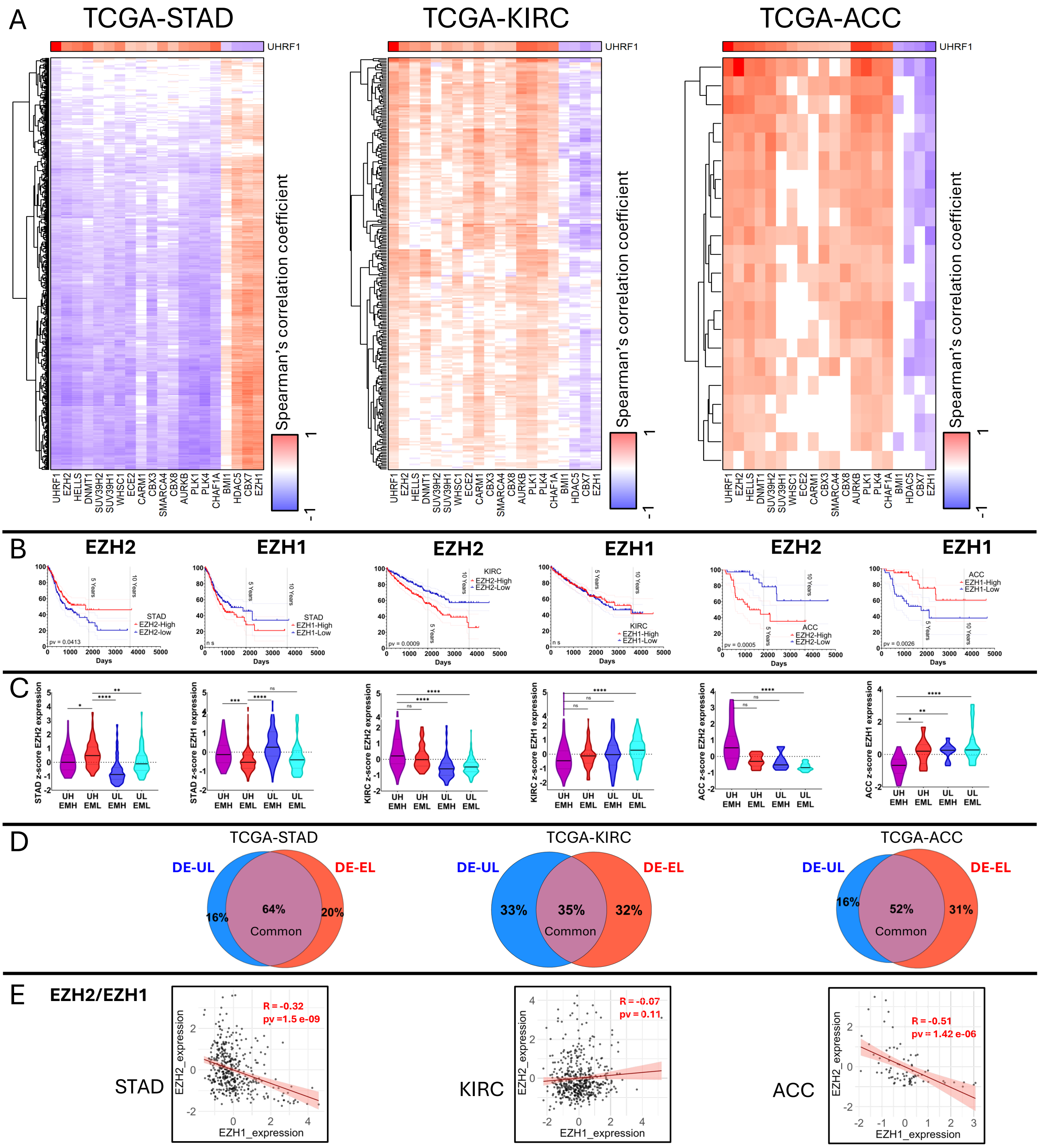


**Figure S12. Tumour-specific UHRF1-centred epigenetic factors oppositely correlate with oncofoetal programmes across cancer types. A)** heatmap reporting the Spearman coefficient of the correlation between epigenetic effectors (columns) and *onco-foetal* genes (rows). The coefficient varies from -1 (blue, anti-correlation) to 1 (red, correlation); significant correlations were defined by a Spearman R value of ≥0.2 or ≤-0.2, with a p-value <0.0001. B) survival analysis dependent upon EZH1 and EZH2 expression in TCGA-STAD, KIRC and ACC (Log Ran Mantel-Cox test*; red curve: high expression, blue curve: low expression*); C) violin plots: EZH1 and EZH2 expression across UHRF1-EM categories in TCGA-STAD, KIRC and ACC (Kruskal Wallis Dunn post test); D) Venn-Diagrams showing intersection between DE-WP genes calculated according to UHRF1 and EZH2 high and low expression. DE-UL and DE-EL: DE for UHRF1 and EZH2 expression, respectively; E) scatterplot showing the correlation between EZH1 and EZH2 expression in the three tumours (Spearman correlation test).

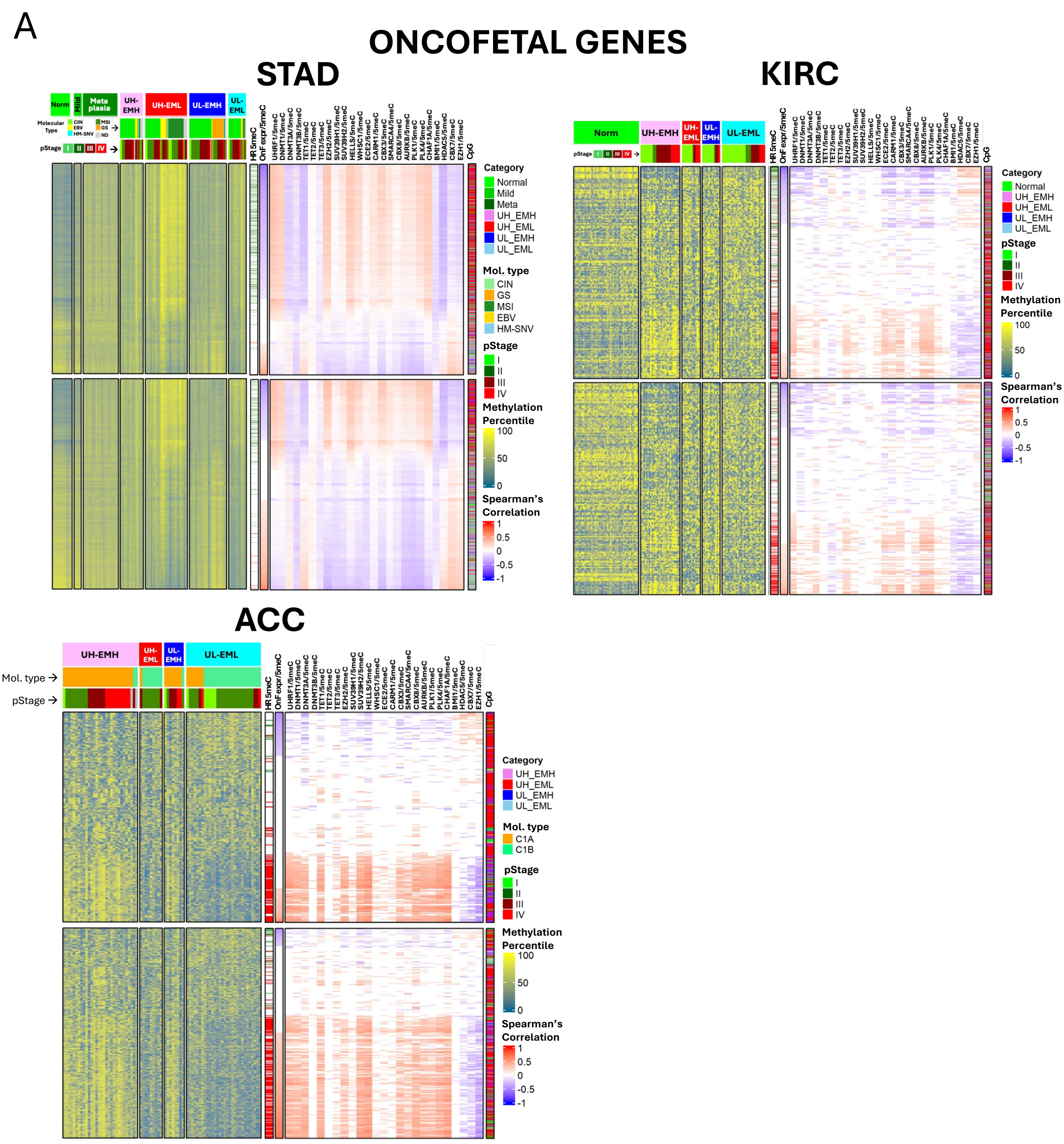


**Figure S13. UHRF1-associated DNA methylation at bivalent sites regulates EM gene expression and influences prognosis.** Heatmap representation of cytosines methylations of OnF genes across the four UHRF1-EM categories in A) TCGA-STAD, B) TCGA-KIRC, C) TCGA-ACC. DNA methylation varies according to a yellow (high methylation) to blue (low methylation) gradient. UHRF1-EM categories, TCGA molecular subtypes and tumour pathological stage are indicated above the heatmap; rows: CpGs; columns: patients; *OnF expr UX-EMX*: expression of OnF gene associated to the correspondent; *CpGi:* genomic context of the reported cytosines (blue: s-shore, light blue: s-shelf, red: island, light green: n-shelf, dark green: n-shore, gray: non-CpG); *HR 5meC*: hazard ratio for high vs low methylation level of the cytosine (green: HR<1, better prognosis, red: HR>1, worse prognosis)*;* *OnF expression/5meC:* correlation between OnF gene expression and Cpg methylation; *UHRF1/5meC:* correlation between the expression of the indicated epigenetic effector (UHRF1, DNMT1…) and CpGs' methylation (Spearman's correlation test, blue: anti-correlation; red: correlation; white: no statistically significant)

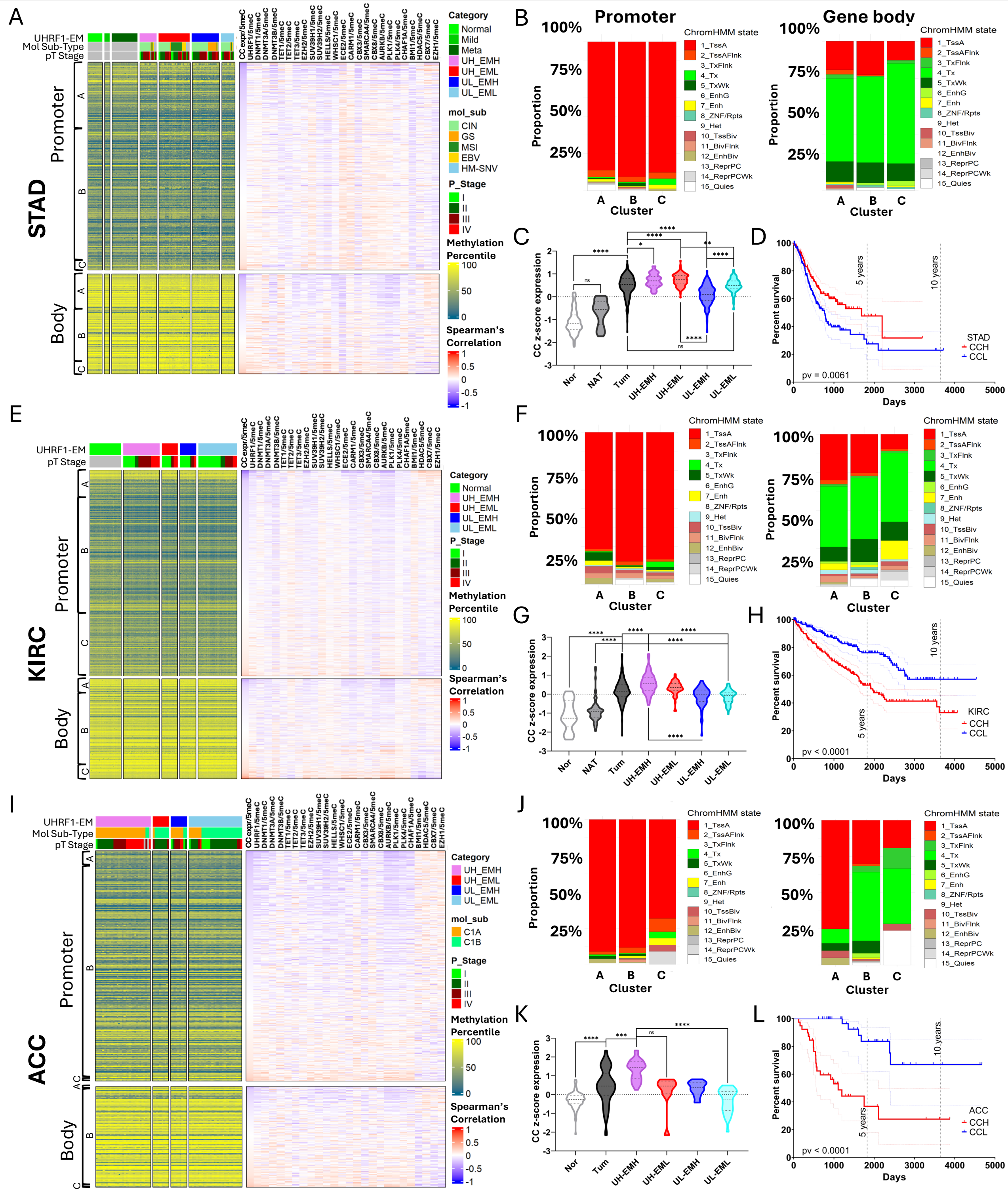

**Figure S14. DNA methylation of Cell Cycle (CC) genes does not vary across UHRF1-EM categories, but CC genes expression is condition-specific.** Heatmap representation of cytosines methylations of CC genes across the four UHRF1-EM categories in A) TCGA-STAD, E) TCGA-KIRC, I) TCGA-ACC. DNA methylation varies according to a yellow (high methylation) to blue (low methylation) gradient. UHRF1-EM categories, TCGA molecular subtypes and tumour pathological stage are indicated. rows: CC CpGs; columns: patients. *CC expression/5meC:* correlation between the CC gene expression and CpG methylation; *UHRF1/5meC:* correlation between the expression of the indicated epigenetic effector (UHRF1, DNMT1…) and CpG methylation (Spearman's correlation test, blue: anti-correlation; red: correlation). B-F-J) Frequency distribution of CC cytosines across Epigenetic Roadmap genomic region annotations (see list of abbreviations); X-axis: CpG sites clusters; y-axis: percentage of cytosines located in annotated genomic regions (left histogram: Promoter, right histogram: Body); C-J-K) CC genes expression (z-score) across UHRF1-EM categories, normal (Nor) and normal adjacent tumour (NAT) tissue; D-H-L) KM survival curve for high vs low CC signature expression in STAD, KIRC and ACC, respectively.

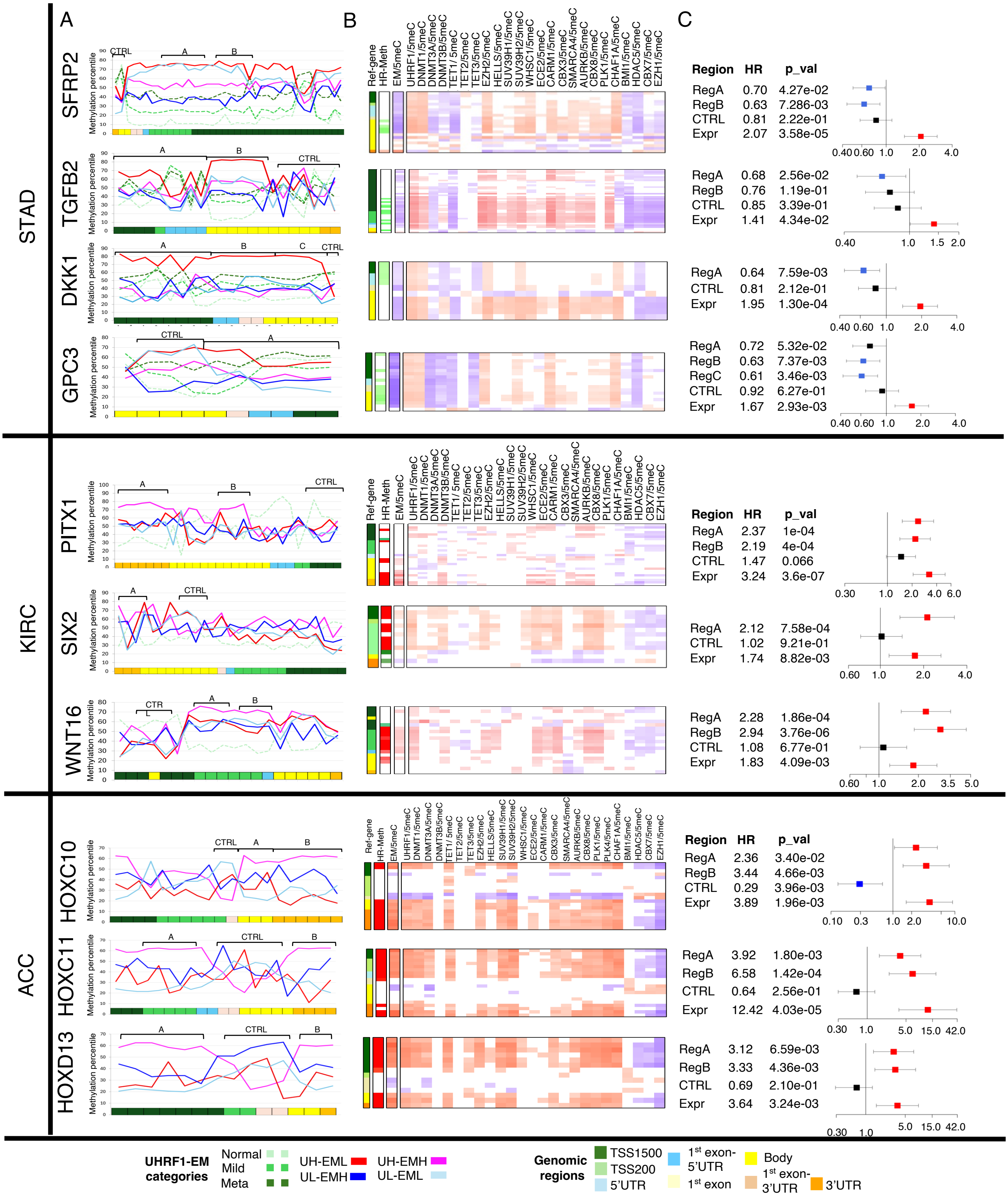


**Figure S15. Differences in DNA methylation and chromatin state in selected EM genes.** From left to right: A) Line-plots: median methylation level of each UHRF1-EM category ; B) heatmap: rows: gene's specific CpGs; columns: annotated Illumina coordinates *(Ref Gene); HR 5meC:* hazard ratio for high vs low methylation level of the CpG (green: HR<1, better prognosis, red: HR>1, worse prognosis); *EM/5meC*: correlation between the EM single gene expression and CpG methylation; *UHRF1/5meC:* correlation between the expression of the indicated epigenetic effector and cytosines' methylation (Spearman's correlation test, blue: anti-correlation; red: correlation; white: no statistically significant) C) Hazard ratios dependent upon high versus low gene expression and for high versus low methylation of the examined genes and CpGs (SFRP2, TGFB2, DKK1, GPC3, PITX1, SIX2, WNT16, HOXC10, HOXC11, HOXD13). Red squares: HR>1, blue squares: HR<1, black squares: HR not significant. Univariate Cox proportional-hazards model was used to calculate hazard ratio (HR) and the significance of the comparison for overall survival. Horizontal lines stand for 95% confidence intervals for the ratios;
